# Drought shifts sorghum root metabolite and microbiome profiles and enriches the stress response factor pipecolic acid

**DOI:** 10.1101/2020.11.08.373399

**Authors:** Daniel F. Caddell, Katherine Louie, Benjamin Bowen, Julie A. Sievert, Joy Hollingsworth, Jeffery Dahlberg, Elizabeth Purdom, Trent Northen, Devin Coleman-Derr

**Affiliations:** Plant Gene Expression Center, USDA-ARS, Albany, CA 94710, USA; Plant and Microbial Biology Department, UC Berkeley, Berkeley, CA 94720, USA; Lawrence Berkeley National Laboratory, Berkeley, CA 94720, USA; Joint Genome Institute, Berkeley, CA 94720, USA; Kearney Agricultural Research & Extension Center, Parlier, CA 93648, USA; Department of Statistics, UC Berkeley, Berkeley, CA 94720, USA

**Author notes:** Corresponding author: D.Coleman-Derr.

**Keywords:** drought, roots, 16S rRNA, amplicon sequencing, microbiome, metabolomics, pipecolic acid, sorghum

## Abstract

Interactions between plants and their root-associated microbiome are important for determining host fitness during periods of stress. During drought, monoderm bacteria are more abundant in sorghum roots than in those of watered controls. Additionally, a reversion from monoderm to diderm dominance occurs in drought-stressed roots one week after rewatering. However, the mechanisms driving this rapid microbiome composition shift is currently unknown. To understand if changes in host metabolism are correlated with this shift, we employed 16S amplicon sequencing and metabolomics of root, rhizosphere, and soil at the peak of a preflowering drought and 24 hours after rewatering. The microbiomes of droughted roots, rhizospheres, and soils differed from watered controls, and shifts in bacterial composition were observed in root and rhizosphere 24 hours after rewatering, highlighting the rapid response of microbes to the cessation of drought. Next, we performed metabolomic profiling to identify putative drivers of this process. During drought, we observed a high abundance of abiotic stress response factors, including antioxidants, osmolytes, amino acids, and plant hormones. After rewatering, large shifts in metabolite abundances were observed in rhizosphere, whereas shifts in root and soil were subtle. In addition, pipecolic acid, a well-characterized systemic acquired resistance signalling compound, was enriched in roots and rhizosphere during drought. We found that exogenous application of pipecolic acid suppresses root growth via a systemic acquired resistance-independent mechanism. Collectively, these data provide a comprehensive characterization of metabolite shifts across three compartments during drought, and elucidate a potential role of pipecolic acid in the sorghum drought response.

**IMPORTANCE:** Plant-associated microbial communities shift in composition and contribute to host fitness during drought. In particular, Actinobacteria are enriched in plant roots and rhizosphere during drought. However, the mechanisms plants use to drive this shift are poorly understood. Here we apply a combination of bacterial and metabolite profiling in root, rhizosphere, and soil during drought and drought-recovery to investigate potential contributions of host metabolism towards shifts in bacterial composition. Our results demonstrate that drought alters metabolic profiles and that the response to rewatering differs between compartments; we identify drought-responsive metabolites that are highly correlated with Actinobacteria abundance. Furthermore, our study reports for the first time that pipecolic acid is a drought-enriched metabolite in sorghum roots. We demonstrate that exogenous application of pipecolic acid is able to provoke one of the classic drought responses in roots, root growth suppression, and that this activity functions independently from the systemic acquired resistance pathway.

## INTRODUCTION

Drought is one of the most significant abiotic stresses impacting agricultural production and threatens to become an even bigger concern due to climate change. Root-associated microbial communities, commonly referred to as microbiomes, influence plant responses to a wide-variety of environmental stresses including drought, and are capable of promoting improved fitness (1). In order to engineer microbiomes capable of enhancing plant growth and ameliorating stress, more research is needed to understand what signals plants use to simultaneously recruit beneficial microbes, while repelling potential pathogens, from a broad pool of soil microbes. Taking advantage of advances in high-throughput sequencing technologies, researchers are beginning to understand the dynamic nature of the microbiome and its association with plant roots. For instance, it is now known that microbiomes vary between environments (2–4), between species (5–7), and even varieties of the same species (4, 8–10). Additionally, microbiomes are dynamic and shift with developmental age, particularly as the plant transitions between vegetative and reproductive growth (11–13). The root microbiome also responds to abiotic stresses. For example, enrichment of monoderm bacteria, such as Actinobacteria, during drought has been observed across diverse plant clades (6, 7, 13, 14). Notably, recent studies have demonstrated that plants mediate the shifts in bacterial communities during drought (15), and Actinobacterial enrichment during drought is dependent on signals produced by living roots (16). While monoderm bacteria are dominant during drought, their enrichment is transitory, with diderm bacteria reestablishing after rewatering (13). Despite their significance, the dynamics regulating monoderm enrichment during drought and the subsequent shift to diderm dominance after rewatering is not yet understood.

Plant-derived metabolites are predicted to drive some of the changes in the root microbiome, as exudation patterns between photoperiods (17) and across development (11, 18, 19) both track with changes in the microbiome. Additionally, a recent study of the maize leaf microbiome across 300 maize genotypes observed associations between specific microbial taxa and host metabolic functions (20). Different combinations of root exudates are sufficient to alter microbiome composition (21), likely by impacting attraction and behavior of soil microbes (22, 23). Some root exudates can act as carbon sources for microbes, while others simultaneously promote certain microbial taxa and suppress others (24). Exudation of putative defense-related metabolites, including organic acids, may also act to repel microbes (23, 25–29). Drought promotes changes in exudate composition, stimulating the exudation of primary and secondary metabolites, including osmoprotectants such as sugars and amino acids. These root and rhizosphere metabolites are predicted to play a role in regulating microbiome associations during drought (30, 31).

Abiotic stresses also contribute to the modulation of plant immunity through modifications to the balance of many hormone pathways, including abscisic acid (ABA), salicylic acid (SA), jasmonic acid (JA), and ethylene (32). For example, ABA, which is strongly induced by drought, antagonizes systemic acquired resistance (SAR) both upstream and downstream of SA (33). In sorghum *(Sorghum bicolor* (L.) Moench), reduced expression of both SA and JA-responsive genes occurs during a prolonged drought (34). This phenomenon has also been observed in response to other abiotic stresses as well; sorghum has reduced SA biosynthesis during nitrogen-limiting conditions (35), and *Arabidopsis* represses immune signalling genes in response to phosphate stress (36). This modulation has major implications for plant-associated microbiomes, as microbes that colonize plant roots must either evade or suppress host immune responses in order to thrive; mutants deficient in immune activation (37) and exogenous application of plant defense hormones JA (38, 39) and SA (40) are both sufficient to alter root microbiomes. Collectively, these studies support the hypothesis that the plant immune system is in part responsible for regulating the establishment of the root microbiome and is impacted by abiotic stress.

In this study, we utilize preflowering drought in field grown sorghum to determine if host metabolism drives the enrichment of monoderm bacteria during drought, and whether drought-driven changes in metabolites may play a role in the rapid transition from monoderm to diderm dominance during re-acclimation to watering. Towards this goal, we employed 16S amplicon sequencing and metabolomic profiling of root, rhizosphere, and soil at the peak of drought and 24 hours after rewatering. We observe an enrichment of monoderm bacteria during drought, consistent with previous reports (6, 7, 13). Furthermore, we determine that the microbiome responds rapidly to rewatering, particularly in the rhizosphere. We also discover that drought alters the metabolite profiles of sorghum root, rhizosphere, and soil. In particular, known drought-associated metabolites such as betaine, 4-aminobutanoic acid (GABA), and amino acids including proline are enriched during drought. Notably, the abundance of a large number of rhizosphere metabolites are rapidly depleted by rewatering following drought, whereas few metabolites shift abundance in the root. In addition to known drought-associated metabolites, we report the detection of pipecolic acid (Pip), a lysine catabolite that is an essential component of SAR (41,42), as a drought-enriched metabolite in sorghum. Here, we demonstrate that exogenous application of Pip suppresses plant root growth, and that this activity functions independently from the established SAR pathway.

## RESULTS

### The sorghum root-associated microbiome is influenced by drought and responds rapidly to rewatering

To examine the effect of a prolonged drought on the root microbiome of sorghum, a field experiment was performed at the University of California’s Agriculture and Natural (UC-ANR) Resources Kearney Agricultural Research and Extension (KARE) Center, in which sorghum plants were subjected to a prolonged preflowering drought, where no water was applied between planting and the onset of flowering (TP8), or regularly irrigated throughout the experiment (figure 1a). We performed 16S rRNA community profiling of root, rhizosphere, and soil using Illumina MiSeq, targeting the V3-V4 variable regions (figure 1b-j). In agreement with a previous study of the sorghum drought microbiome, which was performed at the same location (13), alpha diversity significantly differed between sample types (Shannon, F=82.19, P=1.36×10^-13^), with lower diversity in the root, as compared with rhizosphere and soil, and reduced diversity in droughted roots compared with watered roots (ANOVA, Tukey-HSD, P<0.001) (figure 1c). Beta diversity was assessed through principal coordinates analysis (PCoA) using Bray-Curtis dissimilarities. The primary axis distinguished samples foremost by compartment (root, rhizosphere, or soil), and the second axis by watering regime (droughted or well-watered) (figure 1d), suggesting that both compartment and drought were major driving factors in shaping the microbiome. Pairwise permutational multivariate analysis of variance (PERMANOVA) was performed for each compartment, treatment, and the interaction between compartment and treatment, and all were significantly different (q<0.05) (figure 1d).

**Figure 1.**
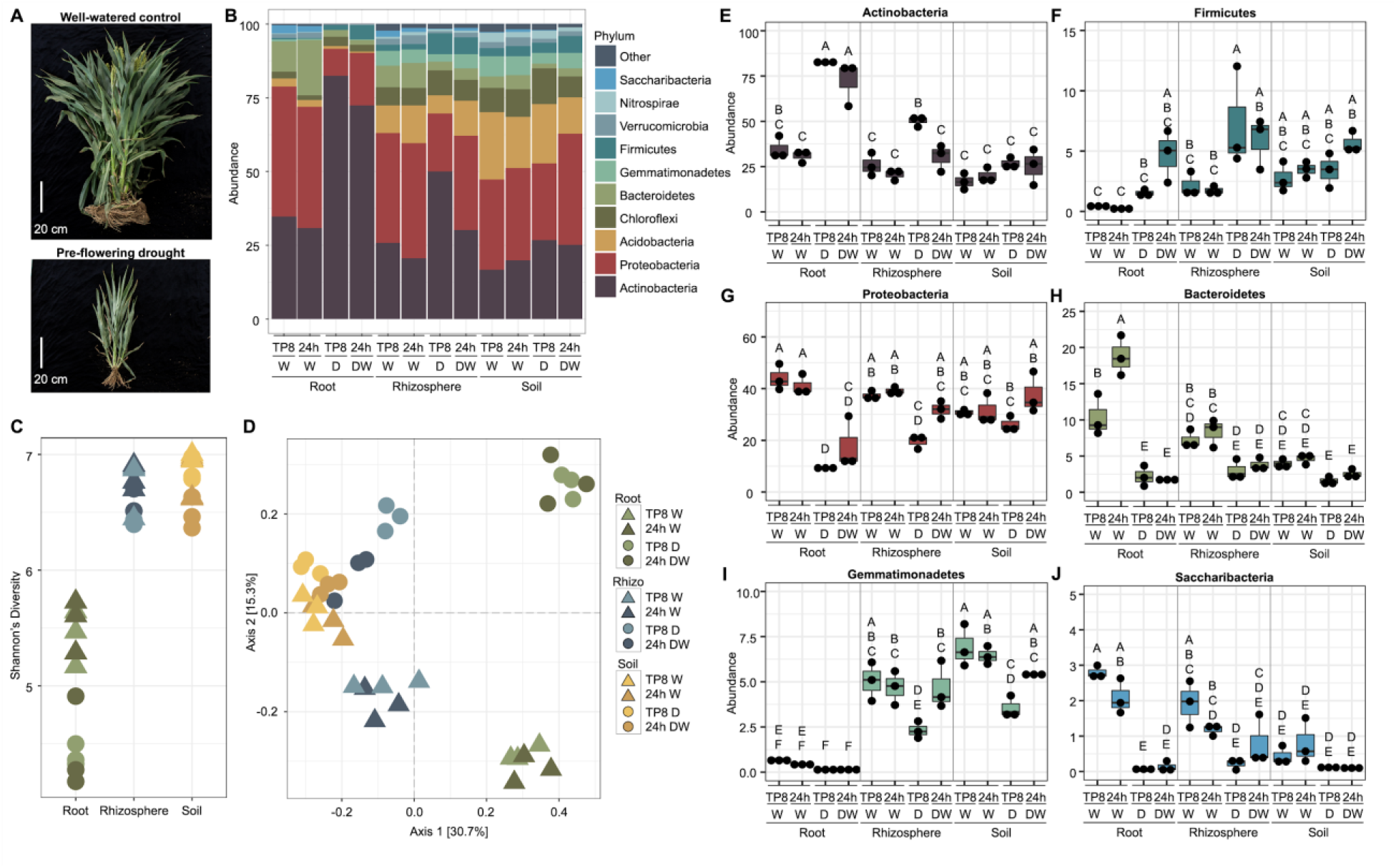
Sorghum root-associated microbiome responds to drought and rewatering. **A** Representative image of sorghum plants following eight weeks of a preflowering drought (TP8). **B** Phylum level relative abundances of sorghum root, rhizosphere, and soil microbiomes at TP8 and 24 hours after rewatering (24h DW) in well-watered (W) or drought (D) plots. **C** Alpha diversity (Shannon) of sorghum root, rhizosphere, and soil. **D** Beta diversity (PCoA) of sorghum root, rhizosphere, and soil microbiomes at TP8 and 24 hours after rewatering in well-watered control or drought plots. **E-J** Relative abundances of individual lineages that displayed a significant difference in abundance between watering treatments *(ANOVA, Tukey-HSD, P<0.05)*.

Similar to previous studies of root microbiomes, we observed a significant enrichment of monoderm bacteria during drought, including taxa belonging to the phylum Actinobacteria in root and rhizosphere, and Firmicutes in the rhizosphere (ANOVA, Tukey-HSD, P<0.05) (figure 1b,e-f). Likewise, diderm lineages were depleted during drought, including Proteobacteria in roots and rhizosphere, Bacteroidetes in roots, and Gemmatimonadetes in the rhizosphere (ANOVA, Tukey-HSD, P<0.05) (figure 1b,g-i). These results suggest that the sorghum root-associated microbiome was responsive to drought, in corroboration with past studies.

Following drought, the sorghum root microbiome responds dramatically to rewatering, with a transition from monoderm back to diderm dominance after a one week recovery period (13). To better understand the early dynamics of this response to rewatering, we watered the droughted sorghum plots after sampling at TP8, and another sampling was performed 24 hours later. Notably, no significant shifts in relative abundance of root phyla were observed (figure 1b). However, a depletion in Actinobacteria and an increase in Gemmatimonadetes occurred in the rhizosphere (ANOVA, Tukey-HSD, P<0.05) (figure 1b,e,i), consistent with monoderm to diderm transitions previously observed after one week of drought recovery (13). Based on levels of beta diversity, the rewatered rhizosphere microbiome appeared more similar to soil, rather than rhizosphere from control samples (figure 1d). These results suggest that the rhizosphere environment provokes a more rapid return to diderm dominance than the root upon rewatering.

### Drought alters the metabolite profiles of sorghum root, rhizosphere, and soil

Recently, substrate utilization was determined to drive microbe community assembly in the rhizosphere across developmental age in another member of the grass lineage, *Avena barbata* (19). To determine if differences in metabolic signals contribute to bacterial community assemblage during drought in field grown sorghum, we performed an untargeted liquid chromatography-mass spectrometry based metabolomic profiling of root, rhizosphere, and soil, using the same samples as bacterial profiling described above. Using a metabolite atlas as reference (43), 112 and 122 polar metabolites were predicted in positive and negative ion modes, respectively, which were then combined to give a total of 168 unique metabolites (supplemental figure 1, supplemental tables 1 and 2). Within these metabolites, we observed different patterns of enrichment across both compartments and treatments, with individual metabolites that were either drought-enriched or drought-depleted (figure 2a). Principal component analysis (PCA) was performed to understand the relationships between samples. PC1 accounted for 59.5% of the total variation and PC2 accounted for 17.1% of the variation, distinguishing samples by both compartment and watering regime (figure 2b). Next, we aimed to determine metabolites that were significantly enriched (*Log_2_ fold change >2, t-test p<0.05)* in each compartment during drought. Twenty-eight, 35, and 16 metabolites were significantly enriched during drought in roots, rhizosphere, and soil, respectively (supplemental table 3), with enrichment that was either compartment specific or observed in multiple compartments (figure 2c).

**Figure 2.**
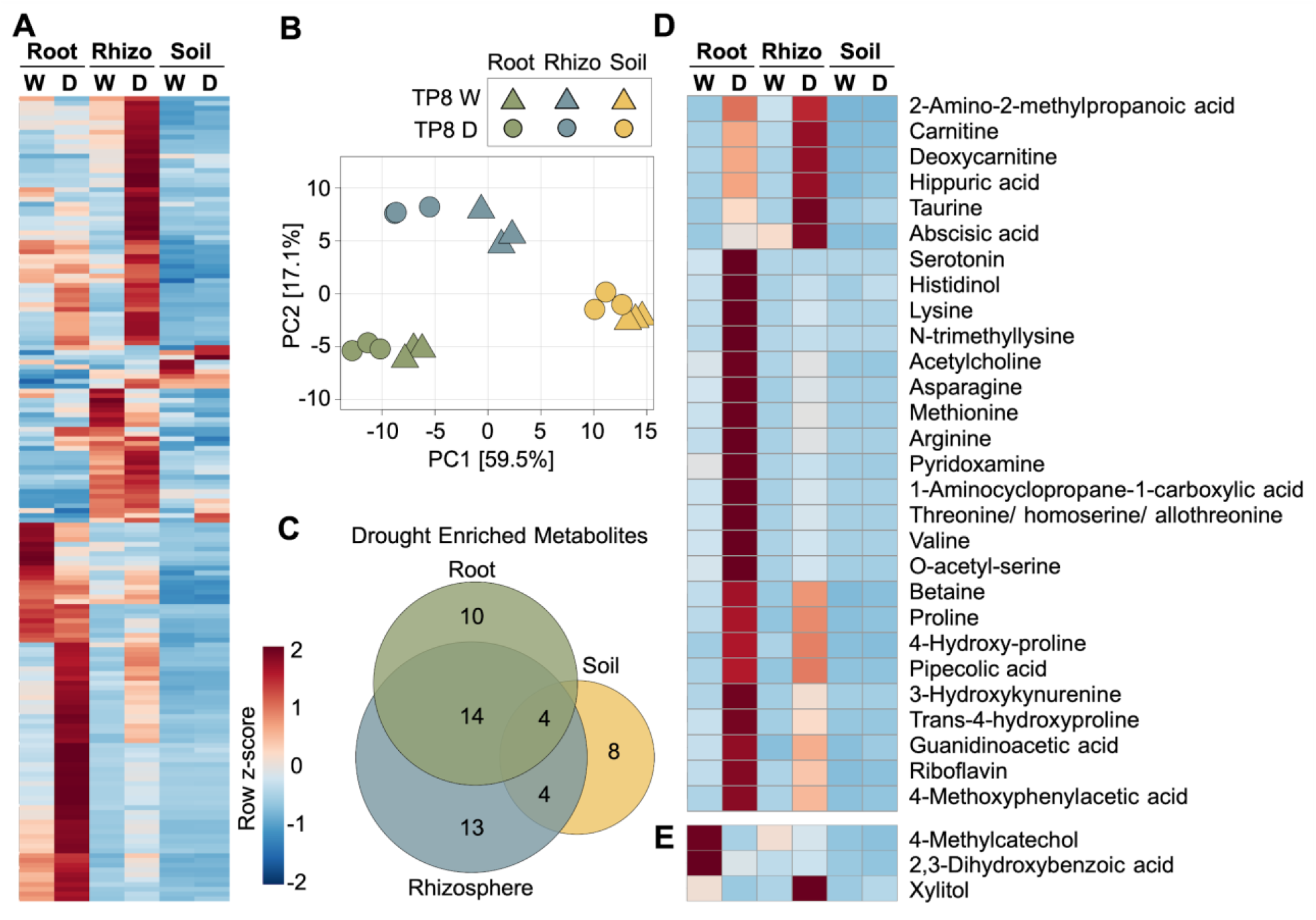
Metabolic profiles during drought differ by compartment. **A** Heatmap of relative peak heights of all observed metabolites (n=168) across root, rhizosphere (rhizo), and soil compartments and watered (W) and drought (D) treatments. **B** Principal component analysis (PCA) plot of root, rhizosphere, and soil metabolites. **C** Proportional Venn diagram of drought enriched metabolites in root, rhizosphere, or soil (D/W Log_2_ fold change >2, t-test p<0.05). **D-E** Heatmap of the subset of metabolites that were enriched or depleted in roots during drought, with the predicted identity of metabolites listed beside each row.

Differences in metabolites observed between compartment and watering treatment suggest that these factors may be responsible for driving associations between plants and their root-associated microbiome. We hypothesized that metabolites with large changes in relative abundance, or fold change, between drought and watered treatments may be important for the observed shifts in the microbiome. In droughted roots, we observed increases in the relative abundance of many putative abiotic stress response factors, including amino acids, osmoprotectants, antioxidants, hormones, and organic acids (figure 2d, table 1). Surprisingly, the important drought markers ABA, 1-aminocyclopropane-1-carboxylic acid (1-ACC, the precursor to ethylene), proline and betaine separated into three distinct clusters of enrichment (figure 2d). In contrast, only 3 metabolites were significantly more abundant in watered roots, including xylitol and the phenolics 2,3-dihydroxybenzoic acid and 4-methylcatechol, which are direct catabolism products of salicylic acid and methylsalicylate, respectively (44, 45) (figure 2e, table 1). Collectively, the observed metabolite enrichment patterns are consistent with sorghum roots responding metabolically to drought.

**Table 1.** Detailed information on significantly changed metabolites between drought and watered roots

| Proposed metabolite | Molecular formula | Measured *m/z* | Measured ppm error | Measured RT peak (minutes) | RT error | MSMS quality score* | Ion mode | Log2 (FC) | p-value (root) | q-value (root) | Significant compartment(s)** |
| --- | --- | --- | --- | --- | --- | --- | --- | --- | --- | --- | --- |
| methionine | C5H11NO2S | 148.044 | 1.748 | 10.6 | 0.1 | 1 | negative | 3.21 | 0.001 | 0.016 | RZS |
| proline | C5H9NO2 | 116.071 | 2.444 | 11.1 | 0.1 | 1 | positive | 2.82 | 0.001 | 0.016 | RZS |
| betaine | C5H12NO2+ | 118.087 | 2.456 | 7.8 | 0.2 | 1 | positive | 2.48 | 0.000 | 0.016 | RZS |
| carnitine | C7H16NO3+ | 162.113 | 0.503 | 13.2 | 0.2 | 1 | positive | 2.43 | 0.012 | 0.038 | RZS |
| N-trimethyllysine | C9H21N2O2+ | 189.160 | 0.178 | 16.8 | 0.1 | 1 | positive | 5.49 | 0.002 | 0.020 | RZ |
| 3-hydroxykynurenine | C10H12N2O4 | 225.087 | 0.497 | 11.2 | 0.1 | 0 | positive | 4.54 | 0.012 | 0.038 | RZ |
| lysine | C6H14N2O2 | 147.113 | 0.569 | 17.0 | 0.1 | 1 | positive | 4.24 | 0.001 | 0.016 | RZ |
| taurine | C2H7NO3S | 126.022 | 1.176 | 12.3 | 0.0 | 0 | positive | 3.94 | 0.012 | 0.038 | RZ |
| arginine | C6H14N4O2 | 175.119 | 0.691 | 17.0 | 0.1 | 1 | positive | 3.81 | 0.002 | 0.018 | RZ |
| pipecolic acid | C6H11NO2 | 130.086 | 0.888 | 11.0 | 0.0 | 1 | positive | 3.39 | 0.001 | 0.016 | RZ |
| allothreonine// homoserine// threonine | C4H9NO3 | 120.066 | 1.912 | 13.7 | 0.1 | 0 | positive | 3.25 | 0.003 | 0.021 | RZ |
| valine | C5H11NO2 | 116.072 | 1.809 | 11.2 | 0.0 | 1 | negative | 3.15 | 0.003 | 0.020 | RZ |
| 1-aminocyclopropane-1-carboxylic acid | C4H7NO2 | 102.055 | 3.955 | 13.7 | 0.6 | 1 | positive | 3.14 | 0.003 | 0.020 | RZ |
| 2-amino-2-methylpropanoic acid | C4H9NO2 | 104.071 | 4.409 | 12.5 | 0.0 | 1 | positive | 2.98 | 0.004 | 0.025 | RZ |
| asparagine | C4H8N2O3 | 133.061 | 0.814 | 14.5 | 0.1 | 1 | positive | 2.91 | 0.006 | 0.027 | RZ |
| cis-4-hydroxy-proline//trans-4-hydroxyproline | C5H9NO3 | 130.051 | 0.823 | 13.4 | 0.3 | 0 | negative | 2.90 | 0.004 | 0.022 | RZ |
| guanidinoacetic acid | C3H7N3O2 | 118.061 | 2.161 | 14.0 | 0.1 | 0 | positive | 2.43 | 0.003 | 0.020 | RZ |
| deoxycarnitine | C7H16NO2+ | 146.118 | 0.662 | 13.3 | 0.2 | 1 | positive | 2.39 | 0.024 | 0.061 | RZ |
| serotonin | C10H12N2O | 177.102 | 0.870 | 8.7 | 0.1 | 0 | positive | 3.47 | 0.042 | 0.087 | RO |
| histidinol | C6H11N3O | 142.098 | 0.107 | 12.9 | 0.2 | 0 | positive | 2.95 | 0.014 | 0.044 | RO |
| trans-4-hydroxyproline | C5H9NO3 | 132.066 | 0.768 | 13.4 | 0.1 | 0 | positive | 2.83 | 0.004 | 0.022 | RO |
| o-acetyl-serine | C5H9NO4 | 148.060 | 0.072 | 11.4 | 0.1 | -1 | positive | 2.83 | 0.016 | 0.045 | RO |
| 4-methoxyphenylacetic acid | C9H10O3 | 184.097 | 0.912 | 1.1 | 0.0 | 0 | positive | 2.79 | 0.005 | 0.026 | RO |
| riboflavin | C17H20N4O6 | 377.146 | 1.147 | 4.6 | 0.0 | 0 | positive | 2.67 | 0.004 | 0.025 | RO |
| hippuric acid | C9H9NO3 | 180.066 | 0.924 | 4.6 | 0.3 | 0 | positive | 2.41 | 0.009 | 0.034 | RO |
| acetylcholine | C7H16NO2+ | 146.118 | 0.496 | 2.1 | 0.1 | 1 | positive | 2.38 | 0.010 | 0.036 | RO |
| pyridoxamine | C8H12N2O2 | 169.097 | 0.267 | 10.2 | 0.0 | 0 | positive | 2.12 | 0.001 | 0.016 | RO |
| abscisic acid | C15H20O4 | 247.133 | 1.332 | 1.1 | 0.0 | 0 | positive | 2.05 | 0.006 | 0.026 | RO |
| 2,3-dihydroxybenzoic acid// 2,5-dihydroxybenzoic acid// 3,4-dihydroxybenzoic acid | C7H6O4 | 153.020 | 1.467 | 4.0 | 0.1 | 0 | negative | -2.11 | 0.000 | 0.012 | RO |
| xylitol | C5H12O5 | 175.058 | 4.087 | 5.1 | 0.0 | 0 | positive | -3.16 | 0.001 | 0.016 | RO |
| 4-methylcatechol | C7H8O2 | 123.045 | 1.158 | 4.3 | 0.4 | 0 | negative | -3.73 | 0.001 | 0.016 | RO |

\*MSMS quality scores: 1 (MSMS matches ref. std.); 0.5 (possible match); 0 (no MSMS collected or no appropriate ref available); -1 (MSMS poor match to ref. std.)
\*\*Compartment(s) with significant Log2 (fold change): Root, rhizosphere, and soil (RZS); root and rhizosphere (RZ); root only (RO)

### Response to rewatering within 24 hours following a prolonged drought varies by compartment

Having established that metabolite composition differs between watered and drought sorghum roots, we next sought to understand whether rapid shifts in metabolite profiles would be observed 24 hours after rewatering the droughted sorghum plots. We hypothesized that rewatering would shift metabolite compositions, particularly in the rhizosphere, that could contribute to the transitions observed in microbial community composition. Consistent with this hypothesis, we observed distinct response patterns in metabolites across all compartments 24 hours after rewatering the droughted plots. Notably, roots were only weakly responsive to rewatering, with no metabolites strongly enriched in rewatered roots (Log_2_ fold change >2, t-test p<0.05) (figure 3a). However, several metabolites were modestly enriched (Log_2_ fold change >1, t-test p<0.05), including cytosine, sphinganine, N-acetylglutamic acid, which promotes growth of root hairs and swelling of root tips (46), and ferulic acid, which is capable of inhibiting root growth and promotes root branching (47) (supplemental table 4). Collectively, these shifts suggest roots have started to respond to changes in water availability, although their overall metabolite profiles are largely unchanged at the time of sampling.

**Figure 3.**
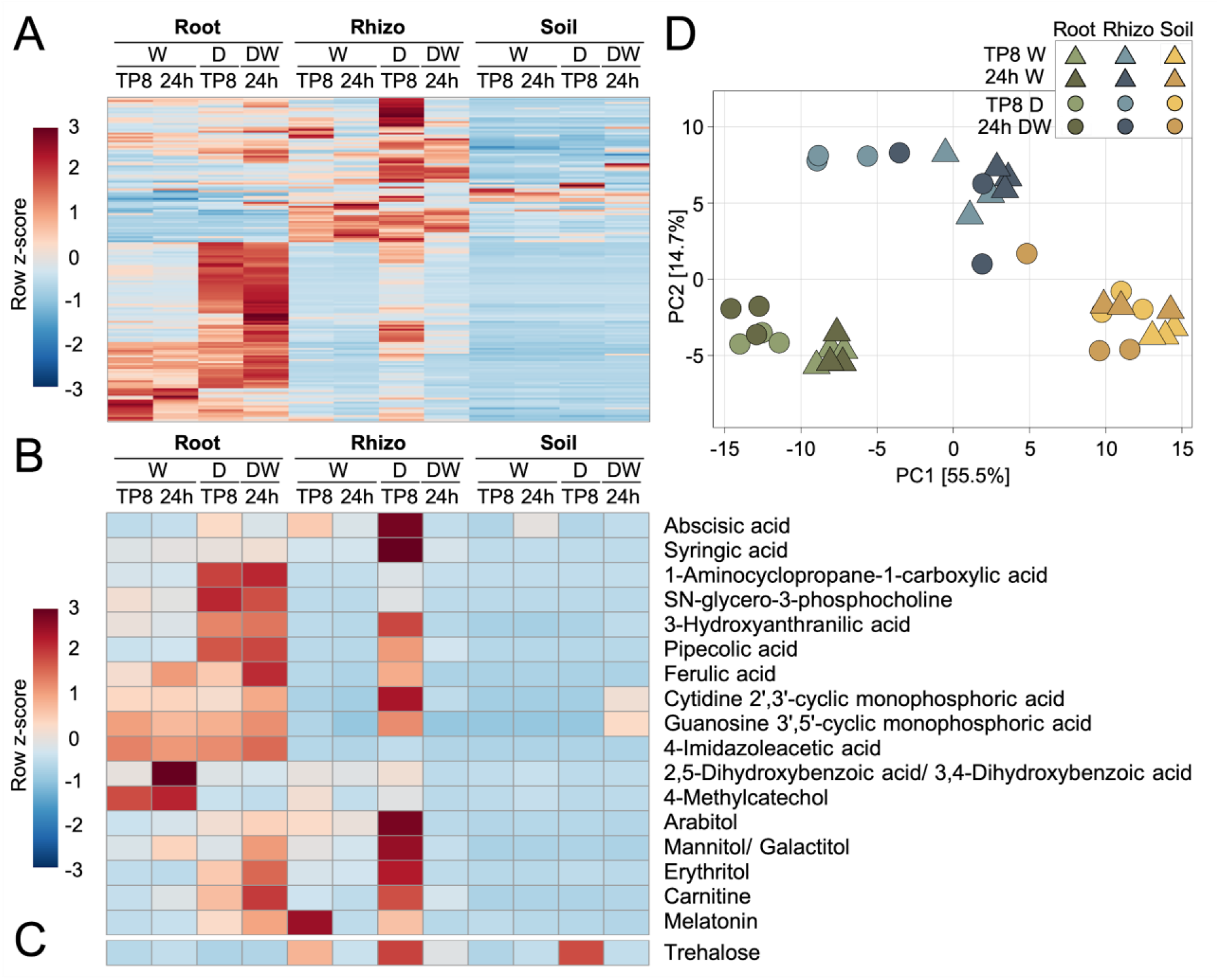
Rewatering depletes rhizosphere metabolites following a prolonged drought. **A** Heatmap of relative peak heights of all observed metabolites (n=168) across three compartments (root, rhizosphere (rhizo), and soil), three treatments (watered (W), drought (D), drought rewatered (DW), and two time points (time point 8 (TP8) and 24 hours later (24h)). **B-C** Heatmap of the subset of metabolites that were depleted after rewatering (DW/D Log_2_ fold change < −2, t-test p<0.05), with the predicted identity of metabolites listed beside each row. Note, all significant depletions were observed in the rhizosphere, except trehalose, which occurred in soil. **D** Principal component analysis (PCA) plot of root, rhizosphere, and soil metabolites.

In contrast to roots, large shifts in metabolite abundances occurred in the rhizosphere after rewatering, with rewatering tending to cause a depletion of rhizosphere metabolites (figure 3a). Significantly depleted metabolites (n=17, Log_2_ fold change < −2, t-test p<0.05) included ten different organic acids, four sugar alcohols, sn-glycero-3-phosphocholine, carnitine, and melatonin (figure 3b, table 2). In soil, only a single metabolite, trehalose, was significantly depleted following rewatering (figure 3c, table 2). Notably, the metabolite composition of rewatered rhizosphere became more similar to watered rhizosphere, and was distinguishable from soil (figure 3d).

**Table 2.** Detailed information on significantly changed metabolites 24 hours after rewatering

| Proposed metabolite | Molecular formula | Measured *m/z* | Measured ppm error | Measured RT peak (minutes) | RT error | MSMS quality score* | Ion mode | Log2 (FC) | p-value (root) | q-value (root) | Significant compartment |
| --- | --- | --- | --- | --- | --- | --- | --- | --- | --- | --- | --- |
| 4-imidazoleacetic acid | C5H6N2O2 | 127.050 | 0.936 | 13.9 | 0.2 | 1 | positive | -2.44 | 0.049 | 0.296 | rhizosphere |
| 1-aminocyclopropane-1-carboxylic acid | C4H7NO2 | 102.055 | 3.955 | 13.7 | 0.6 | 1 | positive | -2.44 | 0.047 | 0.296 | rhizosphere |
| pipecolic acid | C6H11NO2 | 130.086 | 0.888 | 11.0 | 0.0 | 1 | positive | -2.45 | 0.048 | 0.296 | rhizosphere |
| carnitine | C7H16NO3+ | 162.113 | 0.503 | 13.2 | 0.2 | 1 | positive | -2.49 | 0.017 | 0.231 | rhizosphere |
| melatonin | C13H16N2O2 | 233.128 | 1.296 | 1.2 | 0.0 | 0 | positive | -2.89 | 0.011 | 0.213 | rhizosphere |
| 3-hydroxyanthranilic acid | C7H7NO3 | 152.036 | 1.229 | 1.8 | 0.1 | 0 | negative | -3.11 | 0.003 | 0.135 | rhizosphere |
| syringic acid | C9H10O5 | 197.046 | 0.404 | 1.6 | 0.1 | 1 | negative | -3.42 | 0.040 | 0.296 | rhizosphere |
| cytidine 2',3'-cyclic mono-phosphoric acid | C9H12N3O7P | 306.049 | 1.863 | 14.0 | 0.2 | 0 | positive | -3.44 | 0.047 | 0.296 | rhizosphere |
| abscisic acid | C15H20O4 | 247.133 | 1.332 | 1.1 | 0.0 | 0 | positive | -3.56 | 0.002 | 0.135 | rhizosphere |
| galactitol/mannitol | C6H14O6 | 181.072 | 1.680 | 9.7 | 0.1 | 1 | negative | -3.58 | 0.026 | 0.263 | rhizosphere |
| 4-methylcatechol | C7H8O2 | 123.045 | 1.158 | 4.3 | 0.4 | 0 | negative | -4.45 | 0.005 | 0.169 | rhizosphere |
| 2,3-dihydroxybenzoic acid//<br>2,5-dihydroxybenzoic acid//<br>3,4-dihydroxybenzoic acid | C7H6O4 | 153.020 | 1.467 | 4.0 | 0.1 | 0 | negative | -4.45 | 0.040 | 0.296 | rhizosphere |
| erythritol | C4H10O4 | 121.051 | 0.626 | 3.2 | 0.0 | 1 | negative | -5.04 | 0.024 | 0.252 | rhizosphere |
| arabitol | C5H12O5 | 151.061 | 1.358 | 5.6 | 0.2 | 1 | negative | -5.29 | 0.003 | 0.135 | rhizosphere |
| sn-glycero-3-phosphocholine | C8H21NO6P+ | 258.110 | 0.111 | 14.8 | 0.2 | 1 | positive | -5.66 | 0.017 | 0.231 | rhizosphere |
| guanosine 3',5'-cyclic monophosphoric acid | C10H12N5O7P | 346.055 | 0.947 | 13.9 | 0.2 | 0 | positive | -5.73 | 0.000 | 0.019 | rhizosphere |
| ferulic acid | C10H10O4 | 193.051 | 0.614 | 1.3 | 0.1 | 1 | negative | -5.75 | 0.008 | 0.181 | rhizosphere |
| lactose/trehalose | C12H22O11 | 401.131 | 1.143 | 14.4 | 0.1 | 1 | negative | -3.07 | 0.002 | 0.340 | soil |

\*MSMS quality scores: 1 (MSMS matches ref. std.); 0.5 (possible match); 0 (no MSMS collected or no appropriate ref available); -1 (MSMS poor match to ref. std.)

### Pipecolic acid suppresses plant root growth

Betaine represents one of the most robust and widely utilized biomarkers of plant responses to drought (31, 48). We hypothesized that other metabolites with roles in plant drought response would share a similar abundance pattern during drought. To identify other putative drought metabolites, we ranked metabolites based on their correlation coefficients (Pearson’s r) with betaine, across all compartments, treatments, and timepoints (figure 4a-c). The most strongly correlated metabolite, pipecolic acid (Pip), is a lysine catabolite that has recently been identified as a critical component of the systemic acquired resistance (SAR) pathway (41, 42). However, to our knowledge no link between Pip and drought stress response has been demonstrated in plants, although its synthesis is osmotically induced in rapeseed leaf discs (49) and in the halophyte *Triglochin maritima* (50). Beyond Pip, the other nine of the top 10 correlated metabolites have all been previously identified as drought-related metabolites. These include 4-aminobutanoic acid (GABA) (51), allantoin (52, 53), carnitine (54, 55), and the amino acids proline, tyrosine, asparagine, serine, glutamine, and trans-4-hydroxyproline (56–60) (figure 4a).

**Figure 4.**
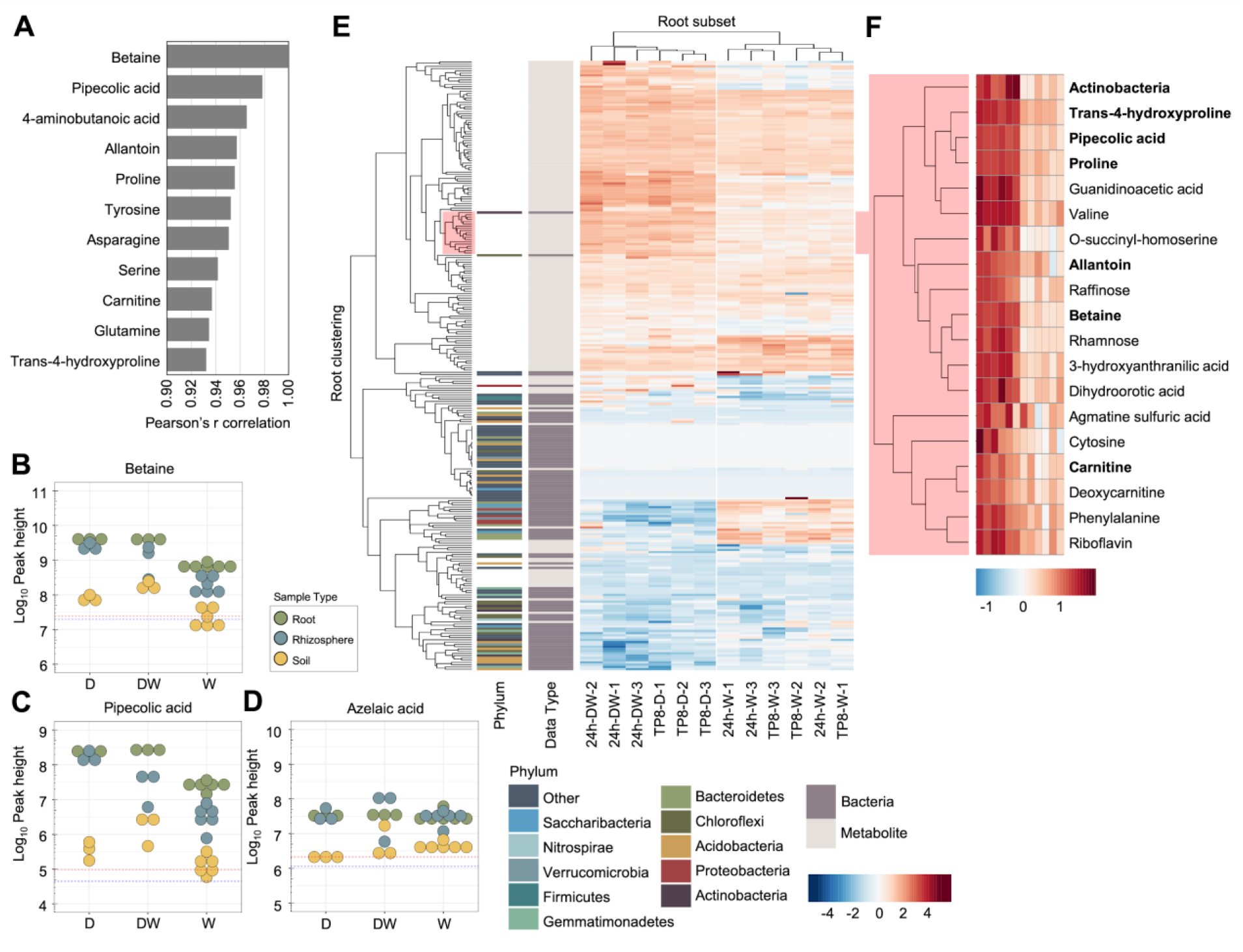
Pipecolic acid abundance pattern mirrors drought markers. **A** The top 10 metabolites correlated with the drought marker betaine across all sample types, treatments, and time points. **B-D** Log_10_ peak heights of individual metabolites. Each point represents an individual sample of root (green), rhizosphere (blue), or soil (yellow). Dashed lines represent the limit of detection for individual metabolites, based on the average log_10_ peak heights of the sample blanks for root (red) or rhizosphere and soil (blue). **E** Heatmap of relative abundance of all metabolites and bacteria ASVs (grouped at the class level), clustered within the root, across treatments (watered (W), drought (D), drought rewatered (DW), and time points (time point 8 (TP8) and 24 hours later (24h)). **F** Zoom-in of Actinobacteria and closely clustering root metabolites, as highlighted in pink in figure 4e. Actinobacteria and the metabolites that are closely correlated with betaine (as in figure 4a) are in bold.

To identify potential interactions between these drought-related metabolites and the root-associated bacterial community, we clustered bacterial ASVs (grouped at the class level) and metabolites based on abundances across all compartments, treatments, and timepoints. Strikingly, while a majority of metabolites and microbes separated into distinct clusters, three microbial taxa, including the Actinobacteria, nested within the metabolite-dominant cluster, just adjacent to the metabolite cluster containing the top ten betaine-correlated metabolites (supplemental figure 2). When clustering was performed based on the abundances in the root, where host control of the microbiome is strongest, we observed that the Actinobacteria formed an even closer linkage with betaine-correlated metabolites, and was tightly clustered with five metabolites including trans-4-hydroxyproline, proline, Pip, valine, and guanidinoacetic acid (figure 4e-f, supplemental figure 3). As previous reports have demonstrated that drought-induced microbial lineages including Actinobacteria in roots are capable of inducing SAR (61) and systemic root-to-root signaling (62), we hypothesized that the increased abundance of Actinobacteria in sorghum roots may contribute to the accumulation of Pip during drought, and potentially the activation of systemic signaling. However, azelaic acid, which functions downstream of Pip in the SAR pathway (63), was not enriched by drought in our data, suggesting that SAR was not being activated (figure 4d). Collectively, these results suggest that Pip may play an as-yet undiscovered role in sorghum drought response.

One classic and easily observable phenotypic shift that occurs in roots during drought is a suppression of root growth (64, 65). We hypothesized that if Pip plays an integral role in the drought response pathway, application of Pip should lead to reduced root growth. To evaluate this possibility, we germinated sorghum in petri dishes containing water plus 0, 0.1, or 1 mM Pip. Seven days after germination, 1 mM Pip treated sorghum displayed significantly reduced root growth (figure 5a-b). Exogenous Pip application is also capable of reducing root growth in *Arabidopsis* (66). We confirmed this result by growing *Arabidopsis* Col-0 on petri dishes containing either 0, 0.001, 0.01, 0.1, 1 mM Pip. Average root growth was reduced in all Pip treatments in a dosage dependent manner, with significant decreases in root growth observed with 0.1 and 1 mM Pip concentrations (figure 5c-d).

**Figure 5.**
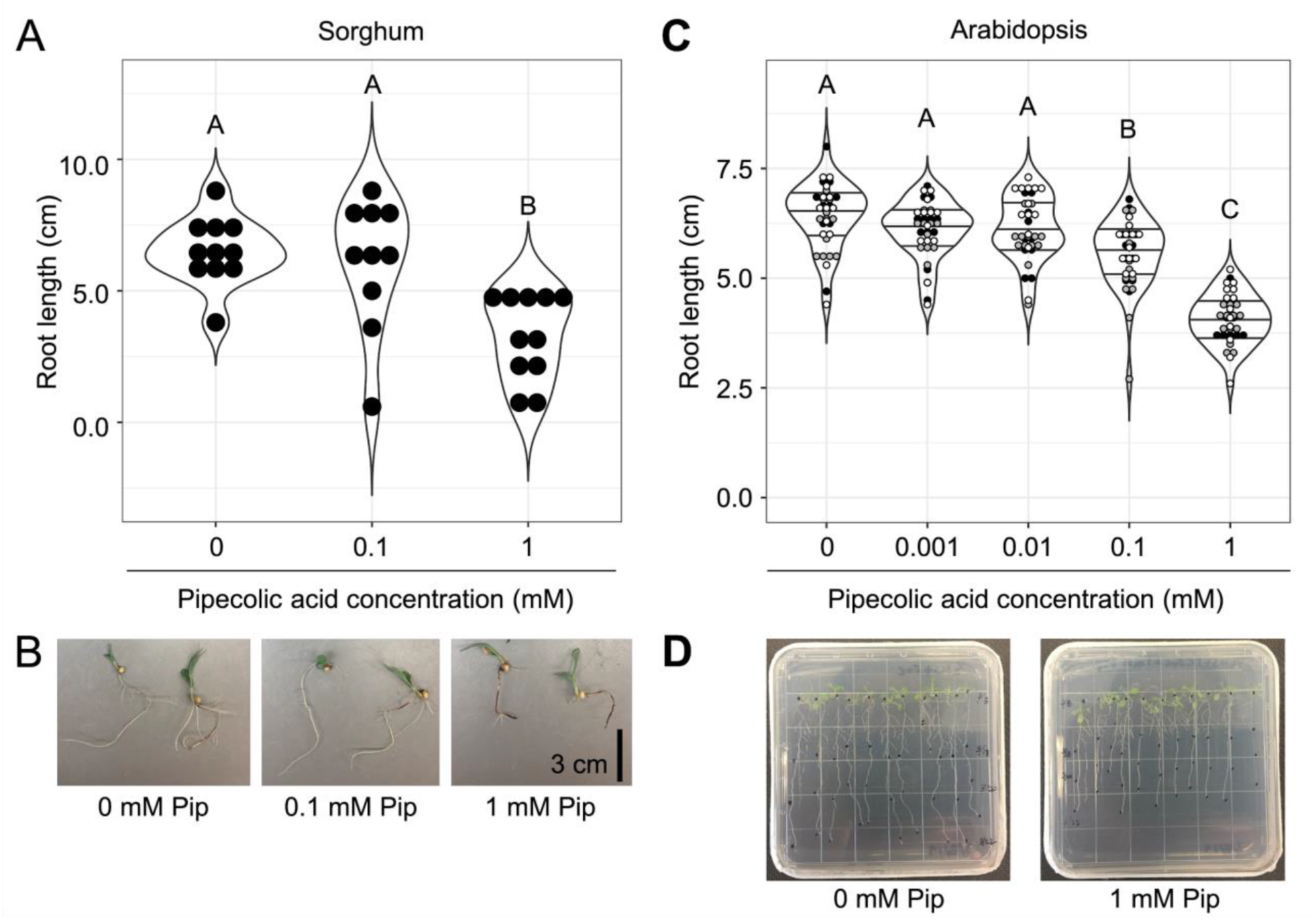
Pipecolic acid reduces root growth. **A** Root lengths of sterilized sorghum seedlings after 7 days of growth in water containing 0, 0.1, or 1 mM pipecolic acid (Pip). Different letters indicate a significant difference in root length (ANOVA, Tukey-HSD, p<0.05). This experiment was performed twice with similar results. **B** Two representative seedlings from each treatment were photographed at the time of measurement. **C** Root lengths of sterilized Arabidopsis seedlings after 10 days of growth in ½ MS+ 1% sucrose agar media containing 0, 0.001, 0.01, 0.1, or 1 mM Pip. Different letters indicate a significant difference in root length (ANOVA, Tukey-HSD, p<0.05). Different colors represent plants from independent experiments (n=3). **D** One representative plate from 0 and 1 mM pipecolic acid treatments were photographed at the time of measurement.

Having confirmed that exogenous Pip is sufficient to reduce root growth, we next aimed to understand the molecular mechanism responsible for this behavior. As Pip plays a key role in systemic signalling during SAR (67), we hypothesized that Pip-mediated root growth reduction may depend on components of this signalling pathway. To test this hypothesis, we utilized publicly available genetic resources in *Arabidopsis,* including *fmo-1, npr1-1, rbohd/rbohf, azi1-2* mutants, which represent critical aspects of the SAR signalling pathway (figure 6). We measured the root growth of each of these validated *Arabidopsis* SAR mutants on media containing 1 mM Pip. Notably, mutants in FLAVIN-CONTAINING MONOOXYGENASE 1 (FMO-1), responsible for conversion of Pip to N-hydroxy-Pip (68)(figure 6), displayed reduced root growth similar to the wild-type plant Col-0 in response to Pip (figure 6), indicating that this conversion is not required to elicit reduced root growth. Likewise, mutants in NON EXPRESSER OF PATHOGENESIS RELATED GENES 1 (NPR1) (69), a SA receptor required for SA-dependent SAR, and mutants in NADPH/RESPIRATORY BURST OXIDASE PROTEINS D and F (RBOH D/F) and the lipid transfer protein AZELAIC ACID INDUCED 1 (AZI1) (70, 71), required for SA-independent SAR, also responded similar to Col-0 (figure 6).Collectively, these data indicate that root growth response to Pip is likely not mediated by the SAR pathway.

**Figure 6.**
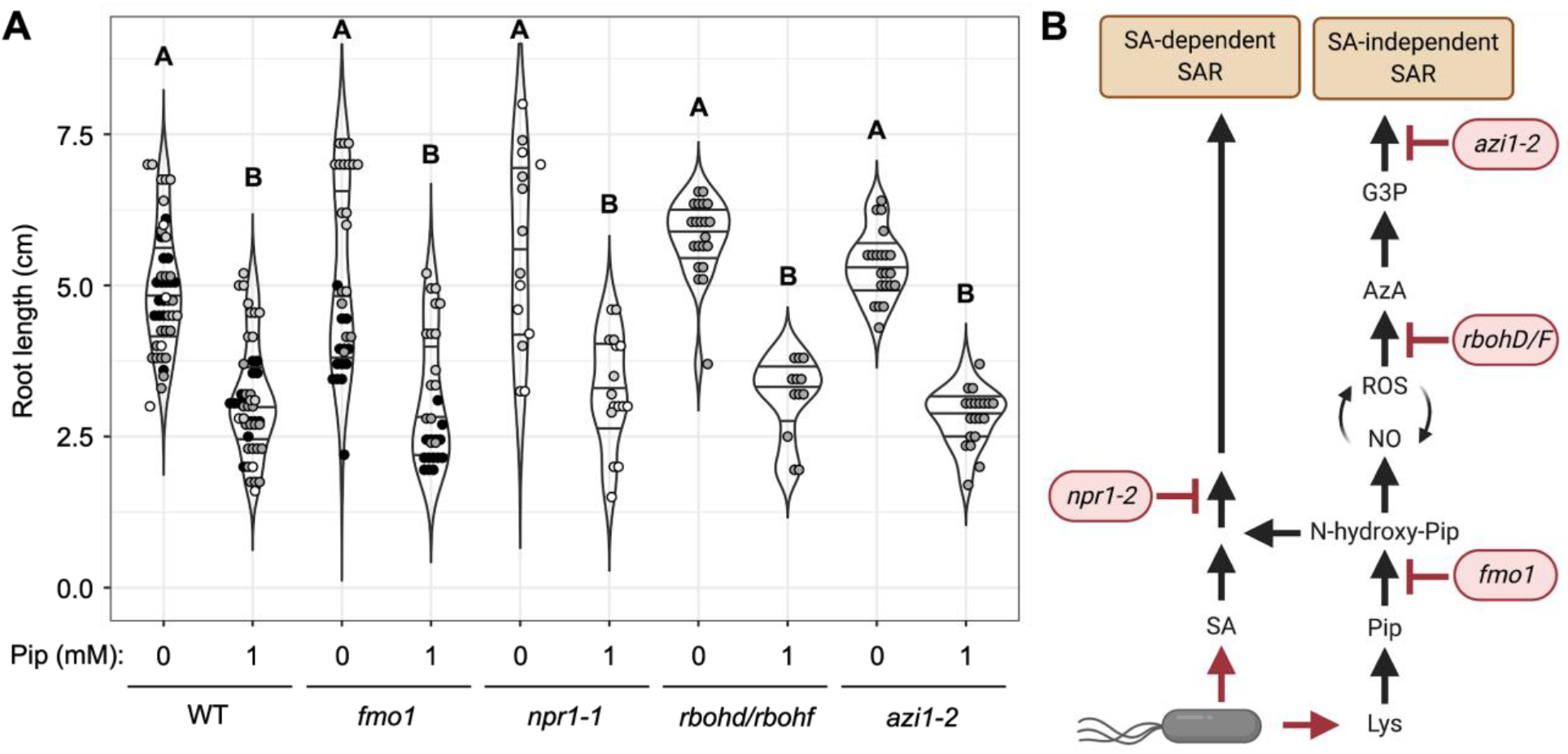
Pipecolic acid root growth reduction is SAR-independent. **A** Root length of Arabidopsis Col-0 (WT) and Arabidopsis mutants grown on 1/2MS + 1% sucrose plates containing 0 or 1 mM Pip. Significance between treatments was evaluated by ANOVA with Tukey’s HSD posthoc test (p<0.05). Different colors represent plants from independent experiments. **B** Simplified SAR pathway. Highlighted in red are the Arabidopsis mutants used to evaluate a potential interaction between SAR and Pip-mediated root growth suppression.

## DISCUSSION

### Metabolite and microbial community compositions shift during drought and rewatering

The assemblage of the microbiomes differ between root, rhizosphere, and soil during drought, though the underlying cause of these differences is not well understood. Plant metabolites and exudates have been hypothesized to drive these changes (30), and plants are known to increase total organic carbon and root exudation (per gram of root biomass) in response to drought (72, 73). The enrichment of specific metabolites that can act as osmoprotectants, such as betaine, sugars, and amino acids, is frequently reported when plants are subjected to drought, and these are commonly used as drought-specific markers. Yet, comprehensive studies of the global metabolite profiles across root, rhizosphere, and soil have been hindered by the complexity of soil metabolite profiles. However, recent advances in metabolomics have allowed for characterization of metabolite profiles within and across complex substrates (74). As a result, we collected both microbiome and metabolite profiles across three different compartments (sorghum root, rhizosphere, and soil) and three treatments (watered, drought, and drought recovery). We observed a general trend that many metabolites were more abundant during drought. This observation is in line with previous evidence that exudation by plants increases during drought (72). Interestingly, we observed both shifts in metabolite abundances during drought that were compartment specific and shifts that occurred across multiple compartments. Likewise, we observed enrichment of specific drought responsive metabolites that correlated with increases in both monoderm dominance during drought and compartment specific shifts upon rewatering. Our findings that changes in host metabolism during drought are correlated with shifts in certain bacterial taxa expand upon a recent finding that substrate utilization also drives microbial community assembly in the rhizosphere across developmental age (19).

A previous study of sorghum root and rhizosphere showed that bacterial community abundance reverts from monoderm dominance during drought to diderm dominance within a week following rewatering (13). To understand the early dynamics of this reversion, we evaluated sorghum root, rhizosphere, and soil at the peak of a preflowering drought and again 24 hours after rewatering. We observed distinct response patterns in metabolites of root, rhizosphere, and soil 24 hours after rewatering, which could impact the establishment of microbiomes following rewatering. Notably, the rhizosphere was the most responsive in both metabolites and microbiome profiles within 24 hours after rewatering. We propose that the more rapid shifts observed in rhizosphere are largely driven by flow of water, which may simultaneously dilute rhizosphere metabolites into the surrounding soil, and promote the mobility of soil microbes to enter the rhizosphere. Supporting this hypothesis, the beta diversity of rewatered rhizosphere microbiome early after rewatering more closely resembled the soil microbiome, rather than watered rhizosphere samples, suggesting that the soil microbiome is a source of inoculum for bacterial establishment in the rhizosphere following rewatering. In contrast, rhizosphere metabolites were characterized by depletion after rewatering, with similarity to watered rhizosphere, which may facilitate the eventual reversion to a watered rhizosphere microbiome, as demonstrated previously (13). Comparable studies on the long-term effects of drought and subsequent rewatering on metabolite profiles in grasses are lacking, however previous studies of trees have reported mixed long-term effects. One study found that increases in exudation can be reversed following recovery from drought, to be indistinguishable from controls (73), while another found that extreme drought led to irreversible changes in exudation (59). Therefore, future studies will benefit from careful selection of multiple time points following drought to understand the long-term effects of drought in roots and rhizosphere following rewatering.

### Pipecolic acid-mediated root growth suppression is not mediated by the systemic acquired resistance pathway

In this study, we observed an enrichment of many putative abiotic stress response factors during drought. Of particular note, Pip was significantly enriched during drought, and its abundance correlated strongly with other highly enriched drought markers including Actinobacteria and the metabolites betaine, proline, and GABA. While Pip induction in response to osmotic stress has been noted in a few previous studies (49, 50), by far Pip’s most notable role is in systemic signalling of stress in response to pathogens. It has recently been shown that the conversion of Pip to N-hydroxy-Pip is required for SAR to be activated, and this activity functions both alongside and independent of SA (67, 75). Notably, previous research has demonstrated that “biotic stress” factors, including SA, can benefit plants responding to abiotic stress. For example, plant drought tolerance can be promoted by increasing endogenous SA in *Arabidopsis* (76), and exogenous SA also promoted drought tolerance via an NPR1-dependent mechanism in *Brassica napus* (77). However, recent studies suggest that SA signaling is inactivated by drought, in part because ABA promotes NPR1 degradation (33, 78). Additionally, a strong suppression of defense-related gene expression occurs in field grown sorghum during preflowering drought (34). Likewise, we did not see an enrichment of azelaic acid, which functions downstream of Pip during SAR, during drought in our data. These findings suggest that the Pip induction observed during drought is likely acting for some other purpose, perhaps as an independent signalling mechanism related to abiotic stress.

Here, we demonstrate that Pip application is able to provoke one of the classic drought responses in roots, namely root growth suppression (64, 65), which suggests it could be involved in the drought response pathway. As exogenous Pip was capable of suppressing root growth in both sorghum and *Arabidopsis,* this suggests that a common mechanism is conserved across plants. We hypothesized that Pip-mediated root growth suppression might share components with the SAR pathway, which functions across diverse plant clades (79). However, using previously validated *Arabidopsis* SAR pathway mutants, we demonstrate that this Pip activity in the root functions independently from the established SAR pathway, which has primarily been evaluated in leaves. These data suggest that although Pip has been primarily characterized as a component of SAR in plants, it may also act as a more general stress response factor to environmental shifts including drought using an alternative mechanism. Furthermore, azelaic acid, another component of SAR that functions downstream of Pip, is not enriched in sorghum roots under drought, which suggests that this pathway, if it exists, acts differently than the one currently known example.

Interestingly, Pip, which was significantly enriched in both roots and rhizosphere, has also been shown to have a direct role on microbes, where it is predicted to function in osmoprotection. For example, Pip has been widely demonstrated to improve the growth of a diverse bacteria challenged by NaCl-induced osmotic stress, including the Actinobacteria lineage *Brevibacterium ammonkgenes* (80), and Proteobacteria lineages *Escherichia coli* (81), *Sinorhizobium meliloti* (82), and *Silicibacter pomeroyi* (83). Supporting its role as an osmoprotectant, exogenous application of another osmoprotectant, betaine, suppressed the salt-induced accumulation of Pip (80). These studies support a possible role of rhizosphere Pip acting as an osmolyte to protect bacteria during drought. However, as Pip appears to be functional across a broad range of bacterial lineages, we do not believe Pip is likely to be responsible for the shifts in monoderm and diderm dominance observed during drought and subsequent rewatering. Collectively, our results highlight the need for future studies that delve further into the potential contribution(s) of Pip to plant drought responses.

## METHODS

### Field experimental design and sample collection

Sorghum cultivar RTx430 plants were grown in the summer of 2017, in a field located at the UC-ANR KARE Center located in Parlier, California (36.6008 N 119.5109 W), as described previously (13, 84). Sorghum seeds were sown into pre-watered fields. Starting in the third week, control treatment plants were watered 1h three times per week by drip irrigation (1.89 L h-1 flow rate), and no water was provided to drought treatment plants. After eight weeks, which coincided with the onset of flowering, roots and rhizosphere samples were harvested prior to watering (TP8). Water was then restored to the drought plots (rewatered), and root and rhizosphere samples were harvested after 24 h (TP8+24h). All field samples were collected between 11am and 12pm using a modified version of the protocol described in detail in (85). Soil samples were collected using a 15 cm soil core sampler, at a distance of approximately 20 cm from the base of the plant. To collect rhizosphere compatible with both microbiome and metabolomic analyses, excavated plants were briefly shaken to dislodge excess soil, and an ethanol-sterilized nylon bristled toothbrush was used to remove closely adhering soil from the root, which we collected as the rhizosphere fraction, prior to vortexing the roots two times for 1 min in epiphyte removal buffer (ice cold 0.75% KH_2_PO_4_, 0.95% K_2_HPO_4_, 1% Triton X-100 in ddH_2_O; filter sterilized at 0.2 μM). Any remaining soil adhering to the root was separated with epiphyte removal buffer and discarded. The roots were again rinsed with clean epiphyte removal buffer and patted dry. All samples were immediately flash frozen in LN_2_ in the field and stored at −80 °C until sample processing.

### DNA extraction, amplification, and amplicon sequencing

DNA extraction was performed using the protocol for collection of root endosphere, rhizosphere, and soil samples using Qiagen DNeasy Powersoil DNA extraction Kit with 0.15 g (root) and 0.25 g (rhizosphere and soil) as starting material in the provided Powersoil collection vials, as described in detail in (85). The V3-V4 region of the 16S rRNA gene was PCR amplified from 25 ng of genomic DNA using dual-indexed 16S rRNA Illumina iTags 341F (5’-CCTACGGGNBGCASCAG-3’) and 785R (5’-GACTACNVGGGTATCTAATCC-3’). Barcoded 16S rRNA amplicons were quantified using Qubit dsDNA HS assay kit on a Qubit 3.0 fluorometer (Invitrogen, Carlsbad, CA, USA), pooled in equimolar concentrations, purified using Agencourt AMPure XP magnetic beads (Beckman Coulter, Indianapolis, IN, USA), quantified using Qubit dsDNA HS assay kit on a Qubit 3.0 fluorometer (Invitrogen, Carlsbad, CA, USA), and diluted to 10 nM in 30 μL total volume before submitting to the QB3 Vincent J. Coates Genomics Sequencing Laboratory facility at the University of California, Berkeley for sequencing using Illumina Miseq 300 bp pair-end with v3 chemistry.

### Amplicon sequence processing and analysis

16S amplicon sequencing reads were demultiplexed in QIIME2 (86) and then passed to DADA2 (87) to generate Amplicon Sequence Variants (ASVs), with taxonomies assigned using the August 2013 version of GreenGenes 16S rRNA gene database as described previously (16). All subsequent 16S statistical analyses were performed in R-v3.6.1 (88). To account for differences in sequencing read depth across samples, samples were normalized by dividing the reads per ASV in a sample by the sum of usable reads in that sample, resulting in a table of relative abundance frequencies, which were used for analyses, with the exception of alpha-diversity calculations, for which all samples were normalized to an even read depth of 29,918 ASVs per sample. Alpha diversity was determined with the estimate_richness function in the R package phyloseq-v1.30.0 (89), and significance was tested by ANOVA using the aov function in the R stats package. Beta diversity (PCoA) was performed using the ordinate function in the R package phyloseq-v1.30.0 (89). Sample type separation was determined by pairwise PERMANOVA with 1,000 permutations using the adonis and calc_pairwise_permanovas functions in the R packages vegan-v2.5.6 (90) and mctoolsr-v0.1.1.2. Tukey-HSD tests used the HSD.test function in the R package Agricolae-v1.3.1. The combined metabolite and bacterial ASV heatmaps were generated using the R package pheatmap-v1.0.12.

### Metabolite extraction and LC-MS

Root water content was estimated by lyophilizing root tissue and calculating the difference between wet and dry weights. Root samples for submission were then normalized such that the lightest sample was 0.2 g (wet weight) and each other sample was at least 0.2 g. Rhizosphere and soil water contents were estimated to obtain similar amounts of material. The overall difference in percent water content between samples was minimal (2.5-6.5%). For extraction of polar metabolites from root tissue (0.2-0.3 g wet weight), samples were first lyophilized dry, then 500 μL of methanol was added, followed by a brief vortex and sonication in a water bath for 10 min. Samples were centrifuged 5 min at 5,000 rpm, then supernatant transferred to 2 mL tubes, dried in a SpeedVac (SPD111V, Thermo Scientific, Waltham, MA), and extracts stored at −80 °C. For soil and rhizosphere samples (1.25 g wet weight), polar metabolites were extracted similarly but samples were not lyophilized prior to extraction, 2 mL LC-MS grade water was added followed by vortex and water bath sonication for 30 min, centrifugation for 7 min at 7,000 rpm, then supernatant transferred to a 5 mL tube, frozen and lyophilized dry, and extracts stored at −80 °C.

In preparation for LC-MS, soil and rhizosphere extracts were resuspended with 300 μL methanol containing internal standards (~15 μM average of 5-50 μM of 13C,15N Cell Free Amino Acid Mixture; 4-(3,3-dimethyl-ureido)benzoic acid; 3,6-dihydroxy-4-methylbenzoic acid; d5-benzoic acid; 9-anthracene carboxylic acid; 13C-trehalose; 13C-mannitol), vortexed and sonicated 10 min, centrifuged 5 min at 5,000 rpm, supernatant centrifuge-filtered 2.5 min at 2 500 rpm (0.22 μm hydrophilic PVDF), then 150 μL transferred to LC-MS glass autosampler vials. Root extracts were resuspended similarly, but with resuspension volume varied to normalize by root dry weight.

Chromatography was performed using an Agilent 1,290 LC stack, with MS and MS/MS data collected using a Thermo QExactive Orbitrap MS (Thermo Scientific, Waltham, MA). Full MS spectra were collected from *m/z* 70-1,050 at 70,000 resolution in both positive and negative ion modes, with MS/MS fragmentation data acquired using stepped 10, 20, and 40 eV collision energies at 17,500 resolution. Chromatography was performed using a HILIC column (Agilent InfinityLab Poroshell 120 HILIC-Z, 2.1 x 150 mm, 2.7 μm, #673775-924) at a flow rate of 0.45 mL/min with a 2 μL injection volume.

To detect metabolites, samples were run on the HILIC column at 40 °C equilibrated with 100% buffer B (95:5 ACN:H2O with 5 mM ammonium acetate) for 1 min, diluting buffer B down to 89% with buffer A (100% H2O with 5 mM ammonium acetate and 5 μM methylenediphosphonic acid) over 10 min, down to 70% B over 4.75 min, then down to 20% B over 0.5 min, followed by isocratic elution in 80% buffer A for 2.25 min. Samples consisted of 3 biological replicates each and 3 extraction controls, with sample injection order randomized and an injection blank (2 μL MeOH) run between each sample.

### Metabolite identification and analysis

Metabolite identification was based on exact mass and comparing retention time (RT) and MS/MS fragmentation spectra to that of standards run using the same chromatography and MS/MS method. Custom Python code (91) was used to analyze LC-MS data. For each feature detected (unique *m/z* coupled with RT), a score (0 to 3) was assigned representing the level of confidence in the metabolite identification. Positive identification of a metabolite had detected m/z ≤ 5 ppm or 0.001 Da from theoretical as well as RT ≤ 0.5 min compared to a pure standard run using the same LC-MS method. The highest level of positive identification (score of 3) for a metabolite also had matching MS/MS fragmentation spectra compared to either an outside database (METLIN) (92) or internal database generated from standards run and collected on a QExactive Orbitrap MS. Identifications were invalidated if MS/MS from the sample mismatched that of the standard. MS/MS mirror plots for metabolites are presented in supplemental figure 1.

A total of 112 and 122 polar metabolites were predicted in positive and negative ion modes respectively (supplemental table 1). If a metabolite was observed in both ion modes, the mode with higher peak height was selected for the merged metabolite profile (n=168) used for all analyses. Values below the limit of detection were imputed with the lowest observed values in the dataset rounded down (2,400 or 1,900 for positive or negative ion modes respectively) (supplemental table 2). Principal components analysis of metabolite profiles was performed using the prcomp function in the R stats package. Venn diagram construction utilized Venny-v2.1.0 (93). All other metabolite analyses were performed using MetaboAnalyst-v4.0 (94, 95). Heatmaps were generated using Euclidean distance and Ward clustering algorithms. We evaluated enriched or depleted metabolites with the cutoffs of log_2_ fold change greater than 2 or less than −2, and a p-value of less than 0.05.

### Plant root growth assays

Sterilized seeds of the sorghum cultivar RTx430 were germinated on petri dishes with autoclaved Milli-Q water or autoclaved Milli-Q water containing the defined concentration of Pip overnight in the dark at 28 °C, before being transferred to a growth chamber (28/22 °C, 16 h day, ppf ~250 μmol m^-2^s^-1^). The *Arabidopsis* ecotype Columbia (Col-0) was used in this study. Mutant lines *fmo-1* (SALK_026163) (96), *npr1-1* (CS3726) (97), *rbohd/rbohf* (CS68522) (98), and *azi1-2* (SALK_085727) (70) were obtained from the Arabidopsis Biological Resource Center (99). Sterilized seeds were grown on MS plates containing 1/2× Murashige and Skoog salt mix, 1% sucrose (pH 5.8), 0.8% agar, and the defined concentration of Pip. Plants were first stratified for 3 days at 4°C before being transferred to a growth chamber (21 °C, 16 h day, ppf ~120 μmol m^-2^s^-1^). Root lengths were measured using ImageJ-v1.52a software (100). ANOVA was performed using the aov function in the R stats package and Tukey-HSD tests used the HSD.test function in the R package Agricolae-v1.3.1. SAR pathway image in figure 6b was created with BioRender.com.

## Data availability

All datasets and scripts for analysis are available through github (https://github.com/colemanderr-lab/Caddell-2020) and all short read data can be accessed through NCBI BioProject PRJNA655744. Raw metabolomics data will be made available through the Joint Genome Institute Genome Portal.

## Acknowledgements

We thank Dr. Edi Wipf for their helpful discussions and critical reading of the manuscript.

## Author contributions

D.C. conceived and designed the experiments, performed the experiments, analyzed the data, and prepared figures and/or tables; K.L.,B.B., and T.N. performed the LC-MS, metabolite identification, and analyzed the data; E.P. analyzed the data and prepared figures and/or tables; J.S., J.H., and J.D. performed the field experiment; D.C-D. conceived and designed the experiments and analyzed the data; All authors authored or reviewed drafts of the paper and approved the final draft.

## References cited

1. Zolla G, Badri DV, Bakker MG, Manter DK, Vivanco JM. 2013. Soil microbiomes vary in their ability to confer drought tolerance to Arabidopsis. Appl Soil Ecol 68:1–9.

2. Bulgarelli D, Rott M, Schlaeppi K, Ver Loren van Themaat E, Ahmadinejad N, Assenza F, Rauf P, Huettel B, Reinhardt R, Schmelzer E, Peplies J, Gloeckner FO, Amann R, Eickhorst T, Schulze-Lefert P. 2012. Revealing structure and assembly cues for Arabidopsis root-inhabiting bacterial microbiota. Nature 488:91–95.

3. Lundberg DS, Lebeis SL, Paredes SH, Yourstone S, Gehring J, Malfatti S, Tremblay J, Engelbrektson A, Kunin V, Del Rio TG, Edgar RC, Eickhorst T, Ley RE, Hugenholtz P, Tringe SG, Dangl JL. 2012. Defining the core Arabidopsis thaliana root microbiome. Nature 488:86–90.

4. Peiffer JA, Spor A, Koren O, Jin Z, Tringe SG, Dangl JL, Buckler ES, Ley RE. 2013. Diversity and heritability of the maize rhizosphere microbiome under field conditions. Proc Natl Acad Sci U S A 110:6548–6553.

5. Ofek-Lalzar M, Sela N, Goldman-Voronov M, Green SJ, Hadar Y, Minz D. 2014. Niche and host-associated functional signatures of the root surface microbiome. Nat Commun 5:4950.

6. Naylor D, DeGraaf S, Purdom E, Coleman-Derr D. 2017. Drought and host selection influence bacterial community dynamics in the grass root microbiome. ISME J.

7. Fitzpatrick CR, Copeland J, Wang PW, Guttman DS, Kotanen PM, Johnson MTJ. 2018. Assembly and ecological function of the root microbiome across angiosperm plant species. Proc Natl Acad Sci U S A 115:E1157–E1165.

8. Edwards J, Johnson C, Santos-Medellín C, Lurie E, Podishetty NK, Bhatnagar S, Eisen JA, Sundaresan V. 2015. Structure, variation, and assembly of the root-associated microbiomes of rice. Proc Natl Acad Sci U S A 112:E911–20.

9. Haney CH, Samuel BS, Bush J, Ausubel FM. 2015. Associations with rhizosphere bacteria can confer an adaptive advantage to plants. Nat Plants 1.

10. Walitang DI, Kim C-G, Kim K, Kang Y, Kim YK, Sa T. 2018. The influence of host genotype and salt stress on the seed endophytic community of salt-sensitive and salt-tolerant rice cultivars. BMC Plant Biol 18:51.

11. Chaparro JM, Badri DV, Vivanco JM. 2014. Rhizosphere microbiome assemblage is affected by plant development. ISME J 8:790–803.

12. Edwards JA, Santos-Medellín CM, Liechty ZS, Nguyen B, Lurie E, Eason S, Phillips G, Sundaresan V. 2018. Compositional shifts in root-associated bacterial and archaeal microbiota track the plant life cycle in field-grown rice. PLoS Biol 16:e2003862.

13. Xu L, Naylor D, Dong Z, Simmons T, Pierroz G, Hixson KK, Kim Y-M, Zink EM, Engbrecht KM, Wang Y, Gao C, DeGraaf S, Madera MA, Sievert JA, Hollingsworth J, Birdseye D, Scheller HV, Hutmacher R, Dahlberg J, Jansson C, Taylor JW, Lemaux PG, Coleman-Derr D. 2018. Drought delays development of the sorghum root microbiome and enriches for monoderm bacteria. Proc Natl Acad Sci U S A 201717308.

14. Santos-Medellín C, Edwards J, Liechty Z, Nguyen B, Sundaresan V. 2017. Drought Stress Results in a Compartment-Specific Restructuring of the Rice Root-Associated Microbiomes. MBio 8.

15. Koyama A, Steinweg JM, Haddix ML, Dukes JS, Wallenstein MD. 2018. Soil bacterial community responses to altered precipitation and temperature regimes in an old field grassland are mediated by plants. FEMS Microbiol Ecol 94.

16. Simmons T, Styer AB, Pierroz G, Gonçalves AP, Pasricha R, Hazra AB, Bubner P, Coleman-Derr D. 2020. Drought Drives Spatial Variation in the Millet Root Microbiome. Front Plant Sci 11:599.

17. Baraniya D, Nannipieri P, Kublik S, Vestergaard G, Schloter M, Schöler A. 2018. The Impact of the Diurnal Cycle on the Microbial Transcriptome in the Rhizosphere of Barley. Microb Ecol 75:830–833.

18. Chaparro JM, Badri DV, Bakker MG, Sugiyama A, Manter DK, Vivanco JM. 2013. Root exudation of phytochemicals in Arabidopsis follows specific patterns that are developmentally programmed and correlate with soil microbial functions. PLoS One 8:e55731.

19. Zhalnina K, Louie KB, Hao Z, Mansoori N, da Rocha UN, Shi S, Cho H, Karaoz U, Loqué D, Bowen BP, Firestone MK, Northen TR, Brodie EL. 2018. Dynamic root exudate chemistry and microbial substrate preferences drive patterns in rhizosphere microbial community assembly. Nat Microbiol 3:470–480.

20. Wallace JG, Kremling KA, Kovar LL, Buckler ES. 2018. Quantitative Genetics of the Maize Leaf Microbiome. Phytobiomes Journal 2:208–224.

21. Badri DV, Chaparro JM, Zhang R, Shen Q, Vivanco JM. 2013. Application of natural blends of phytochemicals derived from the root exudates of Arabidopsis to the soil reveal that phenolic-related compounds predominantly modulate the soil microbiome. J Biol Chem 288:4502–4512.

22. Zhang N, Wang D, Liu Y, Li S, Shen Q, Zhang R. 2014. Effects of different plant root exudates and their organic acid components on chemotaxis, biofilm formation and colonization by beneficial rhizosphere-associated bacterial strains. Plant Soil 374:689–700.

23. Huang AC, Jiang T, Liu Y-X, Bai Y-C, Reed J, Qu B, Goossens A, Nützmann H-W, Bai Y, Osbourn A. 2019. A specialized metabolic network selectively modulates Arabidopsis root microbiota. Science 364.

24. Cai T, Cai W, Zhang J, Zheng H, Tsou AM, Xiao L, Zhong Z, Zhu J. 2009. Host legume-exuded antimetabolites optimize the symbiotic rhizosphere. Mol Microbiol 73:507–517.

25. Bressan M, Roncato M-A, Bellvert F, Comte G, Haichar FZ, Achouak W, Berge O. 2009. Exogenous glucosinolate produced by Arabidopsis thaliana has an impact on microbes in the rhizosphere and plant roots. ISME J 3:1243–1257.

26. Iven T, König S, Singh S, Braus-Stromeyer SA, Bischoff M, Tietze LF, Braus GH, Lipka V, Feussner I, Dröge-Laser W. 2012. Transcriptional activation and production of tryptophan-derived secondary metabolites in arabidopsis roots contributes to the defense against the fungal vascular pathogen Verticillium longisporum. Mol Plant 5:1389–1402.

27. Wang Y, Bouwmeester K, van de Mortel JE, Shan W, Govers F. 2013. A novel Arabidopsis-oomycete pathosystem: differential interactions with Phytophthora capsici reveal a role for camalexin, indole glucosinolates and salicylic acid in defence. Plant Cell Environ 36:1192–1203.

28. Wang Z, Zhang J, Wu F, Zhou X. 2018. Changes in rhizosphere microbial communities in potted cucumber seedlings treated with syringic acid. PLoS One 13:e0200007.

29. Hu L, Robert CAM, Cadot S, Zhang X, Ye M, Li B, Manzo D, Chervet N, Steinger T, van der Heijden MGA, Schlaeppi K, Erb M. 2018. Root exudate metabolites drive plant-soil feedbacks on growth and defense by shaping the rhizosphere microbiota. Nat Commun 9:2738.

30. Xu L, Coleman-Derr D. 2019. Causes and consequences of a conserved bacterial root microbiome response to drought stress. Curr Opin Microbiol 49:1–6.

31. Caddell DF, Deng S, Coleman-Derr D. 2019. Role of the Plant Root Microbiome in Abiotic Stress Tolerance, p. 273–311. In Verma, SK, White, Jr, Francis, J (eds.), Seed Endophytes: Biology and Biotechnology. Springer International Publishing, Cham.

32. Bostock RM, Pye MF, Roubtsova TV. 2014. Predisposition in plant disease: exploiting the nexus in abiotic and biotic stress perception and response. Annu Rev Phytopathol 52:517–549.

33. Yasuda M, Ishikawa A, Jikumaru Y, Seki M, Umezawa T, Asami T, Maruyama-Nakashita A, Kudo T, Shinozaki K, Yoshida S, Nakashita H. 2008. Antagonistic interaction between systemic acquired resistance and the abscisic acid-mediated abiotic stress response in Arabidopsis. Plant Cell 20:1678–1692.

34. Varoquaux N, Cole B, Gao C, Pierroz G, Baker CR, Patel D, Madera M, Jeffers T, Hollingsworth J, Sievert J, Yoshinaga Y, Owiti JA, Singan VR, DeGraaf S, Xu L, Blow MJ, Harrison MJ, Visel A, Jansson C, Niyogi KK, Hutmacher R, Coleman-Derr D, O’Malley RC, Taylor JW, Dahlberg J, Vogel JP, Lemaux PG, Purdom E. 2019. Transcriptomic analysis of field-droughted sorghum from seedling to maturity reveals biotic and metabolic responses. Proc Natl Acad Sci U S A.

35. Sheflin AM, Chiniquy D, Yuan C, Goren E, Kumar I, Braud M, Brutnell T, Eveland AL, Tringe S, Liu P, Kresovich S, Marsh EL, Schachtman DP, Prenni JE. 2019. Metabolomics of sorghum roots during nitrogen stress reveals compromised metabolic capacity for salicylic acid biosynthesis. Plant Direct 3:e00122.

36. Castrillo G, Teixeira PJPL, Paredes SH, Law TF, de Lorenzo L, Feltcher ME, Finkel OM, Breakfield NW, Mieczkowski P, Jones CD, Paz-Ares J, Dangl JL. 2017. Root microbiota drive direct integration of phosphate stress and immunity. Nature 543:513–518.

37. Hein JW, Wolfe GV, Blee KA. 2008. Comparison of rhizosphere bacterial communities in Arabidopsis thaliana mutants for systemic acquired resistance. Microb Ecol 55:333–343.

38. Carvalhais LC, Dennis PG, Badri DV, Kidd BN, Vivanco JM, Schenk PM. 2015. Linking Jasmonic Acid Signaling, Root Exudates, and Rhizosphere Microbiomes. Mol Plant Microbe Interact 28:1049–1058.

39. Liu H, Carvalhais LC, Schenk PM, Dennis PG. 2017. Effects of jasmonic acid signalling on the wheat microbiome differ between body sites. Sci Rep 7:41766.

40. Lebeis SL, Paredes SH, Lundberg DS, Breakfield N, Gehring J, McDonald M, Malfatti S, Glavina del Rio T, Jones CD, Tringe SG, Dangl JL. 2015. PLANT MICROBIOME. Salicylic acid modulates colonization of the root microbiome by specific bacterial taxa. Science 349:860–864.

41. Návarová H, Bernsdorff F, Döring A-C, Zeier J. 2012. Pipecolic acid, an endogenous mediator of defense amplification and priming, is a critical regulator of inducible plant immunity. Plant Cell 24:5123–5141.

42. Bernsdorff F, Döring A-C, Gruner K, Schuck S, Bräutigam A, Zeier J. 2016. Pipecolic Acid Orchestrates Plant Systemic Acquired Resistance and Defense Priming via Salicylic Acid-Dependent and – Independent Pathways. Plant Cell 28:102–129.

43. Bowen BP, Northen TR. 2010. Dealing with the unknown: metabolomics and metabolite atlases. J Am Soc Mass Spectrom 21:1471–1476.

44. Cámara B, Bielecki P, Kaminski F, dos Santos VM, Plumeier I, Nikodem P, Pieper DH. 2007. A gene cluster involved in degradation of substituted salicylates via ortho cleavage in Pseudomonas sp. strain MT1 encodes enzymes specifically adapted for transformation of 4-methylcatechol and 3-methylmuconate. J Bacteriol 189:1664–1674.

45. Zhang K, Halitschke R, Yin C, Liu C-J, Gan S-S. 2013. Salicylic acid 3-hydroxylase regulates Arabidopsis leaf longevity by mediating salicylic acid catabolism. Proc Natl Acad Sci U S A 110:14807–14812.

46. Philip-Hollingsworth$ S, Hollingsworth RI, Dazzo ST FB. 1991. N-Acetylglutamic Acid: An Extracellular nod Signal of Rhizobium trifolii ANUS43 That Induces Root Hair Branching and Nodule-like Primordia in White Clover Roots. Plan Perspect 16854:16858.

47. Caspersen S, Sundin P, Munro M, Aðalsteinsson S, Hooker JE, Jensén P. 1999. Interactive effects of lettuce (Lactuca sativa L.), irradiance, and ferulic acid in axenic, hydroponic culture. Plant Soil 210:115–126.

48. Sakamoto A, Murata N. 2000. Genetic engineering of glycinebetaine synthesis in plants: current status and implications for enhancement of stress tolerance. J Exp Bot 51:81–88.

49. Moulin M, Deleu C, Larher F, Bouchereau A. 2006. The lysine-ketoglutarate reductase-saccharopine dehydrogenase is involved in the osmo-induced synthesis of pipecolic acid in rapeseed leaf tissues. Plant Physiol Biochem 44:474–482.

50. Goas G, Goas M, Larher F. 1976. Formation de l’acide pipécolique chez Triglochin maritima. Can J Bot 54:1221–1227.

51. Bown AW, Shelp BJ. 2016. Plant GABA: Not Just a Metabolite. Trends Plant Sci 21:811–813.

52. Nourimand M, Todd CD. 2017. Allantoin contributes to the stress response in cadmium-treated Arabidopsis roots. Plant Physiol Biochem 119:103–109.

53. Irani S, Todd CD. 2018. Exogenous allantoin increases Arabidopsis seedlings tolerance to NaCl stress and regulates expression of oxidative stress response genes. J Plant Physiol 221:43–50.

54. Khan N, Bano A, Babar MA. 2019. Metabolic and physiological changes induced by plant growth regulators and plant growth promoting rhizobacteria and their impact on drought tolerance in Cicer arietinum L. PLoS One 14:e0213040.

55. Oney-Birol S. 2019. Exogenous L-Carnitine Promotes Plant Growth and Cell Division by Mitigating Genotoxic Damage of Salt Stress. Sci Rep 9:17229.

56. Ranieri A, Bernardi R, Lanese P, Soldatini GF. 1989. Changes in free amino acid content and protein pattern of maize seedlings under water stress. Environ Exp Bot 29:351–357.

57. Rai VK. 2002. Role of amino acids in plant responses to stresses. Biol Plant.

58. Fang Y, Xiong L. 2015. General mechanisms of drought response and their application in drought resistance improvement in plants. Cell Mol Life Sci 72:673–689.

59. Gargallo-Garriga A, Preece C, Sardans J, Oravec M, Urban O, Peñuelas J. 2018. Root exudate metabolomes change under drought and show limited capacity for recovery. Sci Rep 8:12696.

60. Fàbregas N, Fernie AR. 2019. The metabolic response to drought. J Exp Bot 70:1077–1085.

61. Conn VM, Walker AR, Franco CMM. 2008. Endophytic actinobacteria induce defense pathways in Arabidopsis thaliana. Mol Plant Microbe Interact 21:208–218.

62. Korenblum E, Dong Y, Szymanski J, Panda S, Jozwiak A, Massalha H, Meir S, Rogachev I, Aharoni A. 2020. Rhizosphere microbiome mediates systemic root metabolite exudation by root-to-root signaling. Proc Natl Acad Sci U S A 117:3874–3883.

63. Wang C, Liu R, Lim G-H, de Lorenzo L, Yu K, Zhang K, Hunt AG, Kachroo A, Kachroo P. 2018. Pipecolic acid confers systemic immunity by regulating free radicals. Sci Adv 4:eaar4509.

64. Tsuji W, Inanaga S, Araki H, Morita S, An P, Sonobe K. 2005. Development and Distribution of Root System in Two GrainSorghum Cultivars Originated from Sudan under Drought Stress. Plant Prod Sci 8:553–562.

65. Sebastian J, Yee M-C, Goudinho Viana W, Rellán-Álvarez R, Feldman M, Priest HD, Trontin C, Lee T, Jiang H, Baxter I, Mockler TC, Hochholdinger F, Brutnell TP, Dinneny JR. 2016. Grasses suppress shoot-borne roots to conserve water during drought. Proc Natl Acad Sci U S A 113:8861–8866.

66. Wang Y, Schuck S, Wu J, Yang P, Döring A-C, Zeier J, Tsuda K. 2018. A MPK3/6-WRKY33-ALD1-Pipecolic Acid Regulatory Loop Contributes to Systemic Acquired Resistance. Plant Cell 30:2480–2494.

67. Hartmann M, Zeier J. 2019. N-hydroxypipecolic acid and salicylic acid: a metabolic duo for systemic acquired resistance. Curr Opin Plant Biol 50:44–57.

68. Hartmann M, Zeier T, Bernsdorff F, Reichel-Deland V, Kim D, Hohmann M, Scholten N, Schuck S, Bräutigam A, Hölzel T, Ganter C, Zeier J. 2018. Flavin Monooxygenase-Generated N-Hydroxypipecolic Acid Is a Critical Element of Plant Systemic Immunity. Cell 173:456–469.e16.

69. Cao H, Bowling SA, Gordon AS, Dong X. 1994. Characterization of an Arabidopsis Mutant That Is Nonresponsive to Inducers of Systemic Acquired Resistance. Plant Cell 6:1583–1592.

70. Jung HW, Tschaplinski TJ, Wang L, Glazebrook J, Greenberg JT. 2009. Priming in systemic plant immunity. Science 324:89–91.

71. Wang C, El-Shetehy M, Shine MB, Yu K, Navarre D, Wendehenne D, Kachroo A, Kachroo P. 2014. Free radicals mediate systemic acquired resistance. Cell Rep 7:348–355.

72. Henry A, Doucette W, Norton J, Bugbee B. 2007. Changes in crested wheatgrass root exudation caused by flood, drought, and nutrient stress. J Environ Qual 36:904–912.

73. Preece C, Farré-Armengol G, Llusià J, Peñuelas J. 2018. Thirsty tree roots exude more carbon. Tree Physiol 38:690–695.

74. You J, Zhang Y, Liu A, Li D, Wang X, Dossa K, Zhou R, Yu J, Zhang Y, Wang L, Zhang X. 2019. Transcriptomic and metabolomic profiling of drought-tolerant and susceptible sesame genotypes in response to drought stress. BMC Plant Biol 19:267.

75. Holmes EC, Chen Y-C, Sattely E, Mudgett MB. 2019. Conservation of N-hydroxy-pipecolic acid-mediated systemic acquired resistance in crop plants. bioRxiv.

76. He Q, Zhao S, Ma Q, Zhang Y, Huang L, Li G, Hao L. 2014. Endogenous Salicylic Acid Levels and Signaling Positively Regulate Arabidopsis Response to Polyethylene Glycol-Simulated Drought Stress. J Plant Growth Regul 33:871–880.

77. La VH, Lee B-R, Islam MT, Park S-H, Jung H-I, Bae D-W, Kim T-H. 2019. Characterization of salicylic acid-mediated modulation of the drought stress responses: Reactive oxygen species, proline, and redox state in Brassica napus. Environ Exp Bot 157:1–10.

78. Ding Y, Dommel M, Mou Z. 2016. Abscisic acid promotes proteasome-mediated degradation of the transcription coactivator NPR 1 in Arabidopsis thaliana. Plant J 86:20–34.

79. Schnake A, Hartmann M, Schreiber S, Malik J, Brahmann L, Yildiz I, von Dahlen J, Rose LE, Schaffrath U, Zeier J. 2020. Inducible biosynthesis and immune function of the systemic acquired resistance inducer N-hydroxypipecolic acid in monocotyledonous and dicotyledonous plants. J Exp Bot.

80. Gouesbet G, Blanco C, Hamelin J, Bernard T. 1992. Osmotic adjustment in Brevibacterium ammoniagenes: pipecolic acid accumulation at elevated osmolalities. J Gen Microbiol 138:959–965.

81. Gouesbet G, Jebbar M, Talibart R, Bernard T, Blanco C. 1994. Pipecolic acid is an osmoprotectant for Escherichia coli taken up by the general osmoporters ProU and ProP. Microbiology 140 (Pt 9):2415–2422.

82. Gouffi K, Bernard T, Blanco C. 2000. Osmoprotection by pipecolic acid in Sinorhizobium meliloti: specific effects of D and L isomers. Appl Environ Microbiol 66:2358–2364.

83. Neshich IAP, Kiyota E, Arruda P. 2013. Genome-wide analysis of lysine catabolism in bacteria reveals new connections with osmotic stress resistance. ISME J 7:2400–2410.

84. Gao C, Montoya L, Xu L, Madera M, Hollingsworth J, Purdom E, Singan V, Vogel J, Hutmacher RB, Dahlberg JA, Coleman-Derr D, Lemaux PG, Taylor JW. 2020. Fungal community assembly in drought-stressed sorghum shows stochasticity, selection, and universal ecological dynamics. Nat Commun 11:34.

85. Simmons T, Caddell DF, Deng S, Coleman-Derr D. 2018. Exploring the Root Microbiome: Extracting Bacterial Community Data from the Soil, Rhizosphere, and Root Endosphere. J Vis Exp.

86. Bolyen E, Rideout JR, Dillon MR, Bokulich NA, Abnet CC, Al-Ghalith GA, Alexander H, Alm EJ, Arumugam M, Asnicar F, Bai Y, Bisanz JE, Bittinger K, Brejnrod A, Brislawn CJ, Brown CT, Callahan BJ, Caraballo-Rodríguez AM, Chase J, Cope EK, Da Silva R, Diener C, Dorrestein PC, Douglas GM, Durall DM, Duvallet C, Edwardson CF, Ernst M, Estaki M, Fouquier J, Gauglitz JM, Gibbons SM, Gibson DL, Gonzalez A, Gorlick K, Guo J, Hillmann B, Holmes S, Holste H, Huttenhower C, Huttley GA, Janssen S, Jarmusch AK, Jiang L, Kaehler BD, Kang KB, Keefe CR, Keim P, Kelley ST, Knights D, Koester I, Kosciolek T, Kreps J, Langille MGI, Lee J, Ley R, Liu Y-X, Loftfield E, Lozupone C, Maher M, Marotz C, Martin BD, McDonald D, McIver LJ, Melnik AV, Metcalf JL, Morgan SC, Morton JT, Naimey AT, Navas-Molina JA, Nothias LF, Orchanian SB, Pearson T, Peoples SL, Petras D, Preuss ML, Pruesse E, Rasmussen LB, Rivers A, Robeson MS 2nd, Rosenthal P, Segata N, Shaffer M, Shiffer A, Sinha R, Song SJ, Spear JR, Swafford AD, Thompson LR, Torres PJ, Trinh P, Tripathi A, Turnbaugh PJ, Ul-Hasan S, van der Hooft JJJ, Vargas F, Vázquez-Baeza Y, Vogtmann E, von Hippel M, Walters W, Wan Y, Wang M, Warren J, Weber KC, Williamson CHD, Willis AD, Xu ZZ, Zaneveld JR, Zhang Y, Zhu Q, Knight R, Caporaso JG. 2019. Reproducible, interactive, scalable and extensible microbiome data science using QIIME 2. Nat Biotechnol 37:852–857.

87. Callahan BJ, McMurdie PJ, Rosen MJ, Han AW, Johnson AJA, Holmes SP. 2016. DADA2: High-resolution sample inference from Illumina amplicon data. Nat Methods 13:581–583.

88. Development Core Team RR. 2011. R: A language and environment for statistical computing.

89. McMurdie PJ, Holmes S. 2013. phyloseq: an R package for reproducible interactive analysis and graphics of microbiome census data. PLoS One 8:e61217.

90. Dixon P. 2003. VEGAN, a package of R functions for community ecology. J Veg Sci 14:927–930.

91. Yao Y, Sun T, Wang T, Ruebel O, Northen T, Bowen BP. 2015. Analysis of Metabolomics Datasets with High-Performance Computing and Metabolite Atlases. Metabolites 5:431–442.

92. Smith CA, O’Maille G, Want EJ, Qin C, Trauger SA, Brandon TR, Custodio DE, Abagyan R, Siuzdak G. 2005. METLIN: a metabolite mass spectral database. Ther Drug Monit 27:747–751.

93. Oliveros JC. 2016. Venny. An interactive tool for comparing lists with Venn’s diagrams. 2007–2015.

94. Chong J, Soufan O, Li C, Caraus I, Li S, Bourque G, Wishart DS, Xia J. 2018. MetaboAnalyst 4.0: towards more transparent and integrative metabolomics analysis. Nucleic Acids Res 46:W486–W494.

95. Chong J, Wishart DS, Xia J. 2019. Using MetaboAnalyst 4.0 for Comprehensive and Integrative Metabolomics Data Analysis. Curr Protoc Bioinformatics 68:e86.

96. Mishina TE, Zeier J. 2006. The Arabidopsis flavin-dependent monooxygenase FMO1 is an essential component of biologically induced systemic acquired resistance. Plant Physiol 141:1666–1675.

97. Cao H, Glazebrook J, Clarke JD, Volko S, Dong X. 1997. The Arabidopsis NPR1 gene that controls systemic acquired resistance encodes a novel protein containing ankyrin repeats. Cell 88:57–63.

98. Torres MA, Dangl JL, Jones JDG. 2002. Arabidopsis gp91phox homologues AtrbohD and AtrbohF are required for accumulation of reactive oxygen intermediates in the plant defense response. Proc Natl Acad Sci U S A 99:517–522.

99. Lamesch P, Berardini TZ, Li D, Swarbreck D, Wilks C, Sasidharan R, Muller R, Dreher K, Alexander DL, Garcia-Hernandez M, Karthikeyan AS, Lee CH, Nelson WD, Ploetz L, Singh S, Wensel A, Huala E. 2012. The Arabidopsis Information Resource (TAIR): improved gene annotation and new tools. Nucleic Acids Res 40:D1202–10.

100. Schneider CA, Rasband WS, Eliceiri KW. 2012. NIH Image to ImageJ: 25 years of image analysis. Nat Methods 9:671–675.

