## Supplemental Figure 1 for "Drought shifts sorghum root metabolite and microbiome profiles and enriches the stress response factor pipecolic acid"

### MSMS Mirror Plots: Contents


| Page # | Index # | Ion mode | Proposed metabolite | Molecular formula | MSMS quality score* |
|---|---|---|---|---|---|
| 4 | 0 | positive | 1-aminocyclopropane-1-carboxylic acid | C4H7NO2 | 1 |
| 5 | 2 | positive | 2-amino-2-methylpropanoic acid | C4H9NO2 | 1 |
| 6 | 3 | positive | 2-hydroxycinnamic acid | C9H8O3 | 0.5 |
| 7 | 4 | positive | 2'-deoxyadenosine | C10H13N5O3 | 1 |
| 8 | 7 | positive | 2'-deoxyguanosine | C10H13N5O4 | 1 |
| 9 | 12 | positive | 4-aminobutanoic acid | C4H9NO2 | 1 |
| 10 | 13 | positive | 4-guanidinobutanoic acid | C5H11N3O2 | 1 |
| 11 | 15 | positive | 4-imidazoleacetic acid | C5H6N2O2 | 1 |
| 12 | 20 | positive | 5'-methylthioadenosine | C11H15N5O3S | 1 |
| 13 | 22 | positive | acetylcholine | C7H16NO2+ | 1 |
| 14 | 23 | positive | adenosine | C10H13N5O4 | 1 |
| 15 | 25 | positive | agmatine sulfuric acid | C5H14N4 | 1 |
| 16 | 26 | positive | allantoin | C4H6N4O3 | 1 |
| 17 | 29 | positive | arginine | C6H14N4O2 | 1 |
| 18 | 30 | positive | asparagine | C4H8N2O3 | 1 |
| 19 | 31 | positive | benzylamine | C7H9N | 1 |
| 20 | 32 | positive | betaine | C5H12NO2+ | 1 |
| 21 | 35 | positive | carnitine | C7H16NO3+ | 1 |
| 22 | 36 | positive | choline | C5H14NO+ | 1 |
| 23 | 37 | positive | choline o-sulfuric acid | C5H14NO4S+ | 1 |
| 24 | 38 | positive | citrulline | C6H13N3O3 | 1 |
| 25 | 41 | positive | cytidine | C9H13N3O5 | 1 |
| 26 | 43 | positive | cytosine | C4H5N3O | 1 |
| 27 | 44 | positive | deoxycarnitine | C7H16NO2+ | 1 |
| 28 | 45 | positive | deoxycytidine | C9H13N3O4 | 1 |
| 29 | 51 | positive | glutamic acid | C5H9NO4 | 1 |
| 30 | 52 | positive | glutamine | C5H10N2O3 | 1 |
| 31 | 54 | positive | guanosine | C10H13N5O5 | 1 |
| 32 | 59 | positive | hypoxanthine | C5H4N4O | 1 |
| 33 | 63 | positive | isoleucine | C6H13NO2 | 1 |
| 34 | 64 | positive | leucine | C6H13NO2 | 1 |
| 35 | 65 | positive | lysine | C6H14N2O2 | 1 |
| 36 | 68 | positive | methionine | C5H11NO2S | 1 |
| 37 | 71 | positive | N-acetyl-glucosamine | C8H15NO6 | 1 |
| 38 | 73 | positive | N-acetylputrescine | C6H14N2O | 1 |
| 39 | 76 | positive | N-trimethyllysine | C9H21N2O2+ | 1 |
| 40 | 77 | positive | nicotinamide | C6H6N2O | 1 |
| 41 | 78 | positive | o-acetyl-serine | C5H9NO4 | -1 |
| 42 | 81 | positive | phenethylamine | C8H11N | 1 |
| 43 | 82 | positive | pipecolic acid | C6H11NO2 | 1 |
| 44 | 83 | positive | proline | C5H9NO2 | 1 |
| 45 | 86 | positive | 3-hydroxyphenylacetic acid// pyridoxine | C8H11NO3 | 0.5 |
| 46 | 89 | positive | raffinose | C18H32O16 | 1 |
| 47 | 92 | positive | serine | C3H7NO3 | 1 |
| 48 | 94 | positive | sn-glycero-3-phosphocholine | C8H21NO6P+ | 1 |
| 49 | 96 | positive | sucrose | C12H22O11 | 1 |
| 50 | 98 | positive | thiamine | C12H17N4OS+ | 1 |
| 51 | 102 | positive | trehalose | C12H22O11 | 1 |
| 52 | 103 | positive | tryptophan | C11H12N2O2 | 1 |
| 53 | 104 | positive | tyrosine | C9H11NO3 | 1 |
| 54 | 105 | positive | uridine | C9H12N2O6 | 1 |
| 55 | 108 | positive | valine | C5H11NO2 | 1 |
| 56 | 109 | positive | xanthine | C5H4N4O2 | 1 |
| 57 | 1 | negative | 2-hydroxycinnamic acid// 4-coumaric acid | C9H8O3 | 1 |
| 58 | 2 | negative | 2-hydroxyphenylacetic acid// 4-hydroxyphenylacetic acid | C8H8O3 | 1 |

| | | | | | |
|---|---|---|---|---|---|
| 59 | 3 | negative | 2-methylglutaric acid// adipic acid | C6H10O4 | 1 |
| 60 | 4 | negative | 2-methylmaleic acid | C5H6O4 | 1 |
| 61 | 7 | negative | 2,3-dihydroxybenzoic acid// 2,5-dihydroxybenzoic acid// 3,4-dihydroxybenzoic acid | C7H6O4 | 1 |
| 62 | 8 | negative | 2'-deoxyadenosine | C10H13N5O3 | 1 |
| 63 | 9 | negative | 2'-deoxyguanosine | C10H13N5O4 | 1 |
| 64 | 10 | negative | 2',3'-cyclic AMP | C10H12N5O6P | 1 |
| 65 | 12 | negative | 3-dehydroshikimic acid | C7H8O5 | 1 |
| 66 | 15 | negative | 3-methyglutaric acid | C6H10O4 | 1 |
| 67 | 17 | negative | 2,3-dihydroxybenzoic acid// 2,5-dihydroxybenzoic acid// 3,4-dihydroxybenzoic acid | C7H6O4 | 0.5 |
| 68 | 20 | negative | 2-hydroxycinnamic acid// 4-coumaric acid | C9H8O3 | 1 |
| 69 | 23 | negative | 4-hydroxybenzoic acid | C7H6O3 | 1 |
| 70 | 24 | negative | 2-hydroxyphenylacetic acid// 4-hydroxyphenylacetic acid | C8H8O3 | 1 |
| 71 | 27 | negative | 5-oxo-proline | C5H7NO3 | 1 |
| 72 | 31 | negative | adenine | C5H5N5 | 1 |
| 73 | 32 | negative | adenosine | C10H13N5O4 | 1 |
| 74 | 33 | negative | 2-methylglutaric acid// adipic acid | C6H10O4 | 1 |
| 75 | 34 | negative | allantoin | C4H6N4O3 | 1 |
| 76 | 35 | negative | allothreonine// homoserine// threonine | C4H9NO3 | 1 |
| 77 | 36 | negative | N-methyl-glutamic acid// alpha-aminoadipic acid | C6H11NO4 | 1 |
| 78 | 38 | negative | arabitol | C5H12O5 | 1 |
| 79 | 40 | negative | asparagine | C4H8N2O3 | 1 |
| 80 | 41 | negative | azelaic acid | C9H16O4 | 1 |
| 81 | 44 | negative | choline o-sulfuric acid | C5H14NO4S+ | 1 |
| 82 | 46 | negative | citrulline | C6H13N3O3 | 1 |
| 83 | 49 | negative | cytidine | C9H13N3O5 | 1 |
| 84 | 51 | negative | deoxycytidine | C9H13N3O4 | 1 |
| 85 | 54 | negative | erythritol | C4H10O4 | 1 |
| 86 | 55 | negative | ferulic acid | C10H10O4 | 1 |
| 87 | 57 | negative | fumaric acid | C4H4O4 | 1 |
| 88 | 58 | negative | galactitol//mannitol | C6H14O6 | 1 |
| 89 | 60 | negative | gluconic acid | C6H12O7 | 1 |
| 90 | 64 | negative | glutamic acid | C5H9NO4 | 1 |
| 91 | 65 | negative | glutamine | C5H10N2O3 | 1 |
| 92 | 66 | negative | guanine | C5H5N5O | 1 |
| 93 | 68 | negative | allothreonine// homoserine// threonine | C4H9NO3 | -1 |
| 94 | 69 | negative | hypoxanthine | C5H4N4O | 1 |
| 95 | 71 | negative | inosine | C10H12N4O5 | 1 |
| 96 | 72 | negative | isocitric acid | C6H8O7 | -1 |
| 97 | 73 | negative | itaconic acid | C5H6O4 | 1 |
| 98 | 74 | negative | lactose// palatinose// sucrose// trehalose | C12H22O11 | 1 |
| 99 | 76 | negative | malonic acid | C3H4O4 | 1 |
| 100 | 77 | negative | galactitol// mannitol | C6H14O6 | 1 |
| 101 | 79 | negative | methionine | C5H11NO2S | 1 |
| 102 | 80 | negative | myo-inositol | C6H12O6 | 1 |
| 103 | 81 | negative | N-acetyl-aspartic acid | C6H9NO5 | 1 |
| 104 | 82 | negative | N-acetyl-glucosamine// N-acetyl-mannosamine | C8H15NO6 | 1 |
| 105 | 83 | negative | N-acetyl-glutamic acid | C7H11NO5 | 1 |
| 106 | 84 | negative | N-acetyl-glucosamine// N-acetyl-mannosamine | C8H15NO6 | 1 |
| 107 | 85 | negative | N-acetyl-serine | C5H9NO4 | 1 |
| 108 | 90 | negative | N-methyl-glutamic acid// alpha-aminoadipic acid | C6H11NO4 | 1 |
| 109 | 93 | negative | orotic acid | C5H4N2O4 | 1 |
| 110 | 94 | negative | lactose// palatinose// sucrose | C12H22O11 | 1 |
| 111 | 95 | negative | phenylalanine | C9H11NO2 | 1 |
| 112 | 96 | negative | proline | C5H9NO2 | 1 |
| 113 | 99 | negative | rhamnose | C6H12O5 | 1 |
| 114 | 101 | negative | shikimic acid | C7H10O5 | 1 |

| | | | | | |
|---|---|---|---|---|---|
| 115 | 103 | negative | lactose// palatinose// sucrose | $C_{12}H_{22}O_{11}$ | 1 |
| 116 | 104 | negative | syringic acid | $C_9H_{10}O_5$ | 1 |
| 117 | 107 | negative | allothreonine// homoserine// threonine | $C_4H_9NO_3$ | 1 |
| 118 | 108 | negative | thymidine | $C_{10}H_{14}N_2O_5$ | 1 |
| 119 | 111 | negative | lactose// trehalose | $C_{12}H_{22}O_{11}$ | 1 |
| 120 | 112 | negative | tryptophan | $C_{11}H_{12}N_2O_2$ | 1 |
| 121 | 113 | negative | tyrosine | $C_9H_{11}NO_3$ | 1 |
| 122 | 114 | negative | uracil | $C_4H_4N_2O_2$ | 1 |
| 123 | 115 | negative | uric acid | $C_5H_4N_4O_3$ | 1 |
| 124 | 116 | negative | uridine | $C_9H_{12}N_2O_6$ | 1 |
| 125 | 118 | negative | valine | $C_5H_{11}NO_2$ | 1 |
| 126 | 119 | negative | vanillic acid | $C_8H_8O_4$ | 1 |
| 127 | 120 | negative | xanthine | $C_5H_4N_4O_2$ | 1 |
| 128 | 121 | negative | xanthosine | $C_{10}H_{12}N_4O_6$ | 1 |

*MSMS quality scores: 1 (MSMS matches ref. std.); 0.5 (possible match); -1 (MSMS poor match to ref. std.); 0 (not included in mirror plots, no MSMS collected or no appropriate ref available)

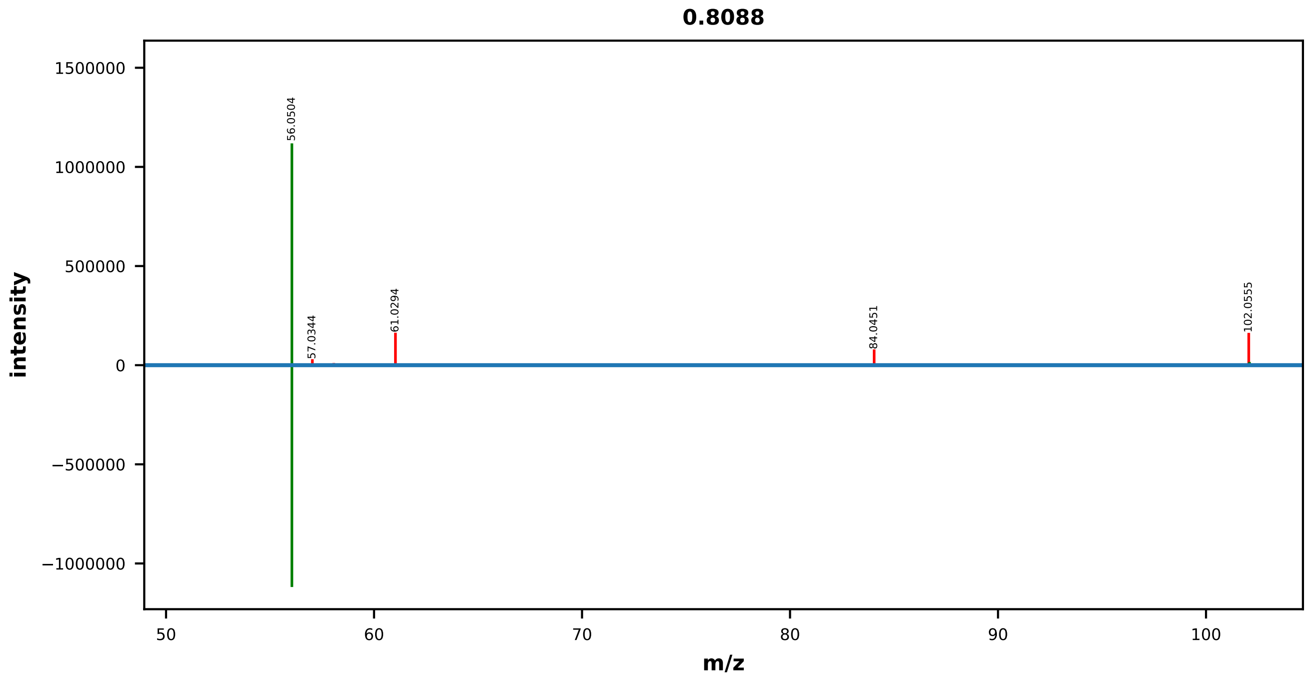


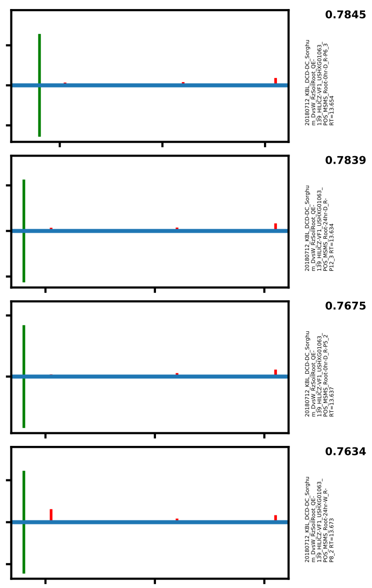


| | EMA Compound Info |
|---|---|
| Name: | 1-Aminocyclopropanecarboxyli c acid |
| Label: | |
| Formula: | C4H7NO2 |
| Polarity: | positive |
| Monoisotopic Mass: | 101.047678464 |
| Theoretical M/Z: | 102.0549785 |
| Adduct: | |

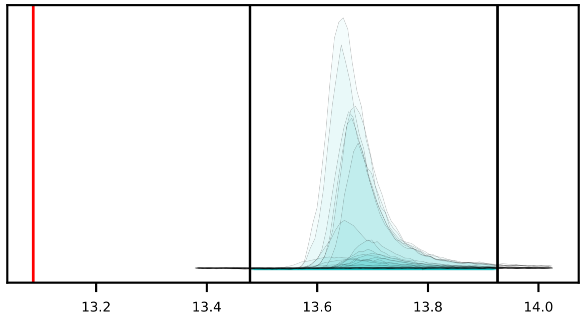

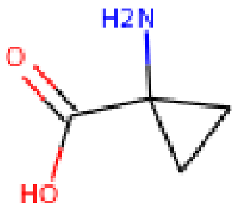


20180712_KBL_DCD-DC_Sorghum_DvsW_RzSoilRoot_QE-139_HILICZ-VF1_USHXG01063_POS_MSMS_Root-24hr-D_R-P10_1

0000_1-aminocyclopropane-1-carboxylic_acid_positive_102p0550_13p09

Measured M/Z = 102.0554, 4.2361 ppm difference

Expected Elution of 13.09 minutes, 13.65 min actual

MSMS Scan at 13.760 minutes

Matching M/Zs above 1E-3*max: 56.050, 102.092, 102.128

All Matching M/Zs: 56.050, 102.092, 102.128

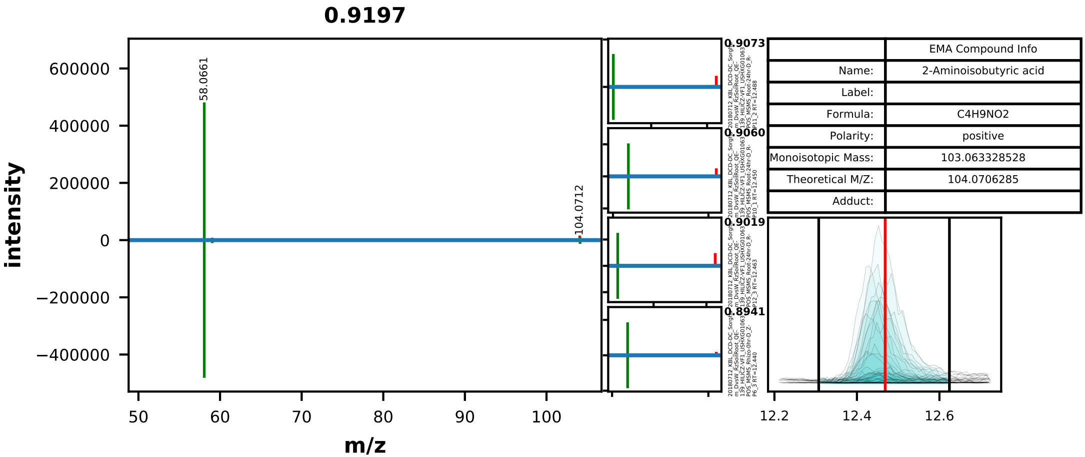


| | EMA Compound Info |
|---|---|
| Name: | 2-Aminoisobutyric acid |
| Label: | |
| Formula: | C4H9NO2 |
| Polarity: | positive |
| Monoisotopic Mass: | 103.063328528 |
| Theoretical M/Z: | 104.0706285 |
| Adduct: | |

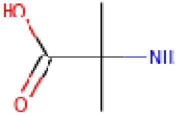


20180712_KBL_DCD-DC_Sorghum_DvsW_RzSoilRoot_QE-139_HILICZ-VF1_USHXG01063_POS_MSMS_Rhizo-0hr-D_Z-P5_2

0002_2-amino-2-methylpropanoic_acid_positive_104p0706_12p47
Measured M/Z = 104.0711, 4.9310 ppm difference
Expected Elution of 12.47 minutes, 12.43 min actual

MSMS Scan at 12.428 minutes

Matching M/Zs above 1E-3*max: 58.066, 59.050, 87.045, 104.108

All Matching M/Zs: 58.066, 59.050, 87.045, 104.108

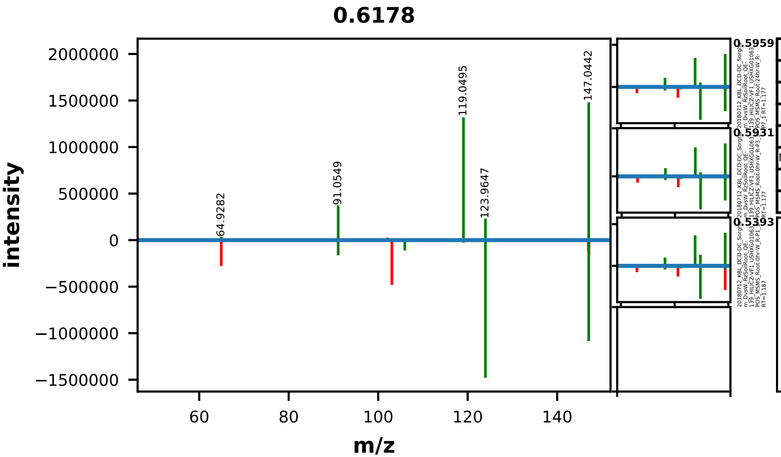


| | EMA Compound Info |
|---|---|
| Name: | 2-Hydroxycinnamic acid |
| Label: | |
| Formula: | C9H8O3 |
| Polarity: | positive |
| Monoisotopic Mass: | 164.047344116 |
| Theoretical M/Z: | 147.0440441 |
| Adduct: | |

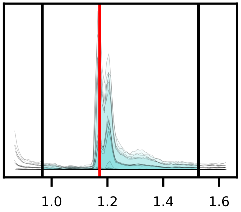

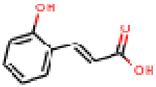


20180712_KBL_DCD-DC_Sorghum_DvsW_RzSoilRoot_QE-139_HILICZ-VF1_USHXG01063_POS_MSMS_Root-24hr-W_R-P8_2

0003_2-hydroxycinnamic_acid_positive_147p0440_1p17
Measured M/Z = 147.0442, 0.7564 ppm difference
Expected Elution of 1.17 minutes, 1.16 min actual

MSMS Scan at 1.178 minutes

Matching M/Zs above 1E-3*max: 64.928, 91.055, 105.071, 105.954, 119.049, 123.965, 146.980, 147.044, 147.117

All Matching M/Zs: 64.928, 91.055, 105.071, 105.954, 119.049, 123.965, 146.980, 147.044, 147.117

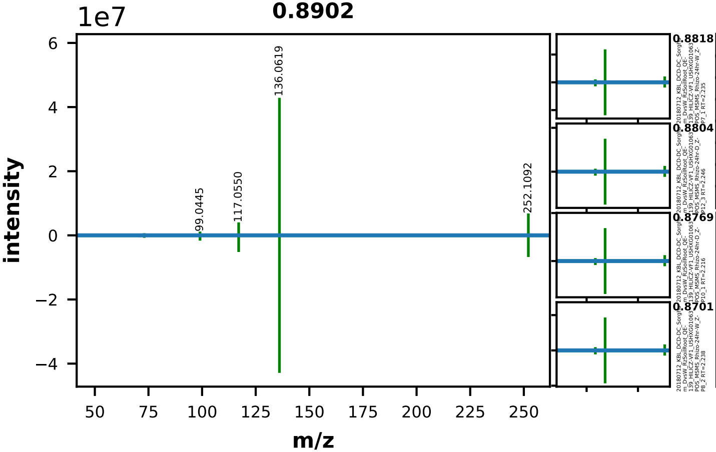


| | EMA Compound Info |
|---|---|
| Name: | 2'-deoxyadenosine |
| Label: | |
| Formula: | C10H13N5O3 |
| Polarity: | positive |
| Monoisotopic Mass: | 251.101839276 |
| Theoretical M/Z: | 252.1091393 |
| Adduct: | |

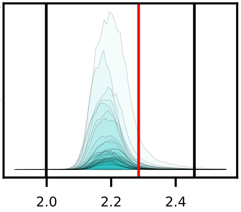

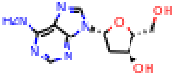


20180712_KBL_DCD-DC_Sorghum_DvsW_RzSoilRoot_QE-139_HILICZ-VF1_USHXG01063_POS_MSMS_Rhizo-0hr-D_Z-P6_3

0004_2deoxyadenosine_positive_252p1091_2p29
Measured M/Z = 252.1092, 0.1260 ppm difference
Expected Elution of 2.29 minutes, 2.19 min actual

MSMS Scan at 2.233 minutes

Matching M/Zs above 1E-3*max: 69.034, 71.050, 73.029, 81.034, 99.045, 117.055, 136.062, 252.109

All Matching M/Zs: 69.034, 71.050, 73.029, 81.034, 99.045, 117.055, 136.062, 252.109

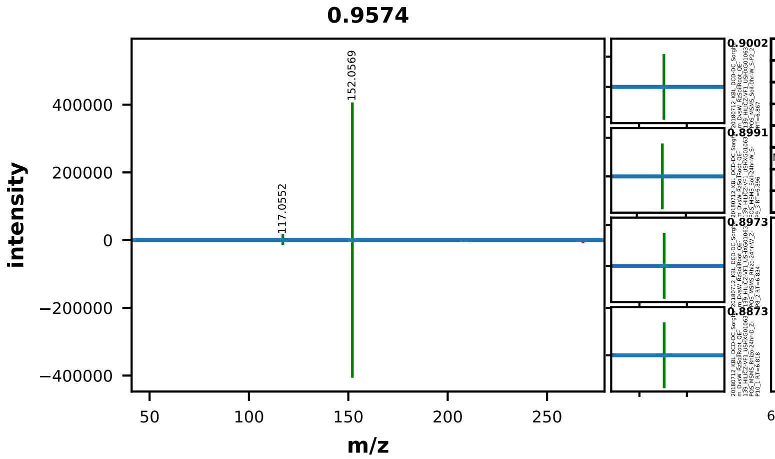


| | EMA Compound Info |
|---|---|
| Name: | 2'-deoxyguanosine |
| Label: | |
| Formula: | C10H13N5O4 |
| Polarity: | positive |
| Monoisotopic Mass: | 267.096753896 |
| Theoretical M/Z: | 268.1040539 |
| Adduct: | |

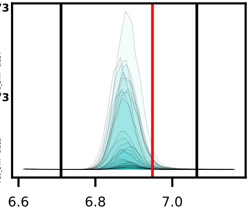


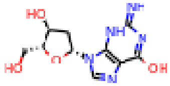

20180712_KBL_DCD-DC_Sorghum_DvsW_RzSoilRoot_QE-139_HILICZ-VF1_USHXG01063_POS_MSMS_Root-0hr-W_R-P1_1

0007_2deoxyguanosine_positive_268p1041_6p95
Measured M/Z = 268.1041, 0.0507 ppm difference
Expected Elution of 6.95 minutes, 6.89 min actual

MSMS Scan at 6.904 minutes

Matching M/Zs above 1E-3*max: 117.055, 152.057

All Matching M/Zs: 117.055, 152.057

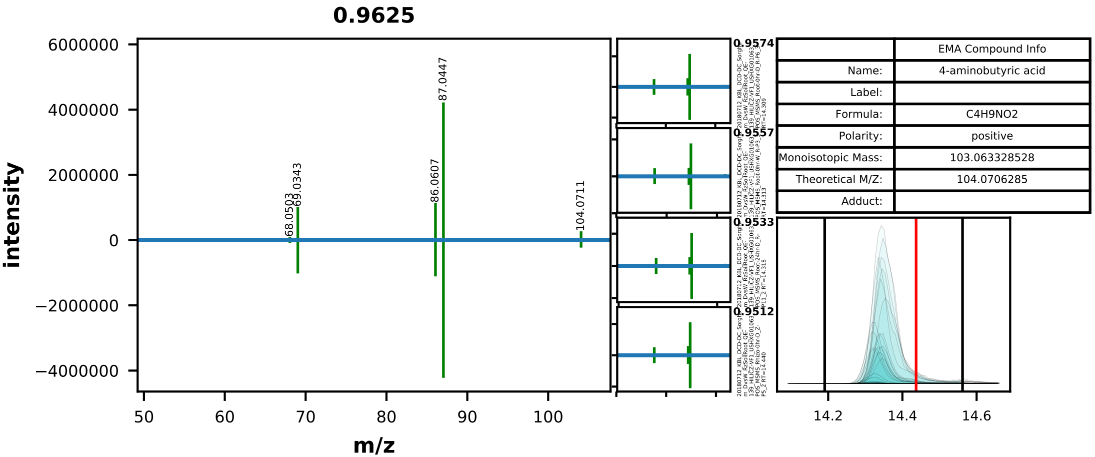


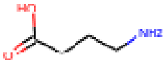


20180712_KBL_DCD-DC_Sorghum_DvsW_RzSoilRoot_QE-139_HILICZ-VF1_USHXG01063_POS_MSMS_Root-24hr-W_R-P9_3

0012_4-aminobutanoic_acid_positive_104p0706_14p44
Measured M/Z = 104.0710, 3.6933 ppm difference
Expected Elution of 14.44 minutes, 14.35 min actual

MSMS Scan at 14.325 minutes

Matching M/Zs above 1E-3*max: 53.003, 68.050, 69.034, 86.061, 87.045, 104.071, 104.107

All Matching M/Zs: 53.003, 68.050, 69.034, 86.061, 87.045, 104.071, 104.107

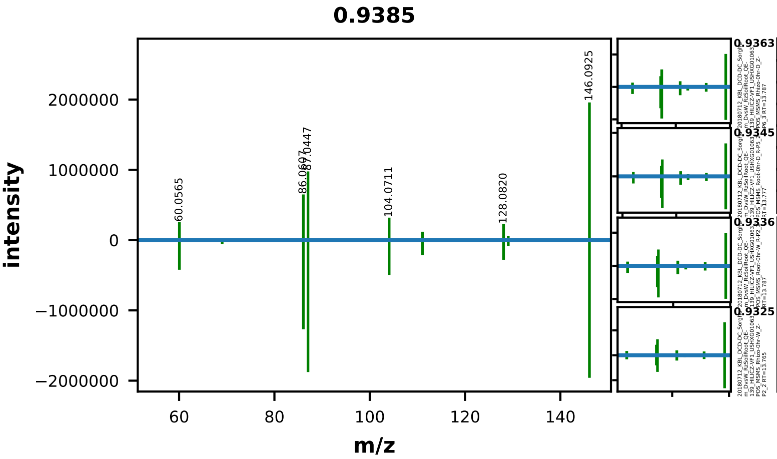


| | EMA Compound Info |
|---|---|
| Name: | 4-Guanidinobutyric acid |
| Label: | |
| Formula: | C5H11N3O2 |
| Polarity: | positive |
| Monoisotopic Mass: | 145.085126592 |
| Theoretical M/Z: | 146.0924266 |
| Adduct: | |

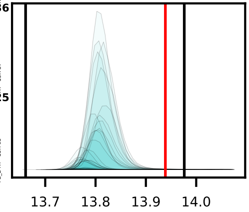


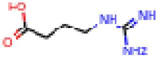


20180712_KBL_DCD-DC_Sorghum_DvsW_RzSoilRoot_QE-139_HILICZ-VF1_USHXG01063_POS_MSMS_Root-0hr-W_R-P3_3

0013_4-guanidinobutanoic_acid_positive_146p0924_13p94
Measured M/Z = 146.0925, 0.3536 ppm difference
Expected Elution of 13.94 minutes, 13.81 min actual

MSMS Scan at 13.781 minutes

Matching M/Zs above 1E-3*max: 60.057, 69.034, 83.061, 86.061, 87.045, 104.071, 111.056, 128.082, 129.066, 146.092



All Matching M/Zs: 60.057, 69.034, 83.061, 86.061, 87.045, 104.071, 111.056, 128.082, 129.066, 146.092

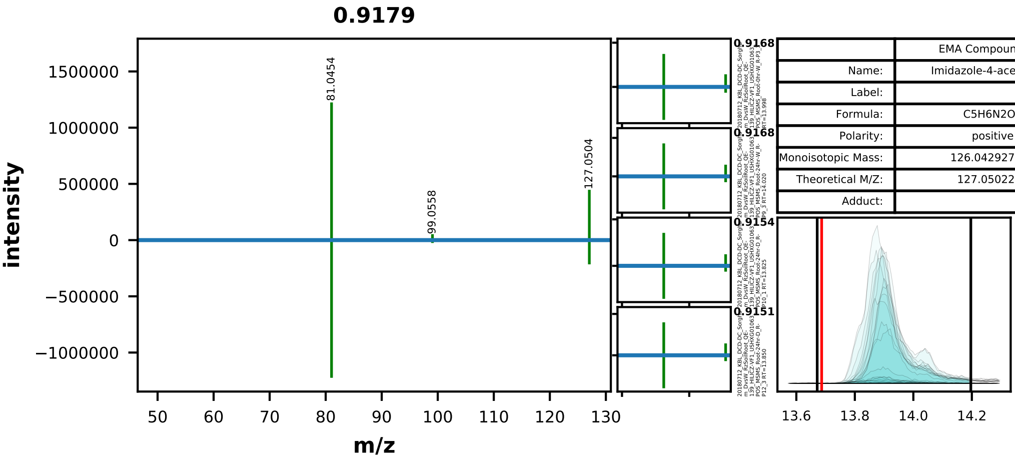


| | EMA Compound Info |
|---|---|
| Name: | Imidazole-4-acetic acid |
| Label: | |
| Formula: | C5H6N2O2 |
| Polarity: | positive |
| Monoisotopic Mass: | 126.042927432 |
| Theoretical M/Z: | 127.0502274 |
| Adduct: | |

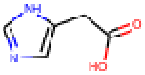

20180712_KBL_DCD-DC_Sorghum_DvsW_RzSoilRoot_QE-139_HILICZ-VF1_USHXG01063_POS_MSMS_Root-0hr-W_R-P2_2

0015_4-imidazoleacetic_acid_positive_127p0502_13p69
Measured M/Z = 127.0504, 1.2444 ppm difference
Expected Elution of 13.69 minutes, 13.90 min actual

MSMS Scan at 13.864 minutes

Matching M/Zs above 1E-3*max: 54.035, 81.045, 99.056, 127.050

All Matching M/Zs: 54.035, 81.045, 99.056, 127.050

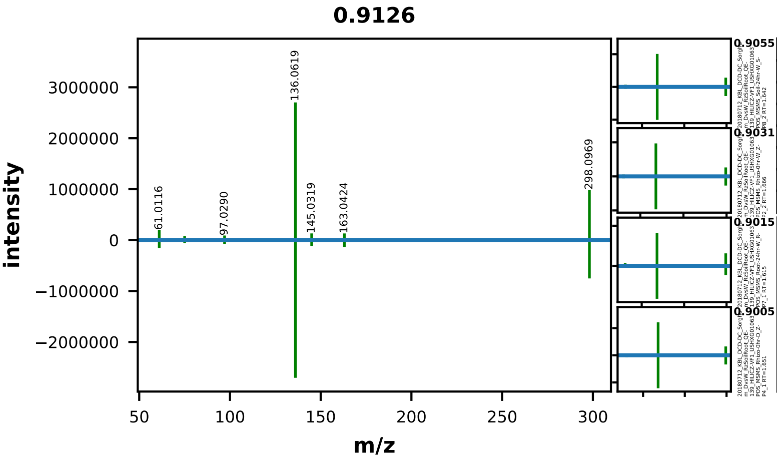


| | EMA Compound Info |
|---|---|
| Name: | Methylthioadenosine |
| Label: | |
| Formula: | C11H15N5O3S |
| Polarity: | positive |
| Monoisotopic Mass: | 297.08956034 |
| Theoretical M/Z: | 298.0968603 |
| Adduct: | |

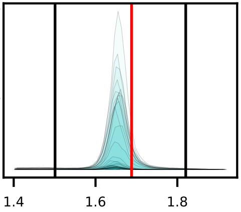


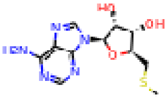


20180712_KBL_DCD-DC_Sorghum_DvsW_RzSoilRoot_QE-139_HILICZ-VF1_USHXG01063_POS_MSMS_Root-24hr-D_R-P10_1

0020_5methylthioadenosine_positive_298p0969_1p69
Measured M/Z = 298.0970, 0.5465 ppm difference
Expected Elution of 1.69 minutes, 1.65 min actual

MSMS Scan at 1.617 minutes

Matching M/Zs above 1E-3*max: 61.012, 69.034, 75.027, 97.029, 103.022, 136.062, 145.032, 163.042, 298.097

All Matching M/Zs: 61.012, 69.034, 75.027, 97.029, 103.022, 136.062, 145.032, 163.042, 298.097

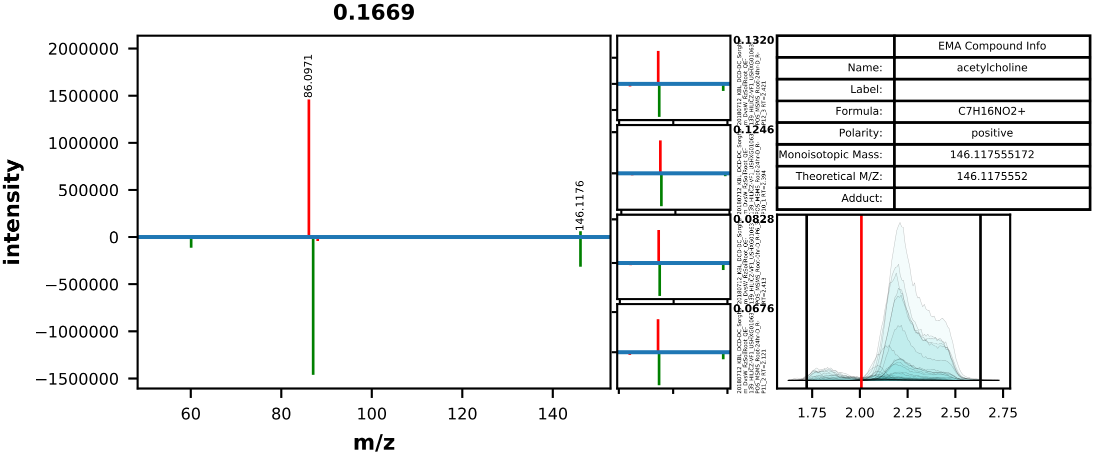


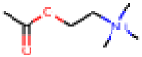

20180712_KBL_DCD-DC_Sorghum_DvsW_RzSoilRoot_QE-139_HILICZ-VF1_USHXG01063_POS_MSMS_Root-0hr-D_R-P5_2

0022_acetylcholine_positive_146p1176_2p01
Measured M/Z = 146.1176, 0.3068 ppm difference
Expected Elution of 2.01 minutes, 2.22 min actual

MSMS Scan at 2.136 minutes

Matching M/Zs above 1E-3*max: 60.081, 87.045, 146.118

All Matching M/Zs: 60.081, 87.045, 146.118

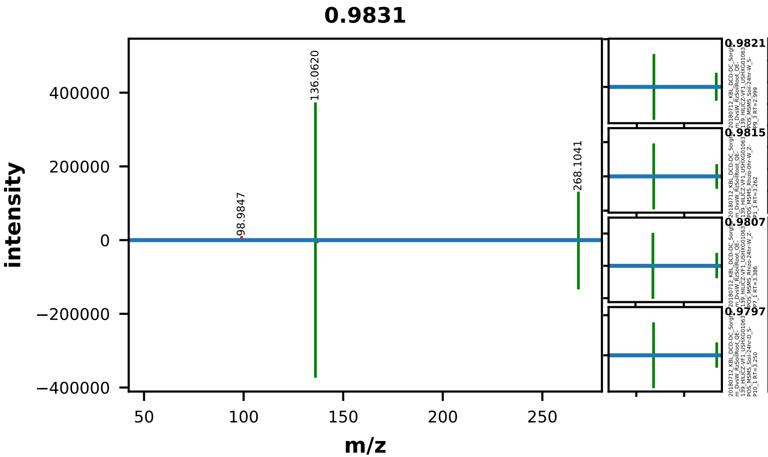


| | EMA Compound Info |
|---|---|
| Name: | adenosine |
| Label: | |
| Formula: | C10H13N5O4 |
| Polarity: | positive |
| Monoisotopic Mass: | 267.096753896 |
| Theoretical M/Z: | 268.1040539 |
| Adduct: | |

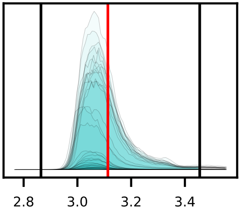


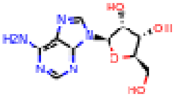


20180712_KBL_DCD-DC_Sorghum_DvsW_RzSoilRoot_QE-139_HILICZ-VF1_USHXG01063_POS_MSMS_Rhizo-24hr-W_Z-P8_2

0023_adenosine_positive_268p1041_3p11
Measured M/Z = 268.1042, 0.3707 ppm difference
Expected Elution of 3.11 minutes, 3.05 min actual

MSMS Scan at 3.401 minutes

Matching M/Zs above 1E-3*max: 136.062, 268.104

All Matching M/Zs: 136.062, 268.104

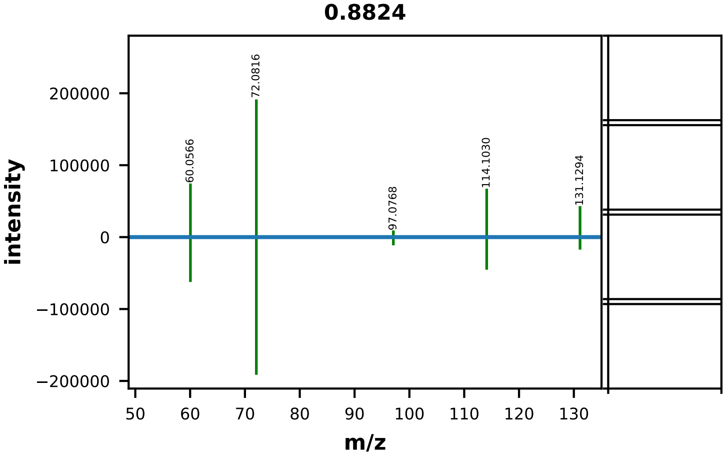


| | EMA Compound Info |
|---|---|
| Name: | agmatine |
| Label: | |
| Formula: | C5H14N4 |
| Polarity: | positive |
| Monoisotopic Mass: | 130.121846448 |
| Theoretical M/Z: | 131.1291464 |
| Adduct: | |

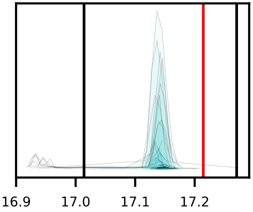


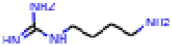

20180712_KBL_DCD-DC_Sorghum_DvsW_RzSoilRoot_QE-139_HILICZ-VF1_USHXG01063_POS_MSMS_Rhizo-24hr-D_Z-P12_3

0025_agmatine_sulfuric_acid_positive_131p1291_17p21
Measured M/Z = 131.1293, 1.3167 ppm difference
Expected Elution of 17.21 minutes, 17.14 min actual

MSMS Scan at 17.151 minutes

Matching M/Zs above 1E-3*max: 60.057, 72.082, 97.077, 114.103, 131.129

All Matching M/Zs: 60.057, 72.082, 97.077, 114.103, 131.129

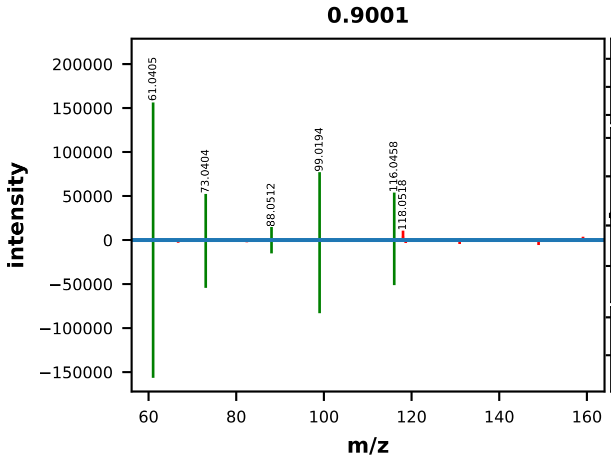


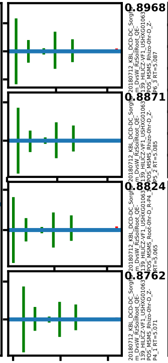


| | EMA Compound Info |
|---|---|
| Name: | allantoin |
| Label: | |
| Formula: | C4H6N4O3 |
| Polarity: | positive |
| Monoisotopic Mass: | 158.043990052 |
| Theoretical M/Z: | 159.0512901 |
| Adduct: | |

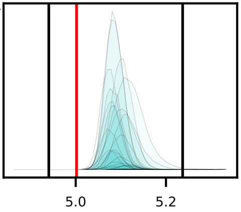

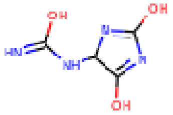


20180712_KBL_DCD-DC_Sorghum_DvsW_RzSoilRoot_QE-139_HILICZ-VF1_USHXG01063_POS_MSMS_Rhizo-24hr-D_Z-P12_3

0026_allantoin_positive_159p0513_5p00
Measured M/Z = 159.0515, 1.1046 ppm difference
Expected Elution of 5.00 minutes, 5.10 min actual

MSMS Scan at 5.079 minutes

Matching M/Zs above 1E-3*max: 61.041, 73.040, 88.051, 99.019, 116.046

All Matching M/Zs: 61.041, 73.040, 88.051, 99.019, 116.046

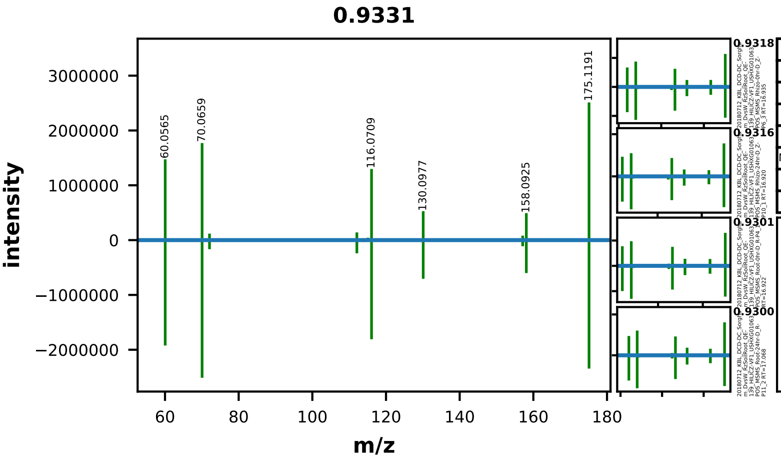


| | EMA Compound Info |
|---|---|
| Name: | L-arginine |
| Label: | |
| Formula: | C6H14N4O2 |
| Polarity: | positive |
| Monoisotopic Mass: | 174.111675688 |
| Theoretical M/Z: | 175.1189757 |
| Adduct: | |

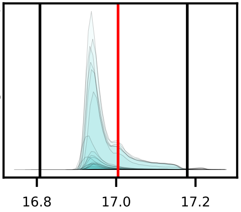


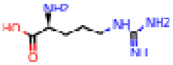

20180712_KBL_DCD-DC_Sorghum_DvsW_RzSoilRoot_QE-139_HILICZ-VF1_USHXG01063_POS_MSMS_Root-24hr-W_R-P7_1

0029_arginine_positive_175p1190_17p00
Measured M/Z = 175.1190, 0.1663 ppm difference
Expected Elution of 17.00 minutes, 16.94 min actual

MSMS Scan at 16.942 minutes

Matching M/Zs above 1E-3*max: 60.057, 70.066, 72.082, 97.076, 112.087, 113.072, 114.102, 115.087, 116.071, 130.098, 157.108, 158.093, 175.119



All Matching M/Zs: 60.057, 70.066, 72.082, 97.076, 112.087, 113.072, 114.102, 115.087, 116.071, 130.098, 157.108, 158.093, 175.119

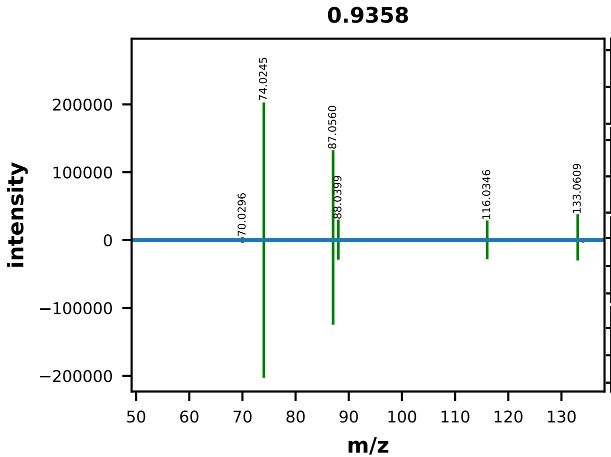


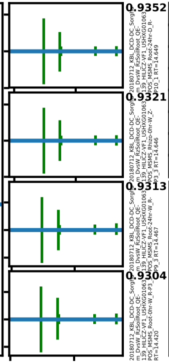


| | EMA Compound Info |
|---|---|
| Name: | L-asparagine |
| Label: | |
| Formula: | C4H8N2O3 |
| Polarity: | positive |
| Monoisotopic Mass: | 132.053492116 |
| Theoretical M/Z: | 133.0607921 |
| Adduct: | |

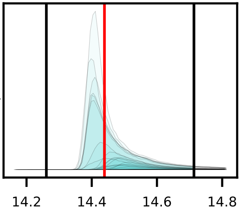


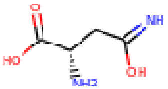


20180712_KBL_DCD-DC_Sorghum_DvsW_RzSoilRoot_QE-139_HILICZ-VF1_USHXG01063_POS_MSMS_Rhizo-24hr-D_Z-P11_2

0030_asparagine_positive_133p0608_14p44
Measured M/Z = 133.0609, 0.7181 ppm difference
Expected Elution of 14.44 minutes, 14.59 min actual

MSMS Scan at 14.557 minutes

Matching M/Zs above 1E-3*max: 70.030, 74.024, 87.056, 88.040, 116.035, 133.061

All Matching M/Zs: 70.030, 74.024, 87.056, 88.040, 116.035, 133.061

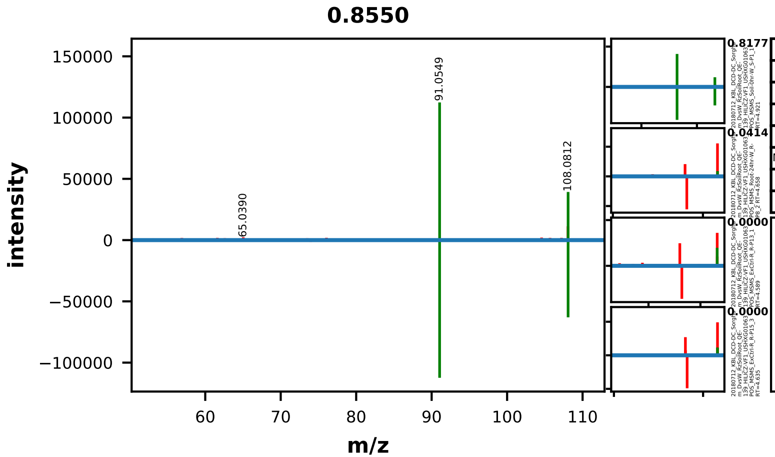


| | EMA Compound Info |
|---|---|
| Name: | BENZYLAMINE |
| Label: | |
| Formula: | C7H9N |
| Polarity: | positive |
| Monoisotopic Mass: | 107.073499288 |
| Theoretical M/Z: | 108.0807993 |
| Adduct: | |

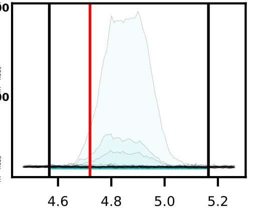


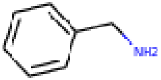


20180712_KBL_DCD-DC_Sorghum_DvsW_RzSoilRoot_QE-139_HILICZ-VF1_USHXG01063_POS_MSMS_Soil-0hr-D_S-P5_2

0031_benzylamine_positive_108p0808_4p72
Measured M/Z = 108.0812, 4.0229 ppm difference
Expected Elution of 4.72 minutes, 4.90 min actual

MSMS Scan at 5.024 minutes

Matching M/Zs above 1E-3*max: 91.055, 108.081

All Matching M/Zs: 91.055, 108.081

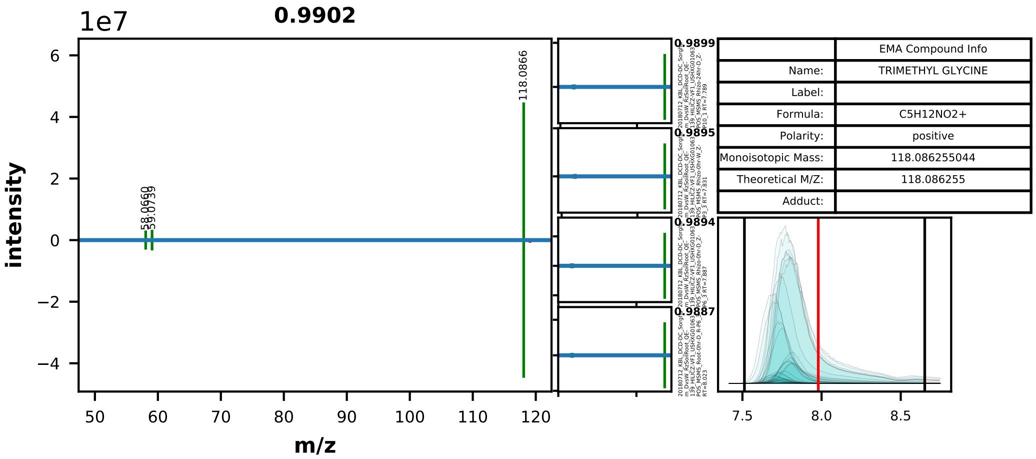


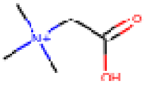


20180712_KBL_DCD-DC_Sorghum_DvsW_RzSoilRoot_QE-139_HILICZ-VF1_USHXG01063_POS_MSMS_Rhizo-24hr-D_Z-P12_3

0032_betaine_positive_118p0863_7p98
Measured M/Z = 118.0866, 2.6444 ppm difference
Expected Elution of 7.98 minutes, 7.75 min actual

MSMS Scan at 8.032 minutes

Matching M/Zs above 1E-3*max: 58.066, 59.074, 118.087

All Matching M/Zs: 58.066, 59.074, 118.087

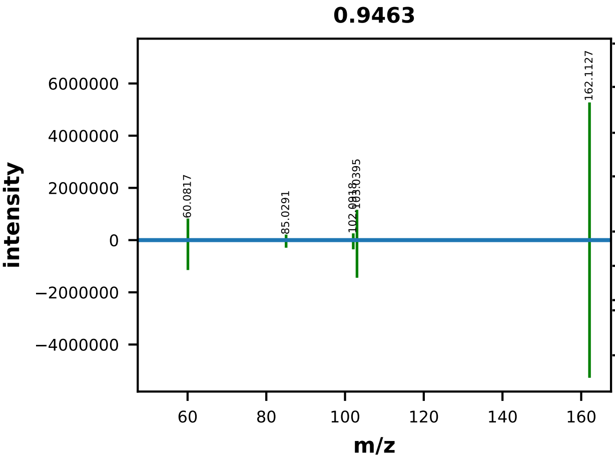


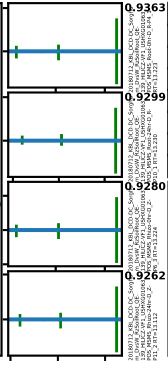


| | EMA Compound Info |
|---|---|
| Name: | CHEBI:39547 |
| Label: | |
| Formula: | C7H16NO3+ |
| Polarity: | positive |
| Monoisotopic Mass: | 162.112469792 |
| Theoretical M/Z: | 162.1124698 |
| Adduct: | |

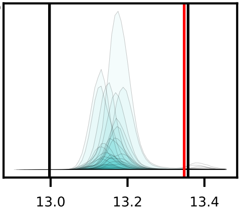


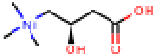

20180712_KBL_DCD-DC_Sorghum_DvsW_RzSoilRoot_QE-139_HILICZ-VF1_USHXG01063_POS_MSMS_Root-24hr-D_R-P11_2

0035_carnitine_positive_162p1125_13p35
Measured M/Z = 162.1126, 0.5329 ppm difference
Expected Elution of 13.35 minutes, 13.19 min actual

MSMS Scan at 13.252 minutes

Matching M/Zs above 1E-3*max: 60.082, 85.029, 102.092, 103.040, 162.113

All Matching M/Zs: 60.082, 85.029, 102.092, 103.040, 162.113

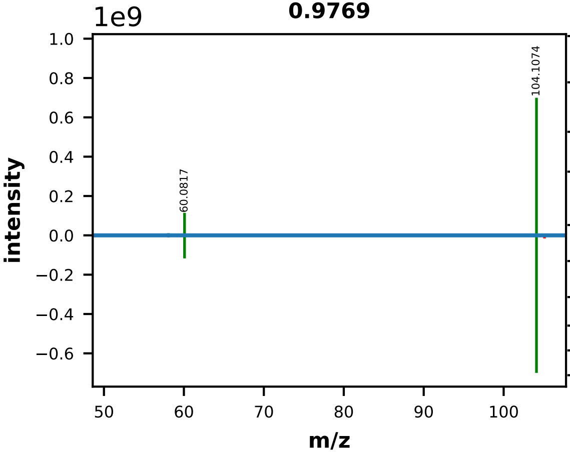


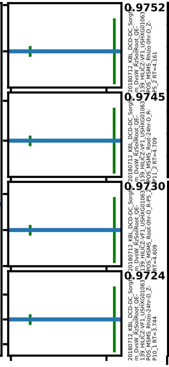


|  | EMA Compound Info |
|---|---|
| Name: | choline |
| Label: |  |
| Formula: | C5H14NO+ |
| Polarity: | positive |
| Monoisotopic Mass: | 104.106990488 |
| Theoretical M/Z: | 104.1069905 |
| Adduct: |  |

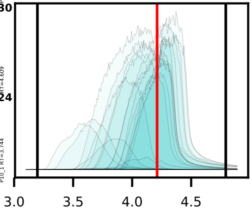

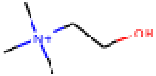

20180712_KBL_DCD-DC_Sorghum_DvsW_RzSoilRoot_QE-
139_HILICZ-VF1_USHXG01063_POS_MSMS_Root-0hr-W_R-P3_3

0036_choline_positive_104p1070_4p21
Measured M/Z = 104.1074, 4.0464 ppm difference
Expected Elution of  4.21 minutes,  4.23 min actual

MSMS Scan at 4.179 minutes

Matching M/Zs above 1E-3*max: 58.066, 59.074, 60.082, 104.107

All Matching M/Zs: 58.066, 59.074, 60.082, 104.107

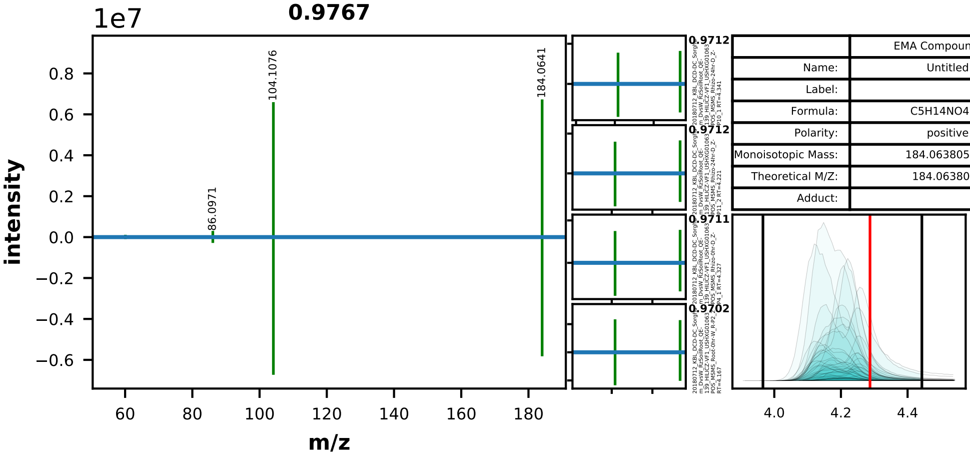


| | EMA Compound Info |
|---|---|
| Name: | Untitled |
| Label: | |
| Formula: | C5H14NO4S+ |
| Polarity: | positive |
| Monoisotopic Mass: | 184.063805348 |
| Theoretical M/Z: | 184.0638053 |
| Adduct: | |

20180712_KBL_DCD-DC_Sorghum_DvsW_RzSoilRoot_QE-139_HILICZ-VF1_USHXG01063_POS_MSMS_Soil-24hr-W_S-P8_2

0037_choline_o-sulfuric_acid_positive_184p0638_4p29
Measured M/Z = 184.0641, 1.7900 ppm difference
Expected Elution of 4.29 minutes, 4.15 min actual

MSMS Scan at 4.226 minutes

Matching M/Zs above 1E-3*max: 58.066, 60.082, 86.097, 104.108, 184.064

All Matching M/Zs: 58.066, 60.082, 86.097, 104.108, 184.064

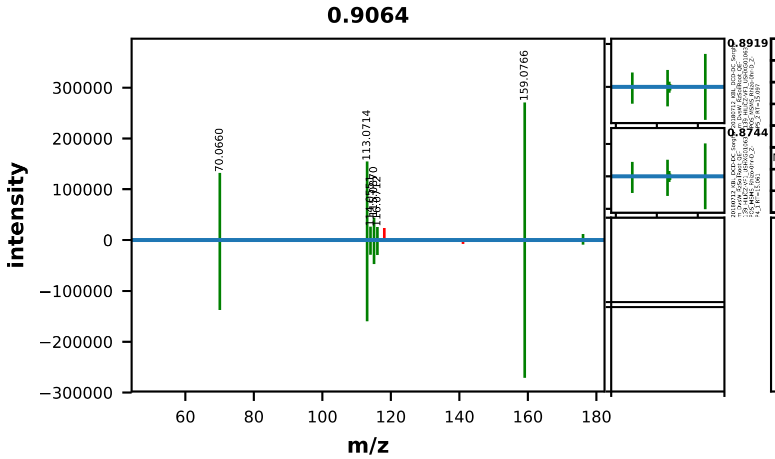


| | EMA Compound Info |
|---|---|
| Name: | L-citrulline |
| Label: | |
| Formula: | C6H13N3O3 |
| Polarity: | positive |
| Monoisotopic Mass: | 175.095691276 |
| Theoretical M/Z: | 176.1029913 |
| Adduct: | |

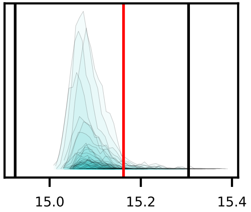


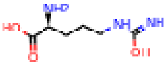

20180712_KBL_DCD-DC_Sorghum_DvsW_RzSoilRoot_QE-139_HILICZ-VF1_USHXG01063_POS_MSMS_Rhizo-0hr-D_Z-P6_3

0038_citrulline_positive_176p1030_15p16
Measured M/Z = 176.1031, 0.4392 ppm difference
Expected Elution of 15.16 minutes, 15.08 min actual

MSMS Scan at 15.103 minutes

Matching M/Zs above 1E-3*max: 70.066, 86.061, 97.076, 113.071, 114.055, 115.087, 116.071, 159.077, 176.103



All Matching M/Zs: 70.066, 86.061, 97.076, 113.071, 114.055, 115.087, 116.071, 159.077, 176.103

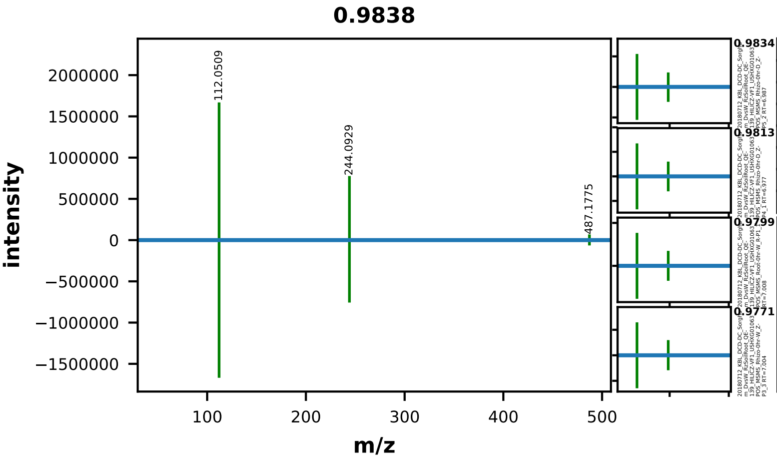


| | EMA Compound Info |
|---|---|
| Name: | cytidine |
| Label: | |
| Formula: | C9H13N3O5 |
| Polarity: | positive |
| Monoisotopic Mass: | 243.085520516 |
| Theoretical M/Z: | 487.178341 |
| Adduct: | |

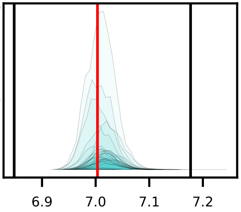


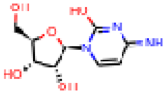

20180712_KBL_DCD-DC_Sorghum_DvsW_RzSoilRoot_QE-139_HILICZ-VF1_USHXG01063_POS_MSMS_Rhizo-0hr-W_Z-P2_2

0041_cytidine_positive_487p1783_7p00
Measured M/Z = 487.1787, 0.7240 ppm difference
Expected Elution of 7.00 minutes, 7.02 min actual

MSMS Scan at 7.007 minutes

Matching M/Zs above 1E-3*max: 112.051, 244.093, 487.178

All Matching M/Zs: 112.051, 244.093, 487.178

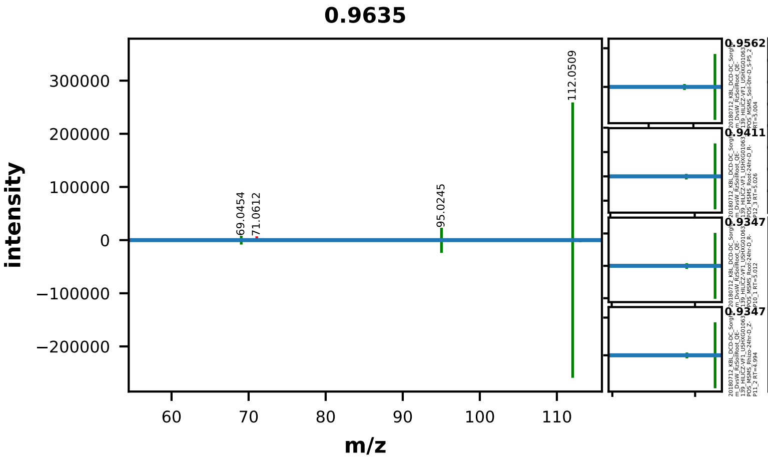


| | EMA Compound Info |
|---|---|
| Name: | cytosine |
| Label: | |
| Formula: | C4H5N3O |
| Polarity: | positive |
| Monoisotopic Mass: | 111.04326178 |
| Theoretical M/Z: | 112.0505618 |
| Adduct: | |

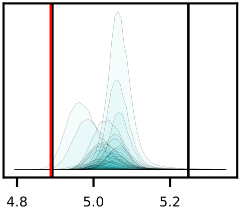


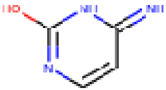


20180712_KBL_DCD-DC_Sorghum_DvsW_RzSoilRoot_QE-139_HILICZ-VF1_USHXG01063_POS_MSMS_Soil-0hr-D_S-P4_1

0043_cytosine_positive_112p0506_4p89
Measured M/Z = 112.0509, 2.7379 ppm difference
Expected Elution of 4.89 minutes, 5.01 min actual

MSMS Scan at 4.992 minutes

Matching M/Zs above 1E-3*max: 69.045, 95.025, 112.051

All Matching M/Zs: 69.045, 95.025, 112.051

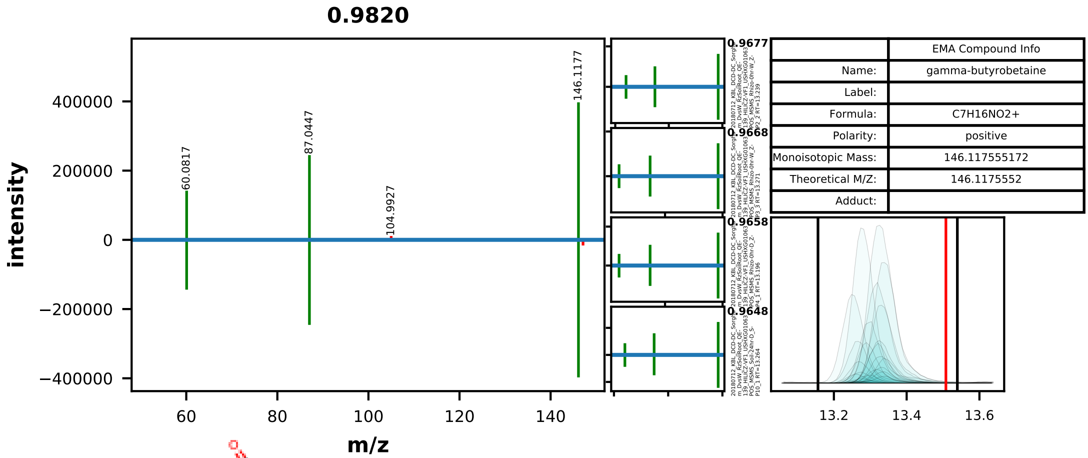


| | EMA Compound Info |
|---|---|
| Name: | gamma-butyrobetaine |
| Label: | |
| Formula: | C7H16NO2+ |
| Polarity: | positive |
| Monoisotopic Mass: | 146.117555172 |
| Theoretical M/Z: | 146.1175552 |
| Adduct: | |

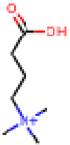


20180712_KBL_DCD-DC_Sorghum_DvsW_RzSoilRoot_QE-139_HILICZ-VF1_USHXG01063_POS_MSMS_Rhizo-24hr-W_Z-P9_3

0044_deoxycarnitine_positive_146p1176_13p51
Measured M/Z = 146.1176, 0.3159 ppm difference
Expected Elution of 13.51 minutes, 13.31 min actual

MSMS Scan at 13.307 minutes

Matching M/Zs above 1E-3*max: 60.082, 87.045, 146.118

All Matching M/Zs: 60.082, 87.045, 146.118

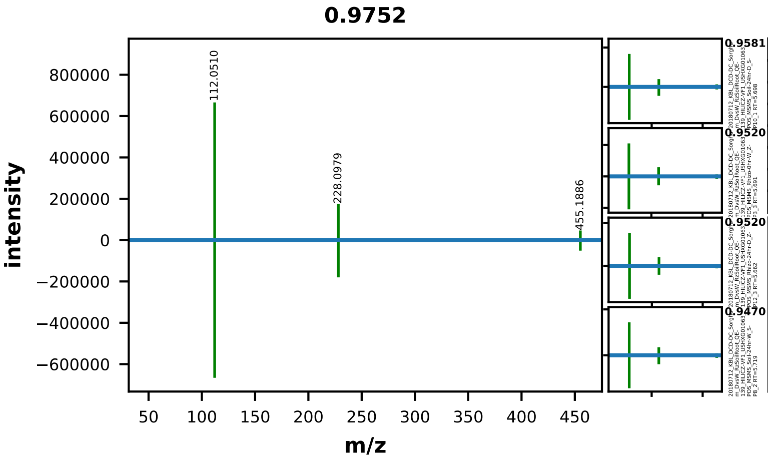


| | EMA Compound Info |
|---|---|
| Name: | 2'-deoxycytidine |
| Label: | |
| Formula: | C9H13N3O4 |
| Polarity: | positive |
| Monoisotopic Mass: | 227.090605896 |
| Theoretical M/Z: | 455.1885118 |
| Adduct: | |

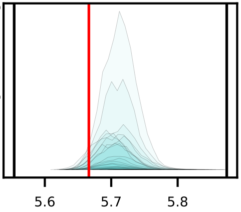


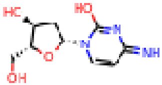


20180712_KBL_DCD-DC_Sorghum_DvsW_RzSoilRoot_QE-139_HILICZ-VF1_USHXG01063_POS_MSMS_Soil-0hr-W_S-P3_3

0045_deoxycytidine_positive_455p1885_5p67
Measured M/Z = 455.1892, 1.6000 ppm difference
Expected Elution of 5.67 minutes, 5.72 min actual

MSMS Scan at 5.723 minutes

Matching M/Zs above 1E-3*max: 112.051, 228.098, 455.189

All Matching M/Zs: 112.051, 228.098, 455.189

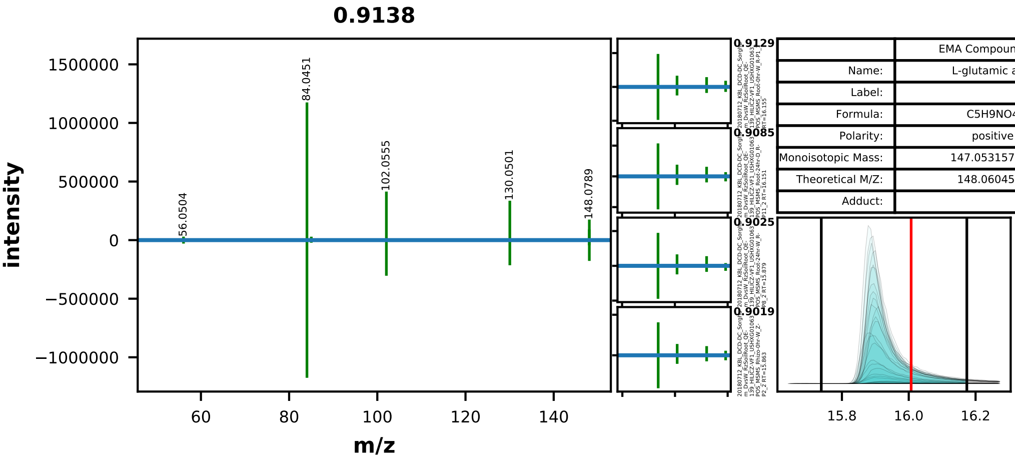


| | EMA Compound Info |
|---|---|
| Name: | L-glutamic acid |
| Label: | |
| Formula: | C5H9NO4 |
| Polarity: | positive |
| Monoisotopic Mass: | 147.053157768 |
| Theoretical M/Z: | 148.0604578 |
| Adduct: | |

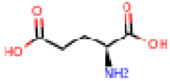

20180712_KBL_DCD-DC_Sorghum_DvsW_RzSoilRoot_QE-139_HILICZ-VF1_USHXG01063_POS_MSMS_Root-0hr-D_R-P4_1

0051_glutamic_acid_positive_148p0605_16p01
Measured M/Z = 148.0606, 0.7197 ppm difference
Expected Elution of 16.01 minutes, 15.88 min actual

MSMS Scan at 16.143 minutes

Matching M/Zs above 1E-3*max: 56.050, 84.045, 85.029, 102.055, 124.043, 130.050, 148.061, 148.079

All Matching M/Zs: 56.050, 84.045, 85.029, 102.055, 124.043, 130.050, 148.061, 148.079

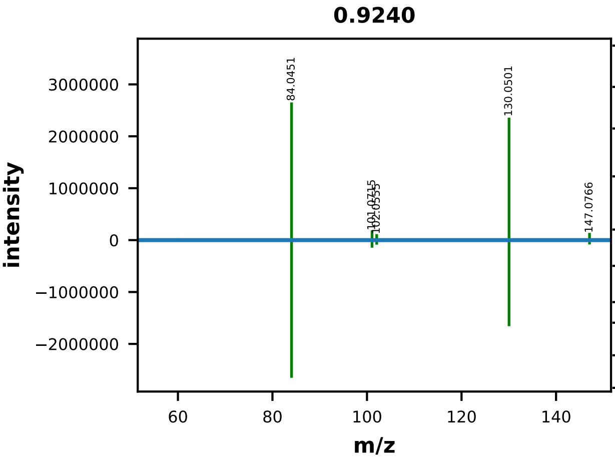


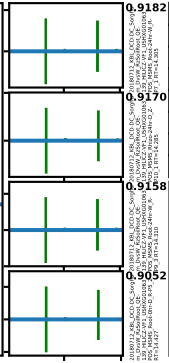


| | EMA Compound Info |
|---|---|
| Name: | L-glutamine |
| Label: | |
| Formula: | C5H10N2O3 |
| Polarity: | positive |
| Monoisotopic Mass: | 146.06914218 |
| Theoretical M/Z: | 147.0764422 |
| Adduct: | |

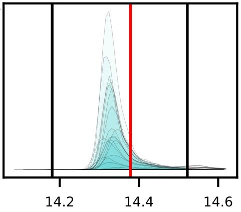


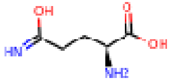


20180712_KBL_DCD-DC_Sorghum_DvsW_RzSoilRoot_QE-139_HILICZ-VF1_USHXG01063_POS_MSMS_Root-0hr-D_R-P4_1

0052_glutamine_positive_147p0764_14p38
Measured M/Z = 147.0765, 0.1311 ppm difference
Expected Elution of 14.38 minutes, 14.32 min actual

MSMS Scan at 14.428 minutes

Matching M/Zs above 1E-3*max: 56.050, 84.045, 85.029, 101.071, 102.056, 123.965, 130.050, 147.077

All Matching M/Zs: 56.050, 84.045, 85.029, 101.071, 102.056, 123.965, 130.050, 147.077

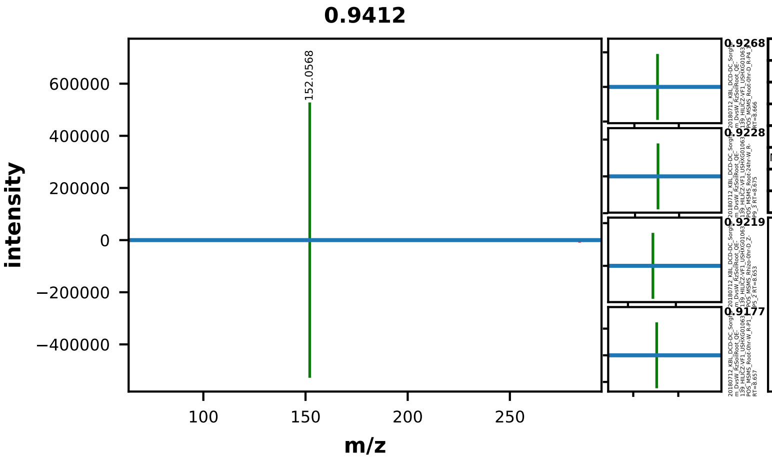


| | EMA Compound Info |
|---|---|
| Name: | guanosine |
| Label: | |
| Formula: | C10H13N5O5 |
| Polarity: | positive |
| Monoisotopic Mass: | 283.091668516 |
| Theoretical M/Z: | 284.0989685 |
| Adduct: | |

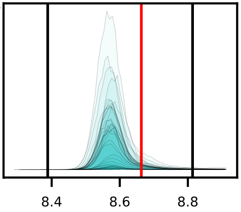

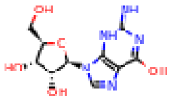

20180712_KBL_DCD-DC_Sorghum_DvsW_RzSoilRoot_QE-139_HILICZ-VF1_USHXG01063_POS_MSMS_Root-24hr-W_R-P8_2

0054_guanosine_positive_284p0990_8p66
Measured M/Z = 284.0993, 1.1089 ppm difference
Expected Elution of 8.66 minutes, 8.57 min actual

MSMS Scan at 8.677 minutes

Matching M/Zs above 1E-3*max: 152.057, 153.040

All Matching M/Zs: 152.057, 153.040

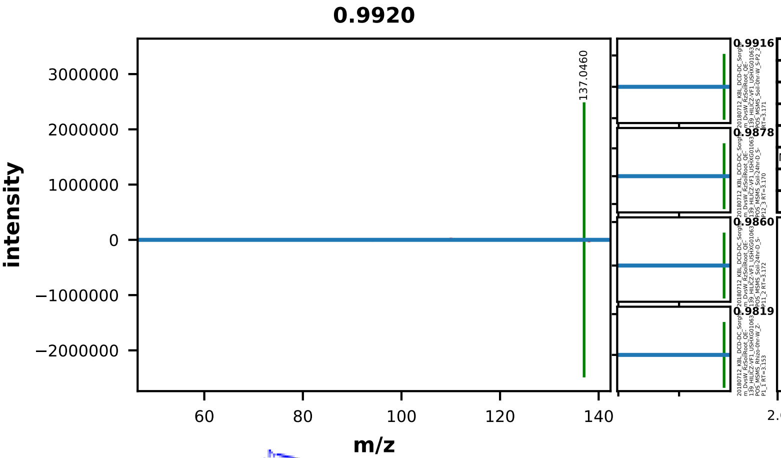


| | EMA Compound Info |
|---|---|
| Name: | hypoxanthine |
| Label: | |
| Formula: | C5H4N4O |
| Polarity: | positive |
| Monoisotopic Mass: | 136.038510748 |
| Theoretical M/Z: | 137.0458107 |
| Adduct: | |

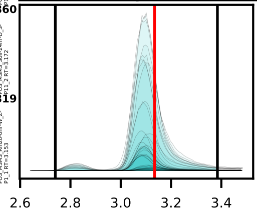


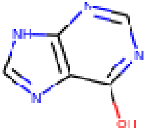


20180712_KBL_DCD-DC_Sorghum_DvsW_RzSoilRoot_QE-139_HILICZ-VF1_USHXG01063_POS_MSMS_Soil-24hr-W_S-P9_3

0059_hypoxanthine_positive_137p0458_3p13
Measured M/Z = 137.0460, 1.2405 ppm difference
Expected Elution of 3.13 minutes, 3.10 min actual

MSMS Scan at 3.171 minutes

Matching M/Zs above 1E-3*max: 94.041, 137.046

All Matching M/Zs: 94.041, 137.046

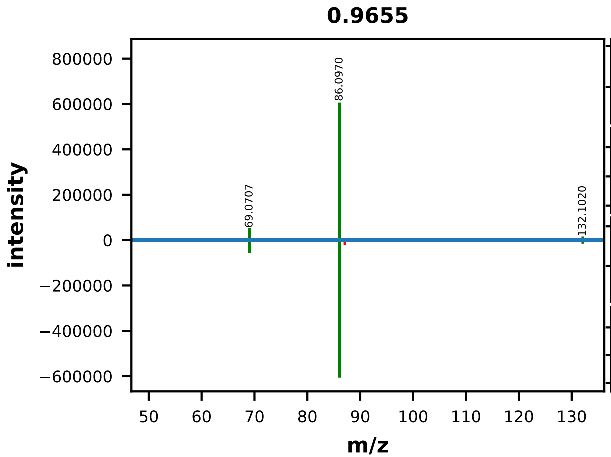


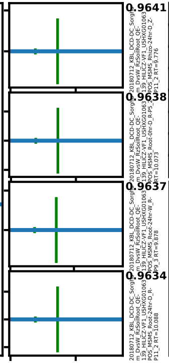


| | EMA Compound Info |
|---|---|
| Name: | l-isoleucine |
| Label: | |
| Formula: | C6H13NO2 |
| Polarity: | positive |
| Monoisotopic Mass: | 131.094628656 |
| Theoretical M/Z: | 132.1019287 |
| Adduct: | |

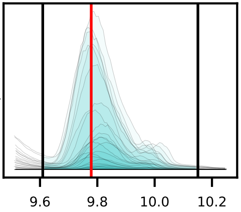


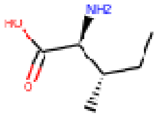


20180712_KBL_DCD-DC_Sorghum_DvsW_RzSoilRoot_QE-139_HILICZ-VF1_USHXG01063_POS_MSMS_Rhizo-24hr-W_Z-P7_1

0063_isoleucine_positive_132p1019_9p78
Measured M/Z = 132.1020, 0.1852 ppm difference
Expected Elution of 9.78 minutes, 9.81 min actual

MSMS Scan at 9.979 minutes

Matching M/Zs above 1E-3*max: 69.071, 86.097, 132.102

All Matching M/Zs: 69.071, 86.097, 132.102

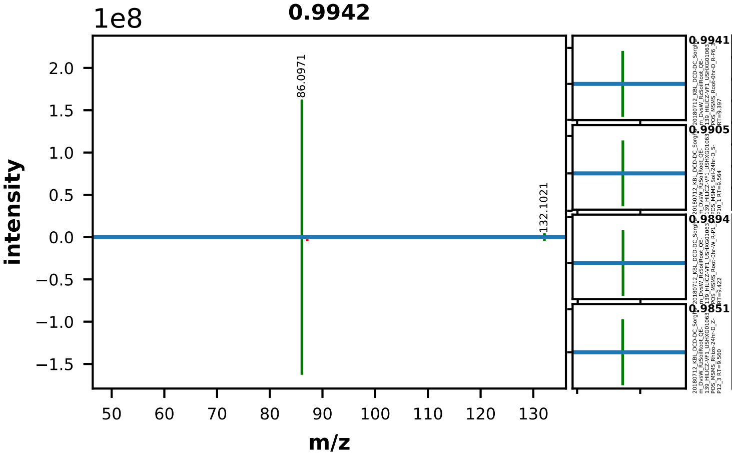


| | EMA Compound Info |
|---|---|
| Name: | DL-Leucine |
| Label: | |
| Formula: | C6H13NO2 |
| Polarity: | positive |
| Monoisotopic Mass: | 131.094628656 |
| Theoretical M/Z: | 132.1019287 |
| Adduct: | |

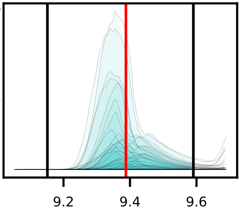


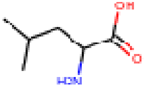


20180712_KBL_DCD-DC_Sorghum_DvsW_RzSoilRoot_QE-139_HILICZ-VF1_USHXG01063_POS_MSMS_Root-24hr-D_R-P11_2

0064_leucine_positive_132p1019_9p39
Measured M/Z = 132.1021, 1.6659 ppm difference
Expected Elution of 9.39 minutes, 9.36 min actual

MSMS Scan at 9.384 minutes

Matching M/Zs above 1E-3*max: 86.097, 132.102

All Matching M/Zs: 86.097, 132.102

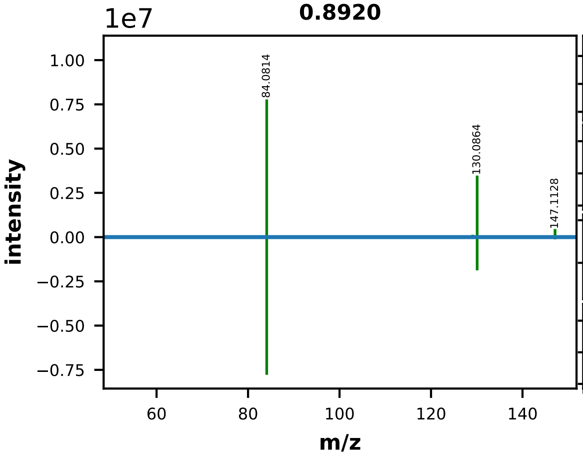


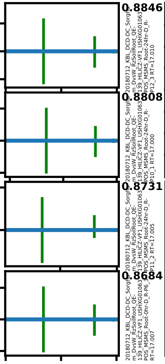


| | EMA Compound Info |
|---|---|
| Name: | L-lysine |
| Label: | |
| Formula: | C6H14N2O2 |
| Polarity: | positive |
| Monoisotopic Mass: | 146.105527688 |
| Theoretical M/Z: | 147.1128277 |
| Adduct: | |

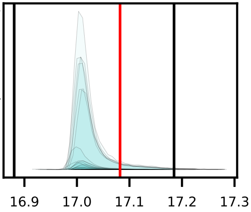

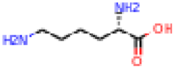


20180712_KBL_DCD-DC_Sorghum_DvsW_RzSoilRoot_QE-139_HILICZ-VF1_USHXG01063_POS_MSMS_Root-0hr-D_R-P5_2

0065_lysine_positive_147p1128_17p08
Measured M/Z = 147.1128, 0.2047 ppm difference
Expected Elution of 17.08 minutes, 17.01 min actual

MSMS Scan at 17.008 minutes

Matching M/Zs above 1E-3*max: 56.050, 67.055, 74.024, 84.081, 85.066, 102.092, 112.076, 123.965, 129.102, 130.086, 147.113

All Matching M/Zs: 56.050, 67.055, 74.024, 84.081, 85.066, 102.092, 112.076, 123.965, 129.102, 130.086, 147.113

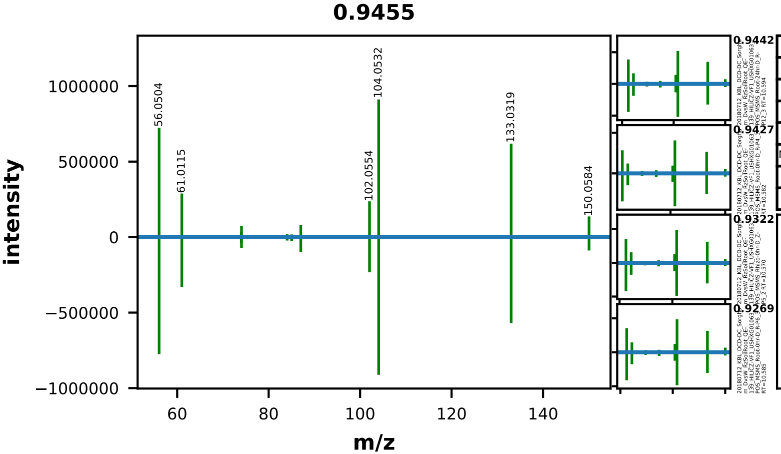


| | EMA Compound Info |
|---|---|
| Name: | L-methionine |
| Label: | |
| Formula: | C5H11NO2S |
| Polarity: | positive |
| Monoisotopic Mass: | 149.051049592 |
| Theoretical M/Z: | 150.0583496 |
| Adduct: | |

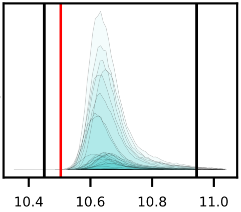


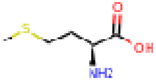


20180712_KBL_DCD-DC_Sorghum_DvsW_RzSoilRoot_QE-139_HILICZ-VF1_USHXG01063_POS_MSMS_Root-0hr-D_R-P5_2

0068_methionine_positive_150p0583_10p50
Measured M/Z = 150.0584, 0.5069 ppm difference
Expected Elution of 10.50 minutes, 10.63 min actual

MSMS Scan at 10.577 minutes

Matching M/Zs above 1E-3*max: 56.050, 61.011, 74.024, 74.061, 75.027, 84.045, 85.029, 87.027, 102.055, 104.053, 105.001, 105.037, 133.032, 150.058

All Matching M/Zs: 56.050, 61.011, 74.024, 74.061, 75.027, 84.045, 85.029, 87.027, 102.055, 104.053, 105.001, 105.037, 133.032, 150.058

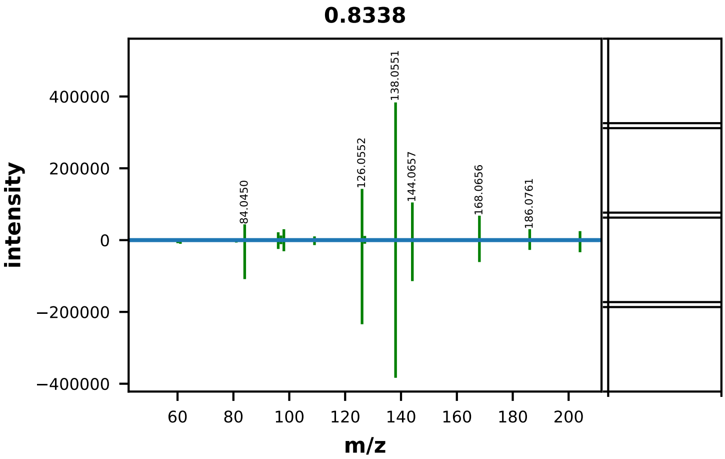


| | EMA Compound Info |
|---|---|
| Name: | N-Acetyl-D-Glucosamine |
| Label: | |
| Formula: | C8H15NO6 |
| Polarity: | positive |
| Monoisotopic Mass: | 221.0899372 |
| Theoretical M/Z: | 204.0866372 |
| Adduct: | |

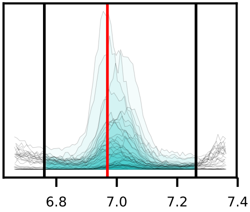


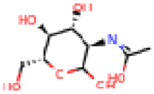


20180712_KBL_DCD-DC_Sorghum_DvsW_RzSoilRoot_QE-139_HILICZ-VF1_USHXG01063_POS_MSMS_Rhizo-0hr-D_Z-P6_3

0071_N-acetyl-glucosamine_positive_204p0866_6p97
Measured M/Z = 204.0867, 0.3428 ppm difference
Expected Elution of 6.97 minutes, 6.96 min actual

MSMS Scan at 6.979 minutes

Matching M/Zs above 1E-3*max: 60.045, 61.029, 69.034, 81.034, 84.045, 96.045, 97.029, 98.061, 108.045, 109.029, 126.055, 127.039, 138.055, 144.066, 168.066, 186.076, 204.087

All Matching M/Zs: 60.045, 61.029, 69.034, 81.034, 84.045, 96.045, 97.029, 98.061, 108.045, 109.029, 126.055, 127.039, 138.055, 144.066, 168.066, 186.076, 204.087

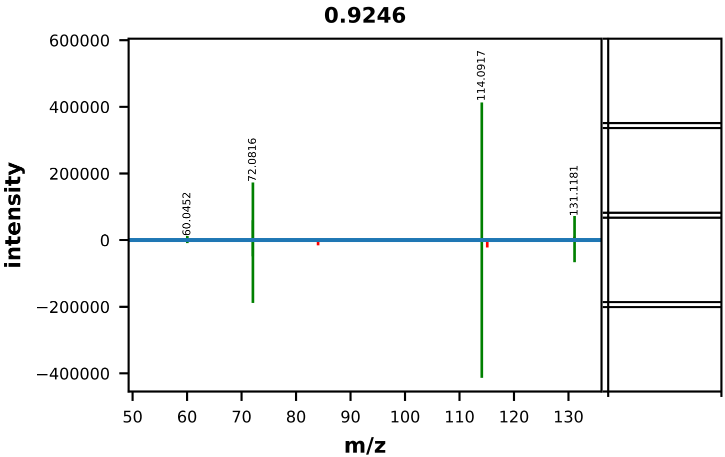


| | EMA Compound Info |
|---|---|
| Name: | N-(4-aminobutyl)acetamide |
| Label: | |
| Formula: | C6H14N2O |
| Polarity: | positive |
| Monoisotopic Mass: | 130.110613068 |
| Theoretical M/Z: | 131.1179131 |
| Adduct: | |

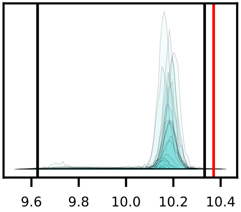


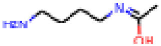

20180712_KBL_DCD-DC_Sorghum_DvsW_RzSoilRoot_QE-139_HILICZ-VF1_USHXG01063_POS_MSMS_Rhizo-0hr-D_Z-P4_1

0073_N-acetylputrescine_positive_131p1179_10p37
Measured M/Z = 131.1180, 0.7995 ppm difference
Expected Elution of 10.37 minutes, 10.16 min actual

MSMS Scan at 10.159 minutes

Matching M/Zs above 1E-3*max: 60.045, 72.045, 72.082, 114.092, 131.118

All Matching M/Zs: 60.045, 72.045, 72.082, 114.092, 131.118

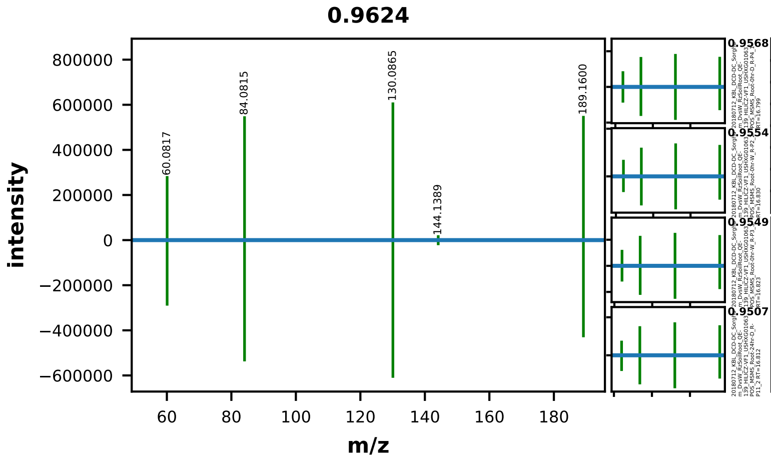


| | EMA Compound Info |
|---|---|
| Name: | Trimethyllysine |
| Label: | |
| Formula: | C9H21N2O2+ |
| Polarity: | positive |
| Monoisotopic Mass: | 189.159754332 |
| Theoretical M/Z: | 189.1597543 |
| Adduct: | |

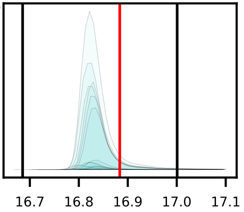


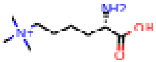


20180712_KBL_DCD-DC_Sorghum_DvsW_RzSoilRoot_QE-139_HILICZ-VF1_USHXG01063_POS_MSMS_Root-24hr-W_R-P9_3

0076_N-trimethyllysine_positive_189p1598_16p88
Measured M/Z = 189.1599, 0.7605 ppm difference
Expected Elution of 16.88 minutes, 16.83 min actual

MSMS Scan at 16.831 minutes

Matching M/Zs above 1E-3*max: 60.082, 84.082, 130.087, 144.139, 189.160

All Matching M/Zs: 60.082, 84.082, 130.087, 144.139, 189.160

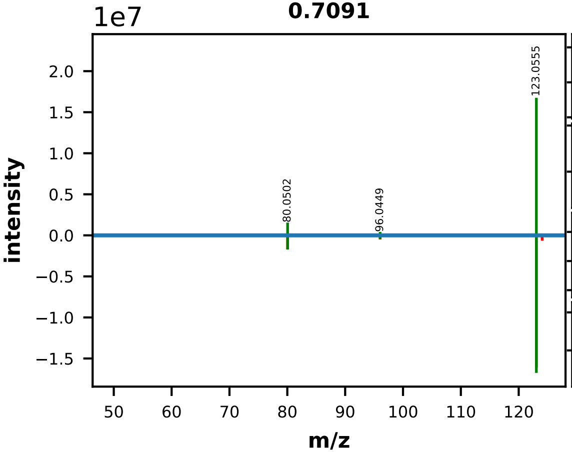


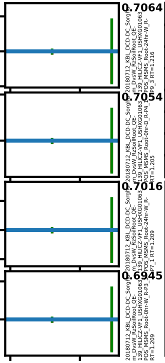


| | EMA Compound Info |
|---|---|
| Name: | nicotinamide |
| Label: | |
| Formula: | C6H6N2O |
| Polarity: | positive |
| Monoisotopic Mass: | 122.048012812 |
| Theoretical M/Z: | 123.0553128 |
| Adduct: | |

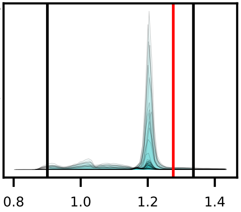


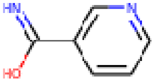


20180712_KBL_DCD-DC_Sorghum_DvsW_RzSoilRoot_QE-139_HILICZ-VF1_USHXG01063_POS_MSMS_Root-24hr-D_R-P10_1

0077_nicotinamide_positive_123p0553_1p28
Measured M/Z = 123.0555, 1.8893 ppm difference
Expected Elution of 1.28 minutes, 1.20 min actual

MSMS Scan at 1.208 minutes

Matching M/Zs above 1E-3*max: 53.039, 80.050, 96.045, 106.029, 123.056, 124.039

All Matching M/Zs: 53.039, 80.050, 96.045, 106.029, 123.056, 124.039

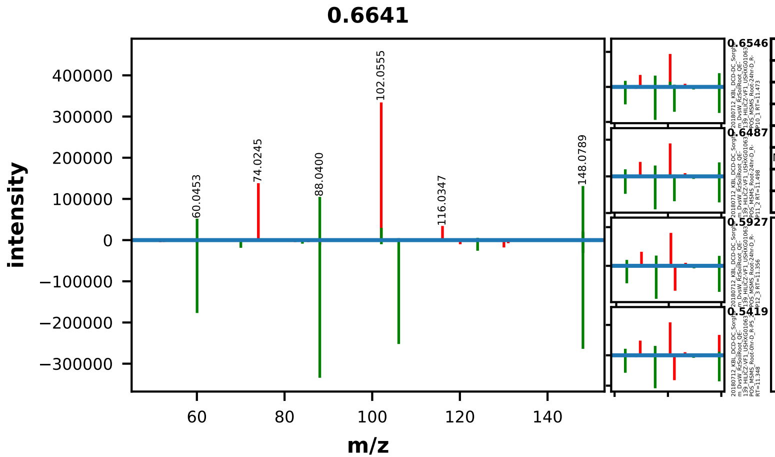


| EMA Compound Info | |
|---|---|
| Name: | O-acetyl-L-serine |
| Label: | |
| Formula: | C5H9NO4 |
| Polarity: | positive |
| Monoisotopic Mass: | 147.053157768 |
| Theoretical M/Z: | 148.0604578 |
| Adduct: | |

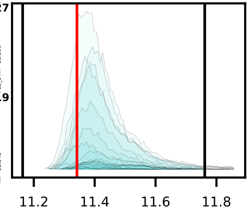


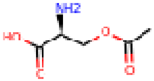


20180712_KBL_DCD-DC_Sorghum_DvsW_RzSoilRoot_QE-139_HILICZ-VF1_USHXG01063_POS_MSMS_Root-0hr-D_R-P4_1

0078_o-acetyl-serine_positive_148p0605_11p34
Measured M/Z = 148.0605, 0.5617 ppm difference
Expected Elution of 11.34 minutes, 11.37 min actual

MSMS Scan at 11.455 minutes

Matching M/Zs above 1E-3*max: 60.043, 60.045, 70.029, 84.081, 88.040, 102.092, 106.050, 124.043, 148.079, 148.096

All Matching M/Zs: 60.043, 60.045, 70.029, 84.081, 88.040, 102.092, 106.050, 124.043, 148.079, 148.096

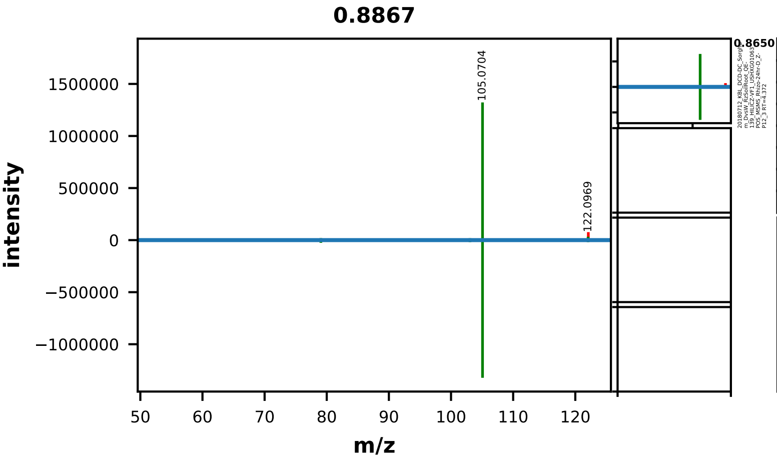


| | EMA Compound Info |
|---|---|
| Name: | Phenethylamine |
| Label: | |
| Formula: | C8H11N |
| Polarity: | positive |
| Monoisotopic Mass: | 121.089149352 |
| Theoretical M/Z: | 122.0964494 |
| Adduct: | |

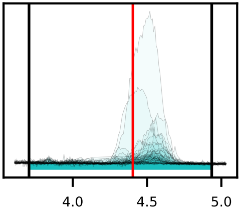


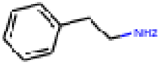


20180712_KBL_DCD-DC_Sorghum_DvsW_RzSoilRoot_QE-139_HILICZ-VF1_USHXG01063_POS_MSMS_Rhizo-0hr-D_Z-P6_3

0081_phenethylamine_positive_122p0964_4p40
Measured M/Z = 122.0968, 2.4797 ppm difference
Expected Elution of 4.40 minutes, 4.52 min actual

MSMS Scan at 4.440 minutes

Matching M/Zs above 1E-3*max: 79.055, 103.055, 105.070, 122.060

All Matching M/Zs: 79.055, 103.055, 105.070, 122.060

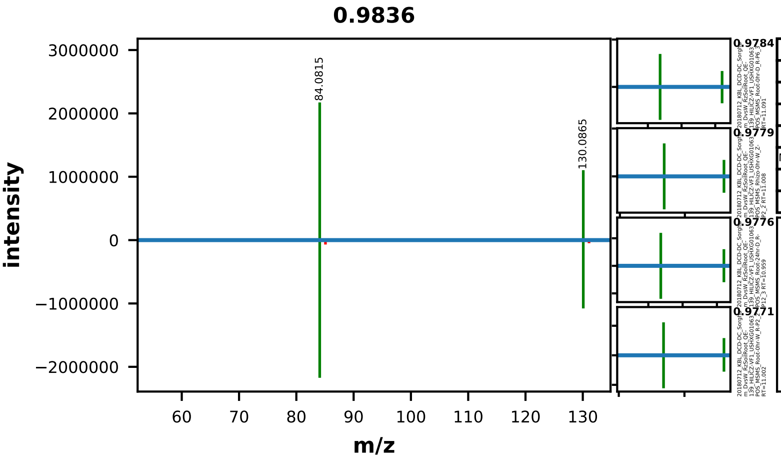


| | EMA Compound Info |
|---|---|
| Name: | l-Pipecolic acid |
| Label: | |
| Formula: | C6H11NO2 |
| Polarity: | positive |
| Monoisotopic Mass: | 129.078978592 |
| Theoretical M/Z: | 130.0862786 |
| Adduct: | |

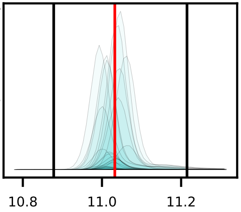


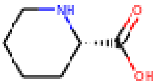


20180712_KBL_DCD-DC_Sorghum_DvsW_RzSoilRoot_QE-139_HILICZ-VF1_USHXG01063_POS_MSMS_Rhizo-24hr-D_Z-P10_1

0082_pipecolic_acid_positive_130p0863_11p03
Measured M/Z = 130.0864, 1.0972 ppm difference
Expected Elution of 11.03 minutes, 11.06 min actual

MSMS Scan at 11.009 minutes

Matching M/Zs above 1E-3*max: 56.050, 84.081, 130.086

All Matching M/Zs: 56.050, 84.081, 130.086

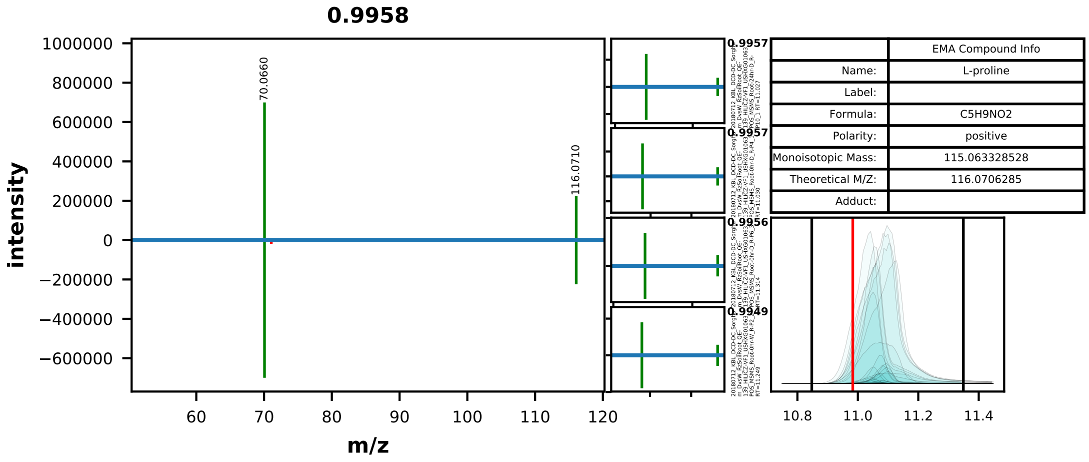


| | EMA Compound Info |
|---|---|
| Name: | L-proline |
| Label: | |
| Formula: | C5H9NO2 |
| Polarity: | positive |
| Monoisotopic Mass: | 115.063328528 |
| Theoretical M/Z: | 116.0706285 |
| Adduct: | |

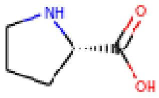


20180712_KBL_DCD-DC_Sorghum_DvsW_RzSoilRoot_QE-139_HILICZ-VF1_USHXG01063_POS_MSMS_Root-24hr-W_R-P9_3

0083_proline_positive_116p0706_10p98
Measured M/Z = 116.0709, 2.2353 ppm difference
Expected Elution of 10.98 minutes, 11.07 min actual

MSMS Scan at 11.276 minutes

Matching M/Zs above 1E-3*max: 70.066, 116.071

All Matching M/Zs: 70.066, 116.071

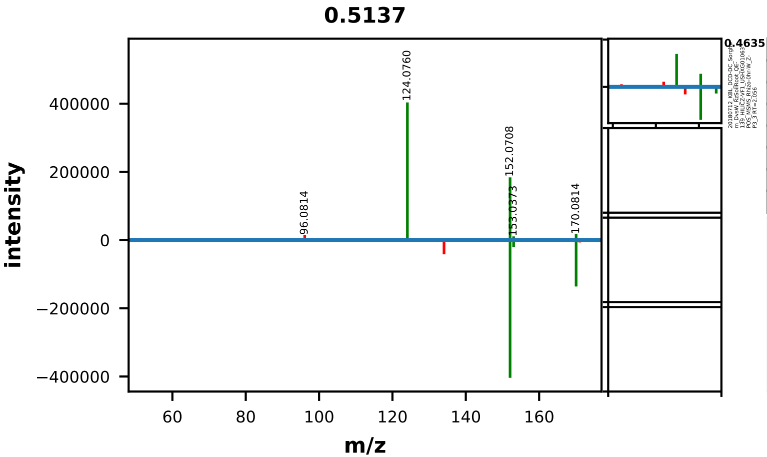


| | EMA Compound Info |
|---|---|
| Name: | pyridoxine |
| Label: | |
| Formula: | C8H11NO3 |
| Polarity: | positive |
| Monoisotopic Mass: | 169.073893212 |
| Theoretical M/Z: | 170.0811932 |
| Adduct: | |

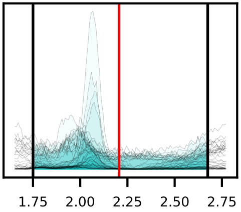


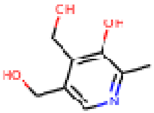


20180712_KBL_DCD-DC_Sorghum_DvsW_RzSoilRoot_QE-139_HILICZ-VF1_USHXG01063_POS_MSMS_Rhizo-24hr-D_Z-P12_3

0086_pyridoxine_positive_170p0812_2p21
Measured M/Z = 170.0813, 0.8824 ppm difference
Expected Elution of 2.21 minutes, 2.06 min actual

MSMS Scan at 2.074 minutes

Matching M/Zs above 1E-3*max: 124.076, 152.071, 153.037, 170.081

All Matching M/Zs: 124.076, 152.071, 153.037, 170.081

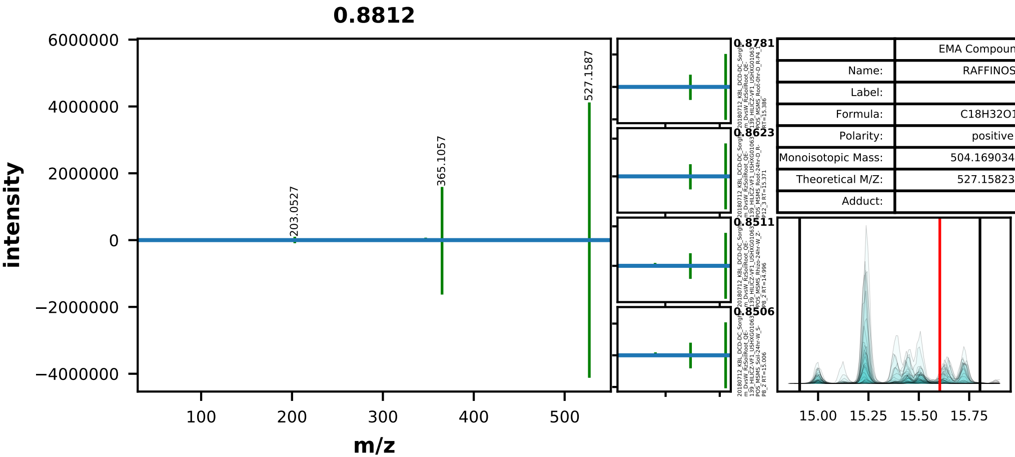


| | EMA Compound Info |
|---|---|
| Name: | RAFFINOSE |
| Label: | |
| Formula: | C18H32O16 |
| Polarity: | positive |
| Monoisotopic Mass: | 504.169034944 |
| Theoretical M/Z: | 527.1582349 |
| Adduct: | |

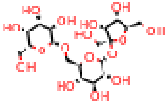

20180712_KBL_DCD-DC_Sorghum_DvsW_RzSoilRoot_QE-139_HILICZ-VF1_USHXG01063_POS_MSMS_Root-24hr-D_R-P10_1

0089_raffinose_positive_527p1582_15p60
Measured M/Z = 527.1591, 1.6809 ppm difference
Expected Elution of 15.60 minutes, 15.24 min actual

MSMS Scan at 15.384 minutes

Matching M/Zs above 1E-3*max: 185.042, 203.053, 305.084, 347.096, 365.106, 527.159

All Matching M/Zs: 185.042, 203.053, 305.084, 347.096, 365.106, 527.159

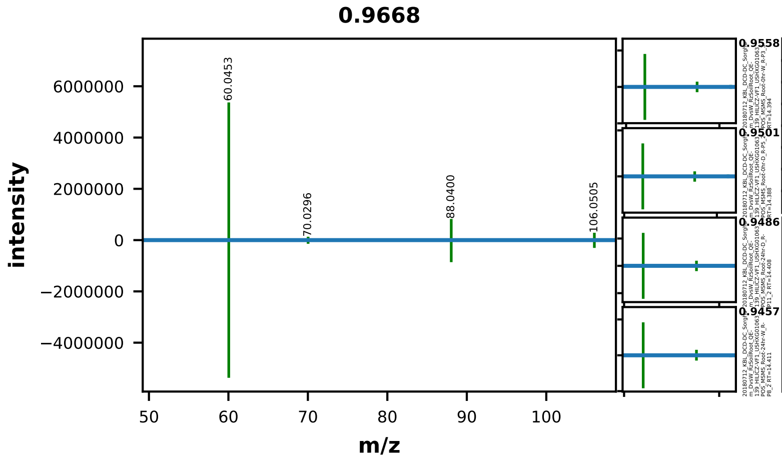


| | EMA Compound Info |
|---|---|
| Name: | L-serine |
| Label: | |
| Formula: | C3H7NO3 |
| Polarity: | positive |
| Monoisotopic Mass: | 105.042593084 |
| Theoretical M/Z: | 106.0498931 |
| Adduct: | |

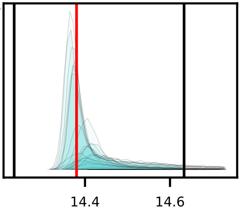


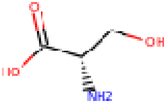


20180712_KBL_DCD-DC_Sorghum_DvsW_RzSoilRoot_QE-139_HILICZ-VF1_USHXG01063_POS_MSMS_Root-24hr-D_R-P10_1

0092_serine_positive_106p0499_14p38
Measured M/Z = 106.0503, 3.7675 ppm difference
Expected Elution of 14.38 minutes, 14.36 min actual

MSMS Scan at 14.394 minutes

Matching M/Zs above 1E-3*max: 60.045, 70.030, 88.040, 106.050

All Matching M/Zs: 60.045, 70.030, 88.040, 106.050

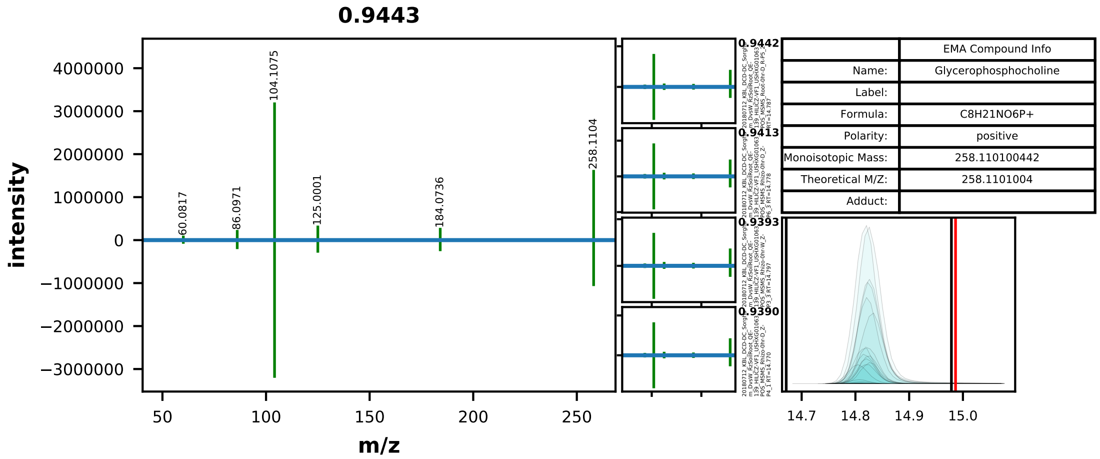


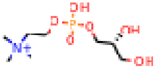

20180712_KBL_DCD-DC_Sorghum_DvsW_RzSoilRoot_QE-139_HILICZ-VF1_USHXG01063_POS_MSMS_Root-24hr-W_R-P7_1

0094_sn-glycero-3-phosphocholine_positive_258p1101_14p99
Measured M/Z = 258.1104, 1.0216 ppm difference
Expected Elution of 14.99 minutes, 14.82 min actual

MSMS Scan at 14.785 minutes

Matching M/Zs above 1E-3*max: 60.082, 86.097, 104.108, 125.000, 184.074, 258.110

All Matching M/Zs: 60.082, 86.097, 104.108, 125.000, 184.074, 258.110

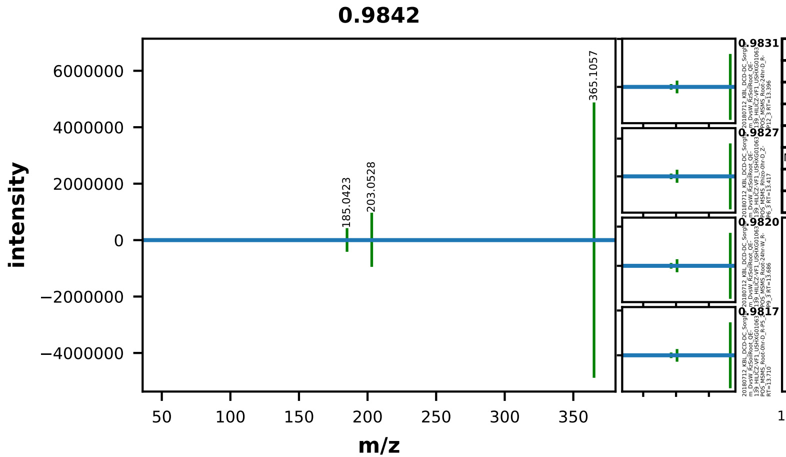


| | EMA Compound Info |
|---|---|
| Name: | sucrose |
| Label: | |
| Formula: | C12H22O11 |
| Polarity: | positive |
| Monoisotopic Mass: | 342.116211524 |
| Theoretical M/Z: | 365.1054115 |
| Adduct: | |

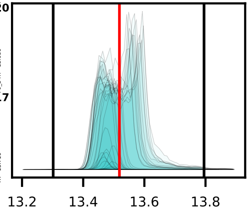


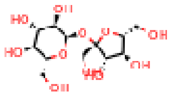

20180712_KBL_DCD-DC_Sorghum_DvsW_RzSoilRoot_QE-139_HILICZ-VF1_USHXG01063_POS_MSMS_Rhizo-24hr-D_Z-P11_2

0096_sucrose_positive_365p1054_13p52
Measured M/Z = 365.1059, 1.2048 ppm difference
Expected Elution of 13.52 minutes, 13.48 min actual

MSMS Scan at 13.450 minutes

Matching M/Zs above 1E-3*max: 185.042, 203.053, 365.106

All Matching M/Zs: 185.042, 203.053, 365.106

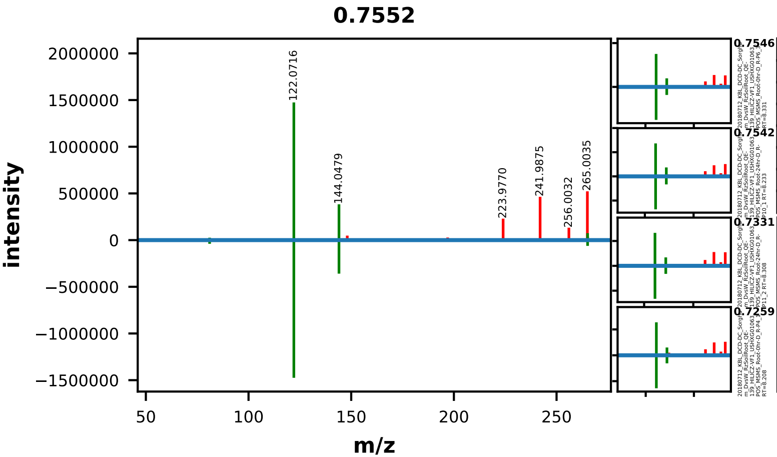


| | EMA Compound Info |
|---|---|
| Name: | thiamine |
| Label: | |
| Formula: | C12H17N4OS+ |
| Polarity: | positive |
| Monoisotopic Mass: | 265.111758584 |
| Theoretical M/Z: | 265.1117586 |
| Adduct: | |

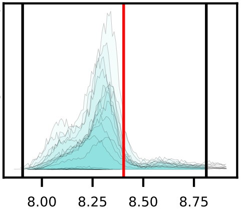


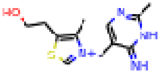

20180712_KBL_DCD-DC_Sorghum_DvsW_RzSoilRoot_QE-139_HILICZ-VF1_USHXG01063_POS_MSMS_Root-24hr-D_R-P12_3

0098_thiamine_positive_265p1118_8p40
Measured M/Z = 265.1120, 1.0779 ppm difference
Expected Elution of 8.40 minutes, 8.34 min actual

MSMS Scan at 8.254 minutes

Matching M/Zs above 1E-3*max: 81.046, 122.072, 144.048, 265.112

All Matching M/Zs: 81.046, 122.072, 144.048, 265.112

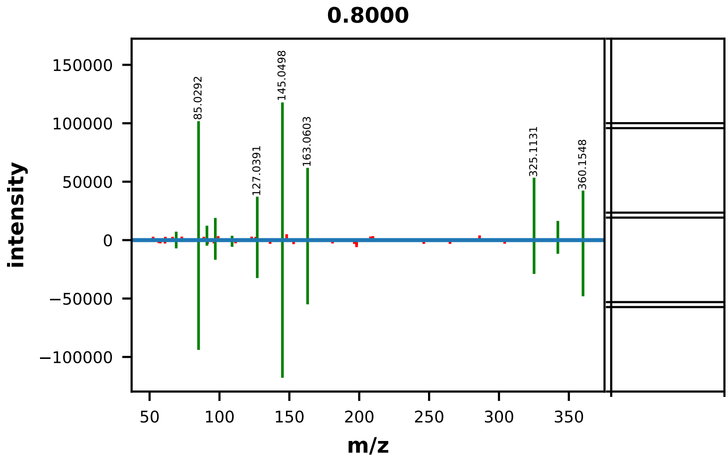


| | EMA Compound Info |
|---|---|
| Name: | trehalose |
| Label: | |
| Formula: | C12H22O11 |
| Polarity: | positive |
| Monoisotopic Mass: | 342.116211524 |
| Theoretical M/Z: | 360.1500115 |
| Adduct: | |

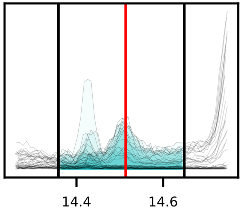


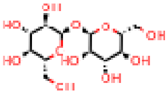

20180712_KBL_DCD-DC_Sorghum_DvsW_RzSoilRoot_QE-139_HILICZ-VF1_USHXG01063_POS_MSMS_Root-0hr-W_R-P1_1

0102_trehalose_positive_360p1500_14p51
Measured M/Z = 360.1505, 1.3960 ppm difference
Expected Elution of 14.51 minutes, 14.51 min actual

MSMS Scan at 14.530 minutes

Matching M/Zs above 1E-3*max: 69.034, 85.029, 91.040, 97.029, 109.029, 127.039, 145.050, 163.060, 325.113, 342.140, 360.155

All Matching M/Zs: 69.034, 85.029, 91.040, 97.029, 109.029, 127.039, 145.050, 163.060, 325.113, 342.140, 360.155

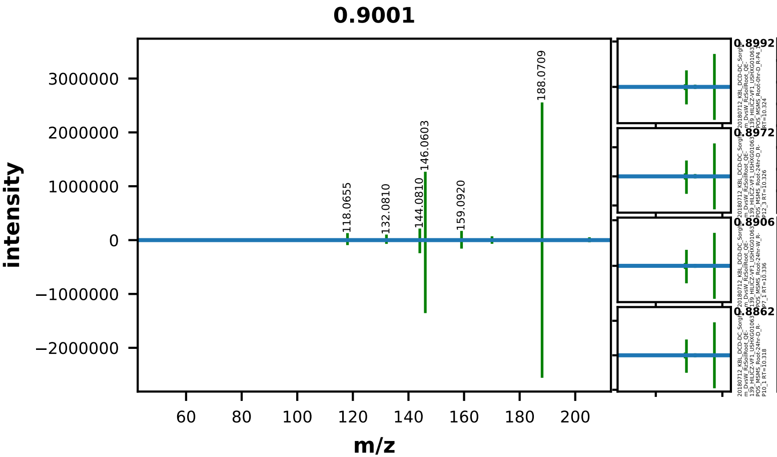


| | EMA Compound Info |
|---|---|
| Name: | D-Tryptophan |
| Label: | |
| Formula: | C11H12N2O2 |
| Polarity: | positive |
| Monoisotopic Mass: | 204.089877624 |
| Theoretical M/Z: | 205.0971776 |
| Adduct: | |

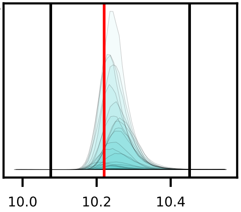


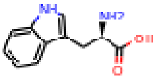


20180712_KBL_DCD-DC_Sorghum_DvsW_RzSoilRoot_QE-139_HILICZ-VF1_USHXG01063_POS_MSMS_Root-24hr-D_R-P11_2

0103_tryptophan_positive_205p0972_10p22
Measured M/Z = 205.0974, 1.0597 ppm difference
Expected Elution of 10.22 minutes, 10.24 min actual

MSMS Scan at 10.182 minutes

Matching M/Zs above 1E-3*max: 74.025, 118.066, 130.066, 132.081, 142.066, 144.081, 146.060, 159.092, 160.076, 170.060, 188.071, 205.097



All Matching M/Zs: 74.025, 118.066, 130.066, 132.081, 142.066, 144.081, 146.060, 159.092, 160.076, 170.060, 188.071, 205.097

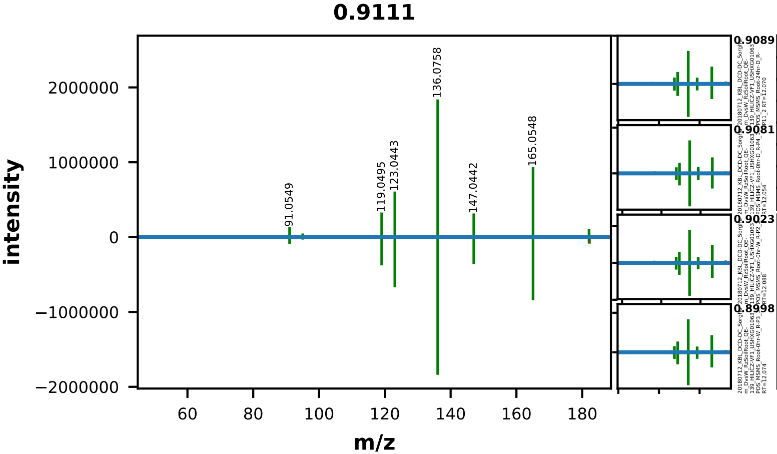


| | EMA Compound Info |
|---|---|
| Name: | L-tyrosine |
| Label: | |
| Formula: | C9H11NO3 |
| Polarity: | positive |
| Monoisotopic Mass: | 181.073893212 |
| Theoretical M/Z: | 182.0811932 |
| Adduct: | |

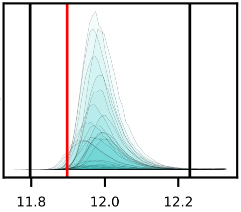


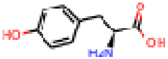


20180712_KBL_DCD-DC_Sorghum_DvsW_RzSoilRoot_QE-139_HILICZ-VF1_USHXG01063_POS_MSMS_Root-24hr-D_R-P10_1

0104_tyrosine_positive_182p0812_11p90
Measured M/Z = 182.0813, 0.7636 ppm difference
Expected Elution of 11.90 minutes, 11.98 min actual

MSMS Scan at 12.057 minutes

Matching M/Zs above 1E-3*max: 91.055, 95.050, 119.049, 123.044, 136.076, 147.044, 164.073, 165.055, 182.081, 182.098



All Matching M/Zs: 91.055, 95.050, 119.049, 123.044, 136.076, 147.044, 164.073, 165.055, 182.081, 182.098

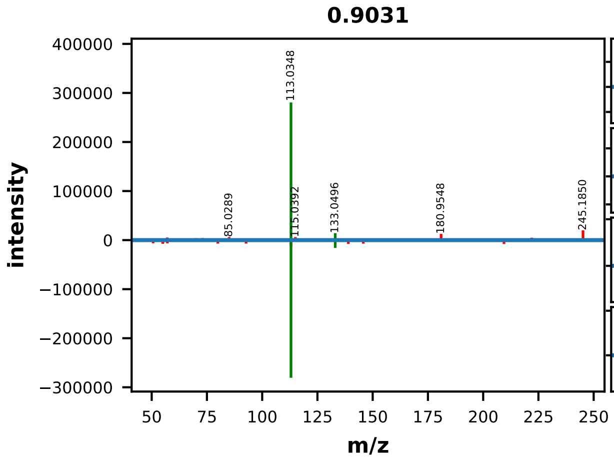


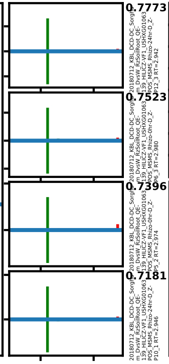


| | EMA Compound Info |
|---|---|
| Name: | uridine |
| Label: | |
| Formula: | C9H12N2O6 |
| Polarity: | positive |
| Monoisotopic Mass: | 244.069536104 |
| Theoretical M/Z: | 245.0768361 |
| Adduct: | |

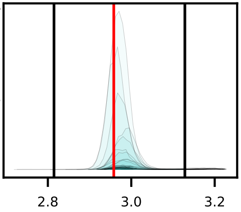


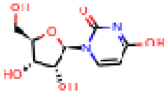

20180712_KBL_DCD-DC_Sorghum_DvsW_RzSoilRoot_QE-139_HILICZ-VF1_USHXG01063_POS_MSMS_Rhizo-24hr-W_Z-P7_1

0105_uridine_positive_245p0768_2p96
Measured M/Z = 245.0768, 0.0169 ppm difference
Expected Elution of 2.96 minutes, 2.98 min actual

MSMS Scan at 2.978 minutes

Matching M/Zs above 1E-3*max: 113.035, 133.050

All Matching M/Zs: 113.035, 133.050

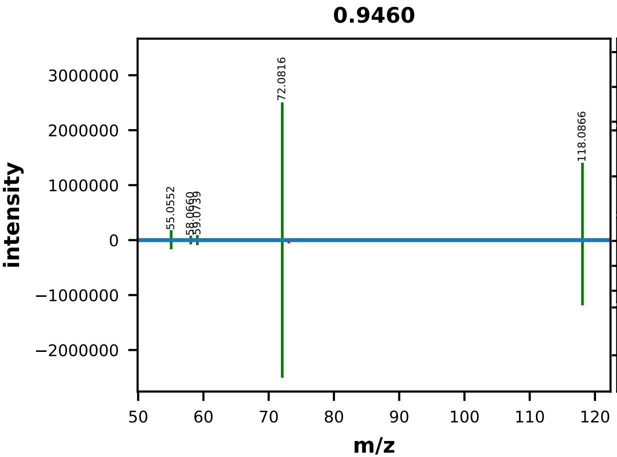


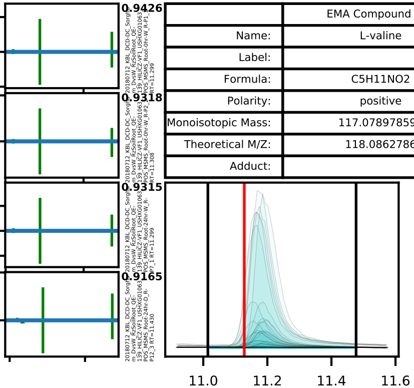


| | EMA Compound Info |
|---|---|
| Name: | L-valine |
| Label: | |
| Formula: | C5H11NO2 |
| Polarity: | positive |
| Monoisotopic Mass: | 117.078978592 |
| Theoretical M/Z: | 118.0862786 |
| Adduct: | |

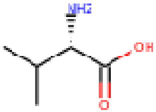


20180712_KBL_DCD-DC_Sorghum_DvsW_RzSoilRoot_QE-139_HILICZ-VF1_USHXG01063_POS_MSMS_Rhizo-24hr-W_Z-P8_2

0108_valine_positive_118p0863_11p13
Measured M/Z = 118.0865, 1.9006 ppm difference
Expected Elution of 11.13 minutes, 11.17 min actual

MSMS Scan at 11.137 minutes

Matching M/Zs above 1E-3*max: 55.055, 58.066, 59.069, 59.074, 72.082, 118.087

All Matching M/Zs: 55.055, 58.066, 59.069, 59.074, 72.082, 118.087

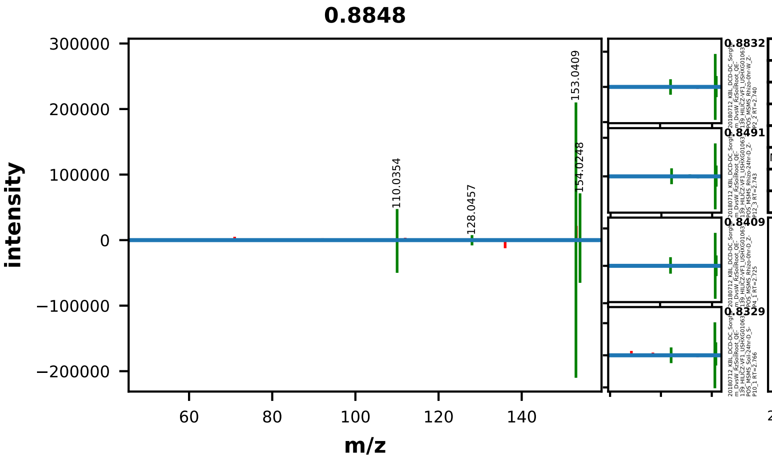


| | EMA Compound Info |
|---|---|
| Name: | xanthine |
| Label: | |
| Formula: | C5H4N4O2 |
| Polarity: | positive |
| Monoisotopic Mass: | 152.033425368 |
| Theoretical M/Z: | 153.0407254 |
| Adduct: | |

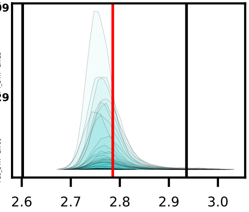

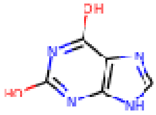


20180712_KBL_DCD-DC_Sorghum_DvsW_RzSoilRoot_QE-139_HILICZ-VF1_USHXG01063_POS_MSMS_Rhizo-0hr-D_Z-P5_2

0109_xanthine_positive_153p0407_2p79
Measured M/Z = 153.0408, 0.5819 ppm difference
Expected Elution of 2.79 minutes, 2.78 min actual

MSMS Scan at 2.759 minutes

Matching M/Zs above 1E-3*max: 110.035, 128.046, 153.041, 154.025

All Matching M/Zs: 110.035, 128.046, 153.041, 154.025

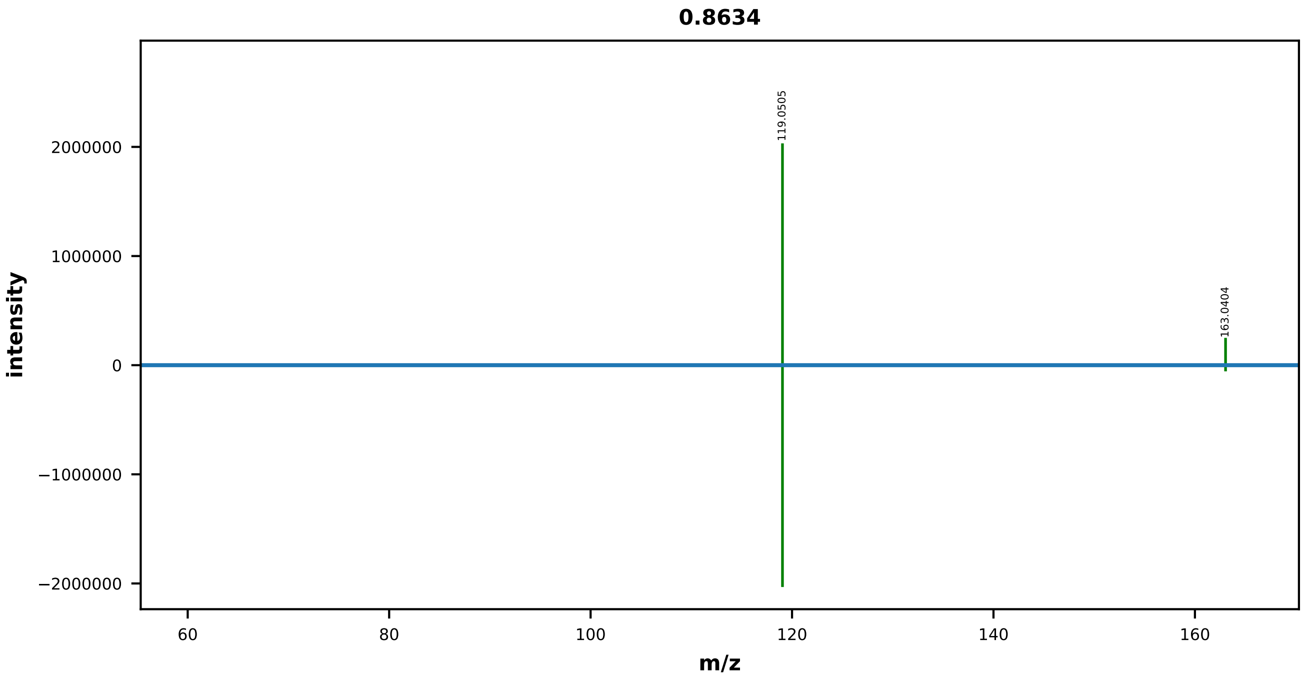


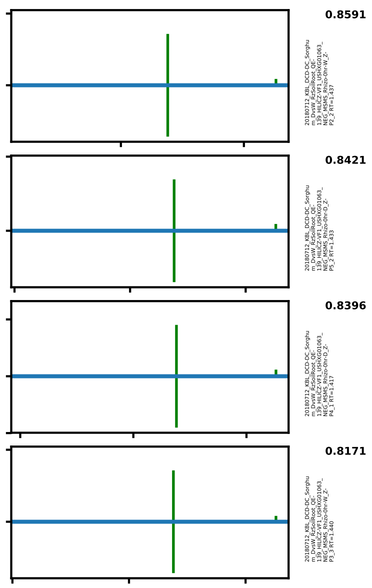


| | EMA Compound Info |
|---|---|
| Name: | 2-Hydroxycinnamic acid |
| Label: | |
| Formula: | C9H8O3 |
| Polarity: | negative |
| Monoisotopic Mass: | 164.047344116 |
| Theoretical M/Z: | 163.0400441 |
| Adduct: | |

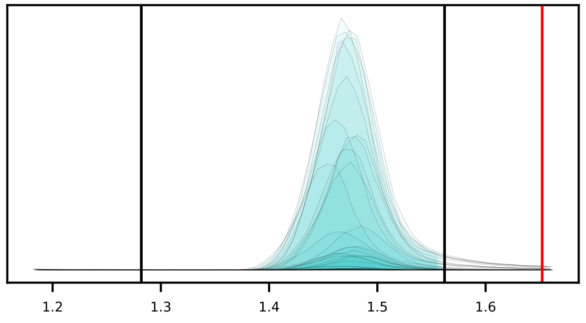


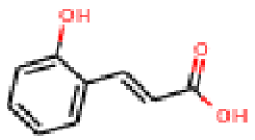


20180712_KBL_DCD-DC_Sorghum_DvsW_RzSoilRoot_QE-139_HILICZ-VF1_USHXG01063_NEG_MSMS_Rhizo-24hr-D_Z-P10_1

0001_2-hydroxycinnamic_acid_negative_163p0400_1p65

Measured M/Z = 163.0403, 1.7133 ppm difference

Expected Elution of 1.65 minutes, 1.46 min actual

MSMS Scan at 1.433 minutes

Matching M/Zs above 1E-3*max: 93.035, 118.924, 119.050, 163.040

All Matching M/Zs: 93.035, 118.924, 119.050, 163.040

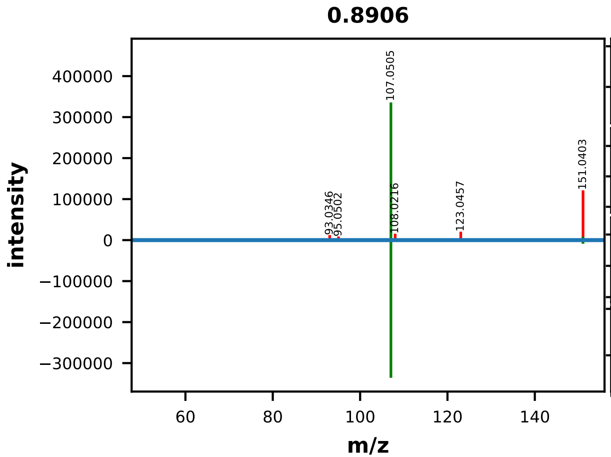


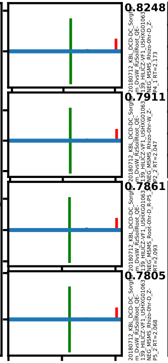


| | EMA Compound Info |
|---|---|
| Name: | (R)-(-)-Mandelic acid |
| Label: | |
| Formula: | C8H8O3 |
| Polarity: | negative |
| Monoisotopic Mass: | 152.047344116 |
| Theoretical M/Z: | 151.0400441 |
| Adduct: | |

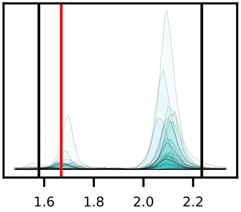


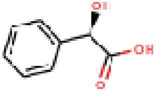


20180712_KBL_DCD-DC_Sorghum_DvsW_RzSoilRoot_QE-139_HILICZ-VF1_USHXG01063_NEG_MSMS_Rhizo-24hr-D_Z-P10_1

0002_2-hydroxyphenylacetic_acid_negative_151p0400_1p67
Measured M/Z = 151.0402, 1.1666 ppm difference
Expected Elution of 1.67 minutes, 2.10 min actual

MSMS Scan at 2.051 minutes

Matching M/Zs above 1E-3*max: 107.050, 151.028

All Matching M/Zs: 107.050, 151.028

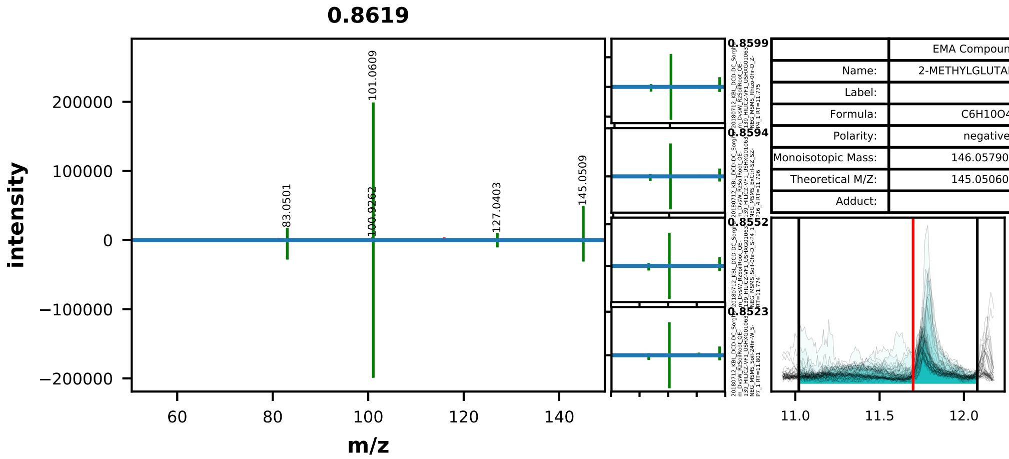


| | EMA Compound Info |
|---|---|
| Name: | 2-METHYLGLUTARIC ACID |
| Label: | |
| Formula: | C6H10O4 |
| Polarity: | negative |
| Monoisotopic Mass: | 146.0579088 |
| Theoretical M/Z: | 145.0506088 |
| Adduct: | |

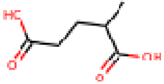

20180712_KBL_DCD-DC_Sorghum_DvsW_RzSoilRoot_QE-139_HILICZ-VF1_USHXG01063_NEG_MSMS_ExCtrl-SZ_SZ-P13_1

0003_2-methylglutaric_acid_negative_145p0506_11p70
Measured M/Z = 145.0506, 0.0711 ppm difference
Expected Elution of 11.70 minutes, 11.80 min actual

MSMS Scan at 11.790 minutes

Matching M/Zs above 1E-3*max: 73.052, 83.050, 101.061, 127.040, 145.051

All Matching M/Zs: 73.052, 83.050, 101.061, 127.040, 145.051

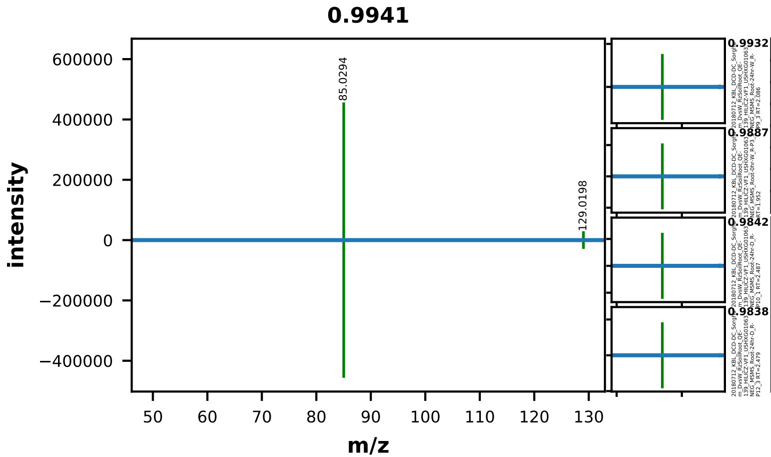


| | EMA Compound Info |
|---|---|
| Name: | Citraconic acid |
| Label: | |
| Formula: | C5H6O4 |
| Polarity: | negative |
| Monoisotopic Mass: | 130.026608672 |
| Theoretical M/Z: | 129.0193087 |
| Adduct: | |

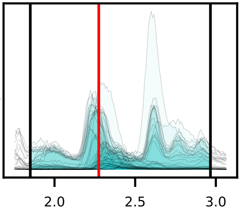


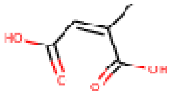


20180712_KBL_DCD-DC_Sorghum_DvsW_RzSoilRoot_QE-139_HILICZ-VF1_USHXG01063_NEG_MSMS_Root-0hr-D_R-P6_3

0004_2-methylmaleic_acid_negative_129p0193_2p27
Measured M/Z = 129.0195, 1.7635 ppm difference
Expected Elution of 2.27 minutes, 2.23 min actual

MSMS Scan at 1.997 minutes

Matching M/Zs above 1E-3*max: 85.029, 129.020

All Matching M/Zs: 85.029, 129.020

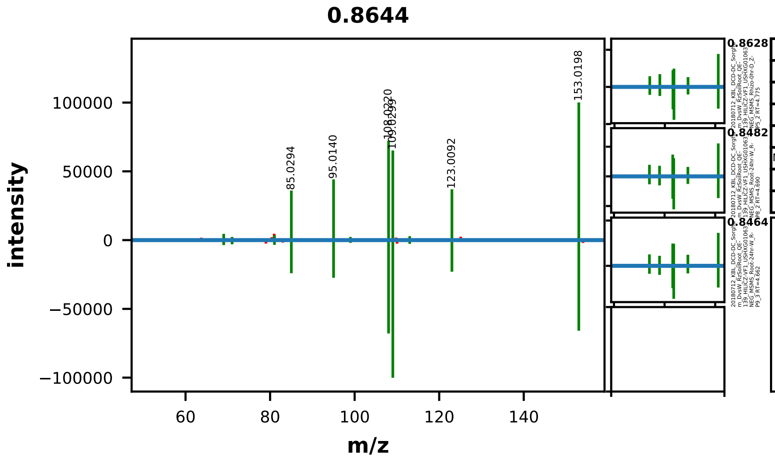


| | EMA Compound Info |
|---|---|
| Name: | 2,5-DIHYDROXYBENZOIC ACID |
| Label: | |
| Formula: | C7H6O4 |
| Polarity: | negative |
| Monoisotopic Mass: | 154.026608672 |
| Theoretical M/Z: | 153.0193087 |
| Adduct: | |

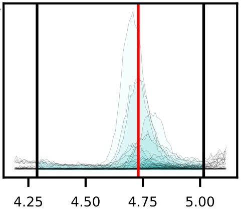


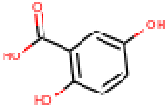


20180712_KBL_DCD-DC_Sorghum_DvsW_RzSoilRoot_QE-139_HILICZ-VF1_USHXG01063_NEG_MSMS_Root-24hr-W_R-P7_1

0007_2_5-dihydroxybenzoic_acid_negative_153p0193_4p73
Measured M/Z = 153.0195, 1.3493 ppm difference
Expected Elution of  4.73 minutes,  4.73 min actual

MSMS Scan at 4.704 minutes

Matching M/Zs above 1E-3*max: 69.034, 71.014, 81.034, 85.029, 95.014, 99.009, 108.022, 109.030, 113.025, 123.009, 153.020



All Matching M/Zs: 69.034, 71.014, 81.034, 85.029, 95.014, 99.009, 108.022, 109.030, 113.025, 123.009, 153.020

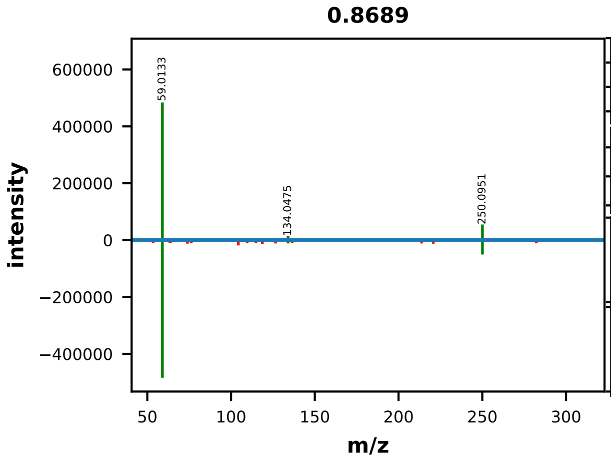


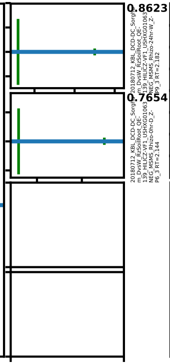


| | EMA Compound Info |
|---|---|
| Name: | 2'-deoxyadenosine |
| Label: | |
| Formula: | C10H13N5O3 |
| Polarity: | negative |
| Monoisotopic Mass: | 251.101839276 |
| Theoretical M/Z: | 310.1157393 |
| Adduct: | |

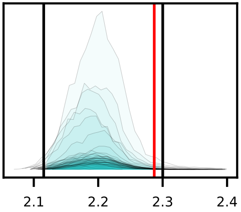


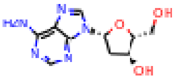


20180712_KBL_DCD-DC_Sorghum_DvsW_RzSoilRoot_QE-139_HILICZ-VF1_USHXG01063_NEG_MSMS_Rhizo-24hr-D_Z-P12_3

0008_2deoxyadenosine_negative_310p1157_2p29
Measured M/Z = 310.1163, 1.7306 ppm difference
Expected Elution of 2.29 minutes, 2.18 min actual

MSMS Scan at 2.151 minutes

Matching M/Zs above 1E-3*max: 59.013, 134.048, 250.095

All Matching M/Zs: 59.013, 134.048, 250.095

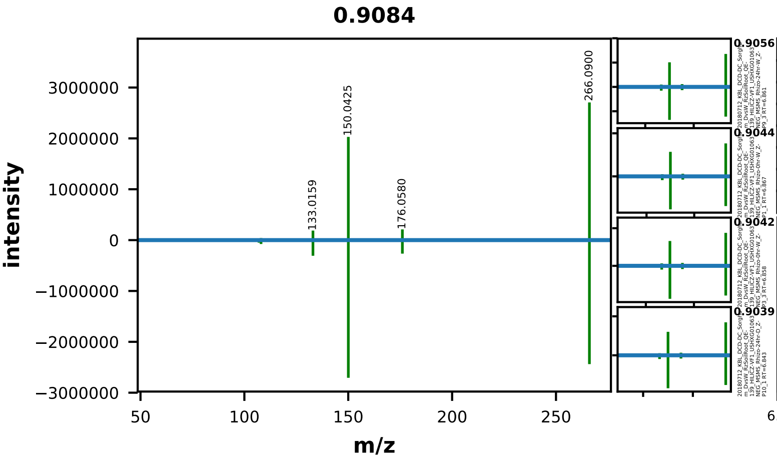


| | EMA Compound Info |
|---|---|
| Name: | 2'-deoxyguanosine |
| Label: | |
| Formula: | C10H13N5O4 |
| Polarity: | negative |
| Monoisotopic Mass: | 267.096753896 |
| Theoretical M/Z: | 266.0894539 |
| Adduct: | |

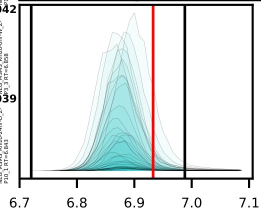


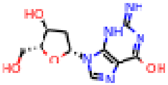


20180712_KBL_DCD-DC_Sorghum_DvsW_RzSoilRoot_QE-139_HILICZ-VF1_USHXG01063_NEG_MSMS_Rhizo-0hr-D_Z-P6_3

0009_2deoxyguanosine_negative_266p0895_6p93
Measured M/Z = 266.0900, 2.2189 ppm difference
Expected Elution of 6.93 minutes, 6.90 min actual

MSMS Scan at 6.865 minutes

Matching M/Zs above 1E-3*max: 107.037, 108.021, 133.016, 150.042, 176.058, 266.090

All Matching M/Zs: 107.037, 108.021, 133.016, 150.042, 176.058, 266.090

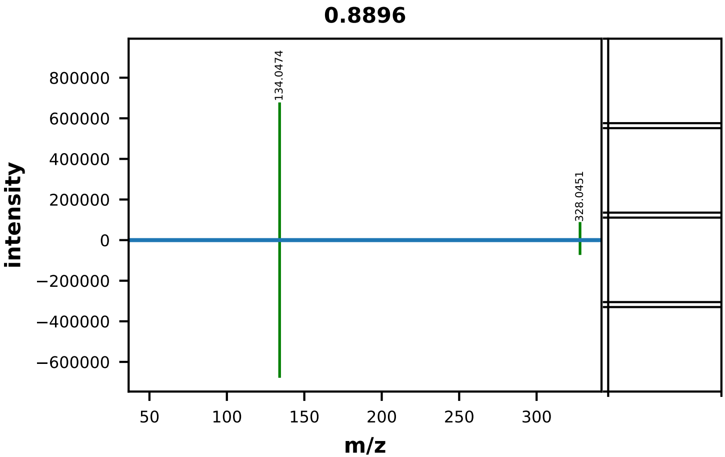


| | EMA Compound Info |
|---|---|
| Name: | 2',3'-Cyclic AMP |
| Label: | |
| Formula: | C10H12N5O6P |
| Polarity: | negative |
| Monoisotopic Mass: | 329.052519734 |
| Theoretical M/Z: | 328.0452197 |
| Adduct: | |

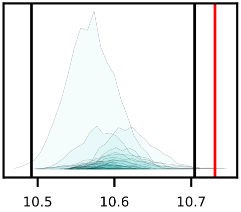


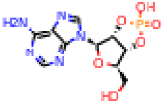


20180712_KBL_DCD-DC_Sorghum_DvsW_RzSoilRoot_QE-139_HILICZ-VF1_USHXG01063_NEG_MSMS_Soil-24hr-D_S-P10_1

0010_2_3cyclic_AMP_negative_328p0452_10p73
Measured M/Z = 328.0453, 0.2699 ppm difference
Expected Elution of 10.73 minutes, 10.57 min actual

MSMS Scan at 10.554 minutes

Matching M/Zs above 1E-3*max: 107.036, 134.047, 328.045

All Matching M/Zs: 107.036, 134.047, 328.045

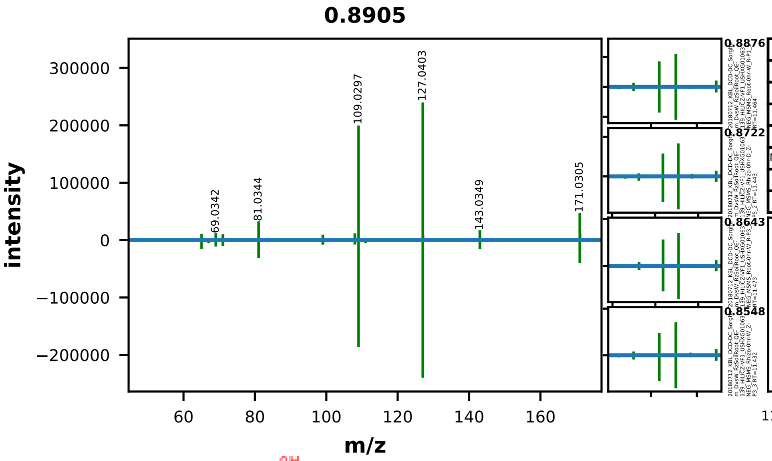


| | EMA Compound Info |
|---|---|
| Name: | 3-dehydroshikimate |
| Label: | |
| Formula: | C7H8O5 |
| Polarity: | negative |
| Monoisotopic Mass: | 172.037173356 |
| Theoretical M/Z: | 171.0298734 |
| Adduct: | |

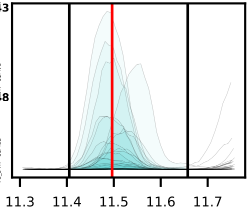


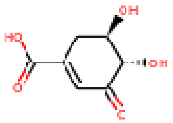


20180712_KBL_DCD-DC_Sorghum_DvsW_RzSoilRoot_QE-139_HILICZ-VF1_USHXG01063_NEG_MSMS_Root-24hr-W_R-P7_1

0012_3-dehydroshikimic_acid_negative_171p0299_11p50
Measured M/Z = 171.0300, 0.8608 ppm difference
Expected Elution of 11.50 minutes, 11.53 min actual

MSMS Scan at 11.497 minutes

Matching M/Zs above 1E-3*max: 65.039, 67.018, 69.034, 71.013, 81.034, 85.030, 99.045, 108.022, 109.030, 111.009, 127.040, 143.035, 171.030

All Matching M/Zs: 65.039, 67.018, 69.034, 71.013, 81.034, 85.030, 99.045, 108.022, 109.030, 111.009, 127.040, 143.035, 171.030

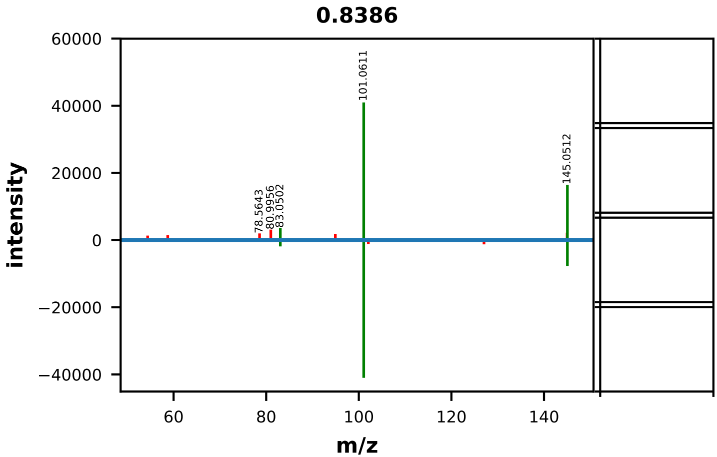


| | EMA Compound Info |
|---|---|
| Name: | 3-METHYLGLUTARIC ACID |
| Label: | |
| Formula: | C6H10O4 |
| Polarity: | negative |
| Monoisotopic Mass: | 146.0579088 |
| Theoretical M/Z: | 145.0506088 |
| Adduct: | |

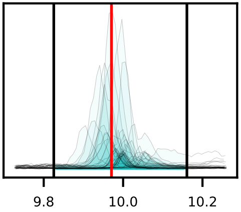


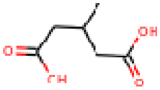


20180712_KBL_DCD-DC_Sorghum_DvsW_RzSoilRoot_QE-139_HILICZ-VF1_USHXG01063_NEG_MSMS_ExCtrl-SZ_SZ-P15_3

0015_3-methyglutaric_acid_negative_145p0506_9p97
Measured M/Z = 145.0508, 1.4875 ppm difference
Expected Elution of 9.97 minutes, 9.98 min actual

MSMS Scan at 9.998 minutes

Matching M/Zs above 1E-3*max: 83.050, 101.061, 145.051

All Matching M/Zs: 83.050, 101.061, 145.051

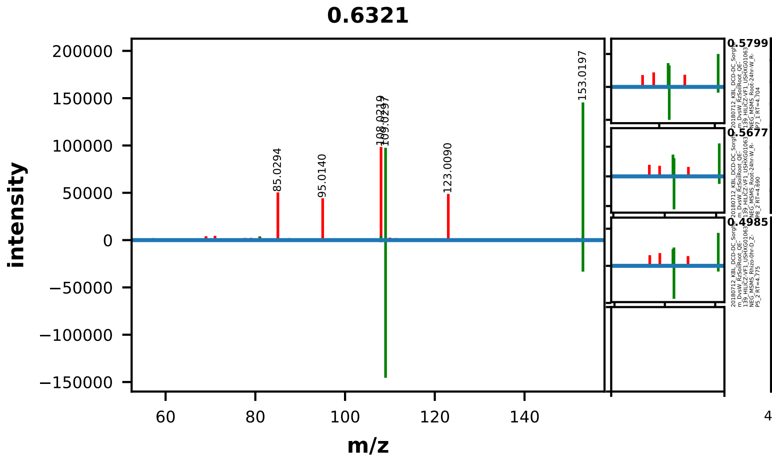


| | EMA Compound Info |
|---|---|
| Name: | 3,4-DIHYDROXYBENZOIC ACID |
| Label: | |
| Formula: | C7H6O4 |
| Polarity: | negative |
| Monoisotopic Mass: | 154.026608672 |
| Theoretical M/Z: | 153.0193087 |
| Adduct: | |

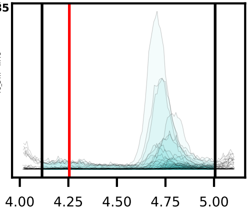

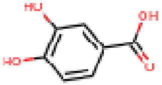

20180712_KBL_DCD-DC_Sorghum_DvsW_RzSoilRoot_QE-139_HILICZ-VF1_USHXG01063_NEG_MSMS_Root-24hr-W_R-P9_3

0017_3_4-dihydroxybenzoic_acid_negative_153p0193_4p26
Measured M/Z = 153.0195, 0.9816 ppm difference
Expected Elution of 4.26 minutes, 4.71 min actual

MSMS Scan at 4.662 minutes

Matching M/Zs above 1E-3*max: 81.034, 108.015, 109.030, 153.020

All Matching M/Zs: 81.034, 108.015, 109.030, 153.020

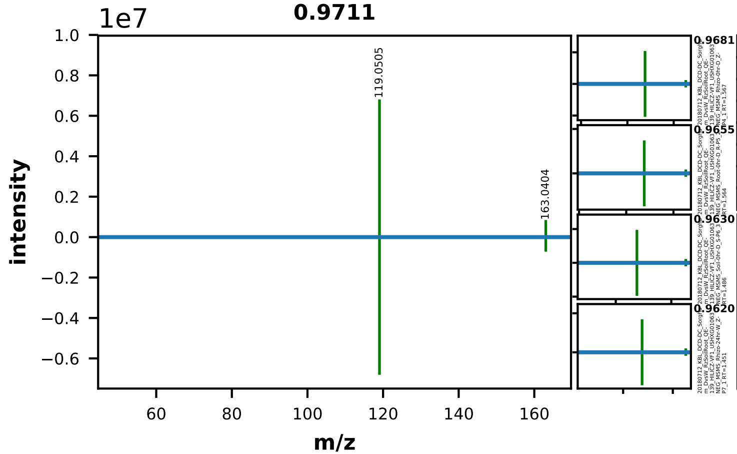


| | EMA Compound Info |
|---|---|
| Name: | p-coumaric acid |
| Label: | |
| Formula: | C9H8O3 |
| Polarity: | negative |
| Monoisotopic Mass: | 164.047344116 |
| Theoretical M/Z: | 163.0400441 |
| Adduct: | |

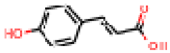

20180712_KBL_DCD-DC_Sorghum_DvsW_RzSoilRoot_QE-139_HILICZ-VF1_USHXG01063_NEG_MSMS_Root-24hr-W_R-P7_1

0020_4-coumaric_acid_negative_163p0400_1p58
Measured M/Z = 163.0403, 1.4890 ppm difference
Expected Elution of 1.58 minutes, 1.47 min actual

MSMS Scan at 1.416 minutes

Matching M/Zs above 1E-3*max: 93.035, 119.050, 163.040

All Matching M/Zs: 93.035, 119.050, 163.040

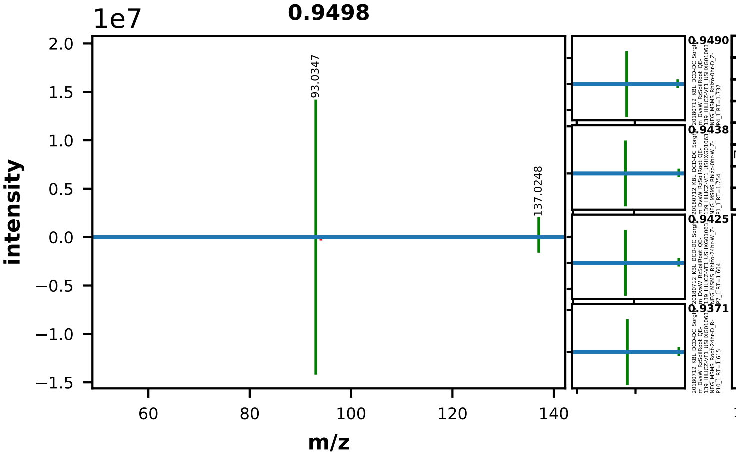


| | EMA Compound Info |
|---|---|
| Name: | 4-HYDROXYBENZOIC ACID |
| Label: | |
| Formula: | C7H6O3 |
| Polarity: | negative |
| Monoisotopic Mass: | 138.031694052 |
| Theoretical M/Z: | 137.0243941 |
| Adduct: | |

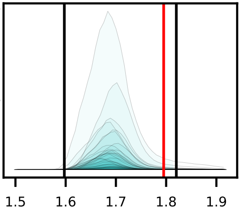


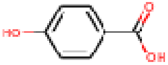


20180712_KBL_DCD-DC_Sorghum_DvsW_RzSoilRoot_QE-139_HILICZ-VF1_USHXG01063_NEG_MSMS_Rhizo-0hr-D_Z-P5_2

0023_4-hydroxybenzoic_acid_negative_137p0244_1p80
Measured M/Z = 137.0248, 2.6529 ppm difference
Expected Elution of 1.80 minutes, 1.70 min actual

MSMS Scan at 1.753 minutes

Matching M/Zs above 1E-3*max: 65.039, 93.035, 137.025

All Matching M/Zs: 65.039, 93.035, 137.025

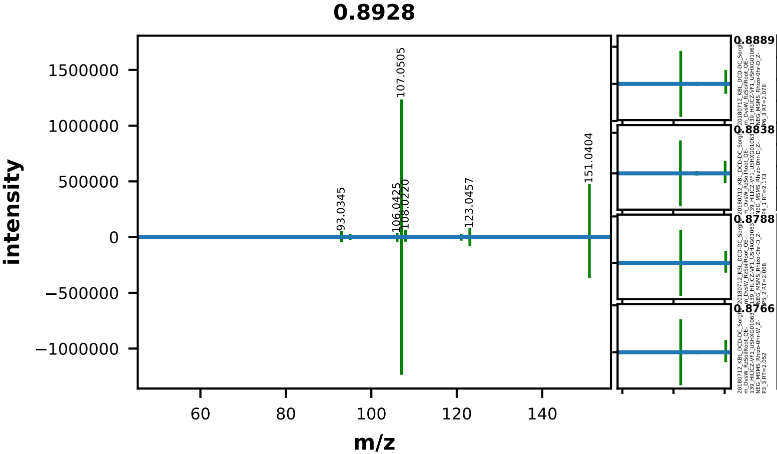


| | EMA Compound Info |
|---|---|
| Name: | 4-hydroxyphenylacetic acid |
| Label: | |
| Formula: | C8H8O3 |
| Polarity: | negative |
| Monoisotopic Mass: | 152.047344116 |
| Theoretical M/Z: | 151.0400441 |
| Adduct: | |

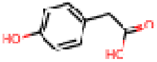

20180712_KBL_DCD-DC_Sorghum_DvsW_RzSoilRoot_QE-139_HILICZ-VF1_USHXG01063_NEG_MSMS_Root-24hr-D_R-P10_1

0024_4-hydroxyphenylacetic_acid_negative_151p0400_2p22
Measured M/Z = 151.0403, 1.7370 ppm difference
Expected Elution of 2.22 minutes, 2.08 min actual

MSMS Scan at 2.052 minutes

Matching M/Zs above 1E-3*max: 79.055, 81.034, 93.035, 95.014, 95.050, 106.043, 107.051, 108.022, 121.029, 123.046, 138.032, 151.040

All Matching M/Zs: 79.055, 81.034, 93.035, 95.014, 95.050, 106.043, 107.051, 108.022, 121.029, 123.046, 138.032, 151.040

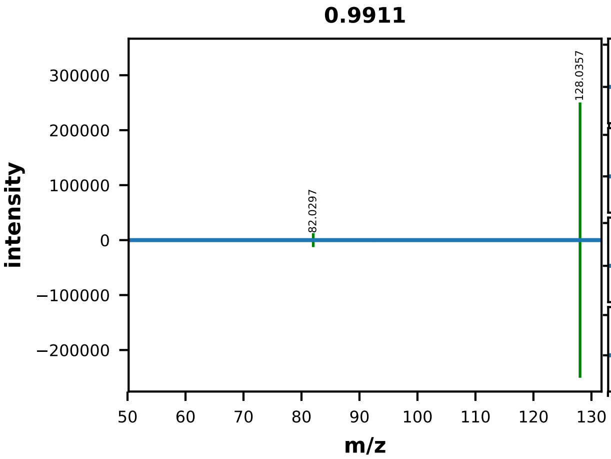


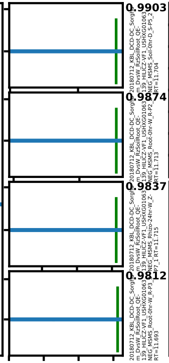


| | EMA Compound Info |
|---|---|
| Name: | L-Pyroglutamic acid |
| Label: | |
| Formula: | C5H7NO3 |
| Polarity: | negative |
| Monoisotopic Mass: | 129.042593084 |
| Theoretical M/Z: | 128.0352931 |
| Adduct: | |

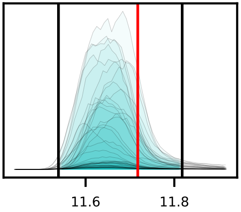


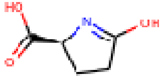


20180712_KBL_DCD-DC_Sorghum_DvsW_RzSoilRoot_QE-139_HILICZ-VF1_USHXG01063_NEG_MSMS_Rhizo-0hr-W_Z-P2_2

0027_5-oxo-proline_negative_128p0353_11p72
Measured M/Z = 128.0356, 2.3040 ppm difference
Expected Elution of 11.72 minutes, 11.64 min actual

MSMS Scan at 11.577 minutes

Matching M/Zs above 1E-3*max: 82.030, 128.036

All Matching M/Zs: 82.030, 128.036

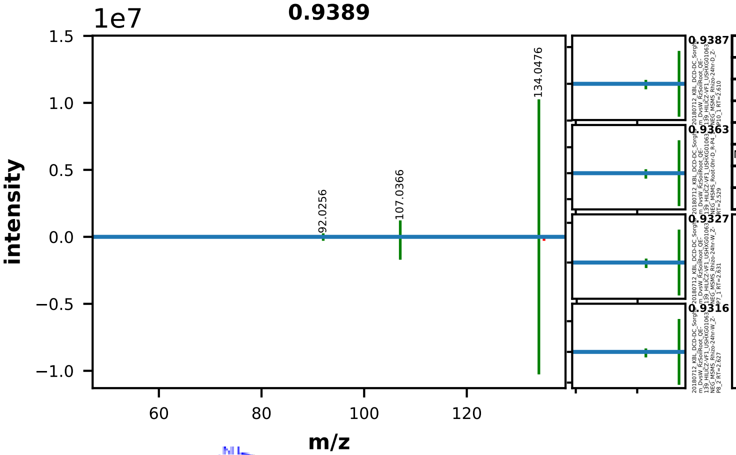


| | EMA Compound Info |
|---|---|
| Name: | adenine |
| Label: | |
| Formula: | C5H5N5 |
| Polarity: | negative |
| Monoisotopic Mass: | 135.05449516 |
| Theoretical M/Z: | 134.0471952 |
| Adduct: | |

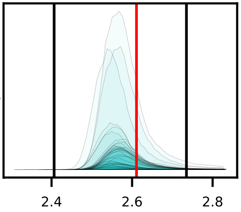

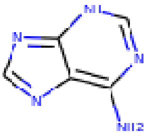

20180712_KBL_DCD-DC_Sorghum_DvsW_RzSoilRoot_QE-139_HILICZ-VF1_USHXG01063_NEG_MSMS_Rhizo-24hr-D_Z-P12_3

0031_adenine_negative_134p0472_2p61
Measured M/Z = 134.0476, 2.8154 ppm difference
Expected Elution of 2.61 minutes, 2.57 min actual

MSMS Scan at 2.627 minutes

Matching M/Zs above 1E-3*max: 92.026, 107.037, 134.048

All Matching M/Zs: 92.026, 107.037, 134.048

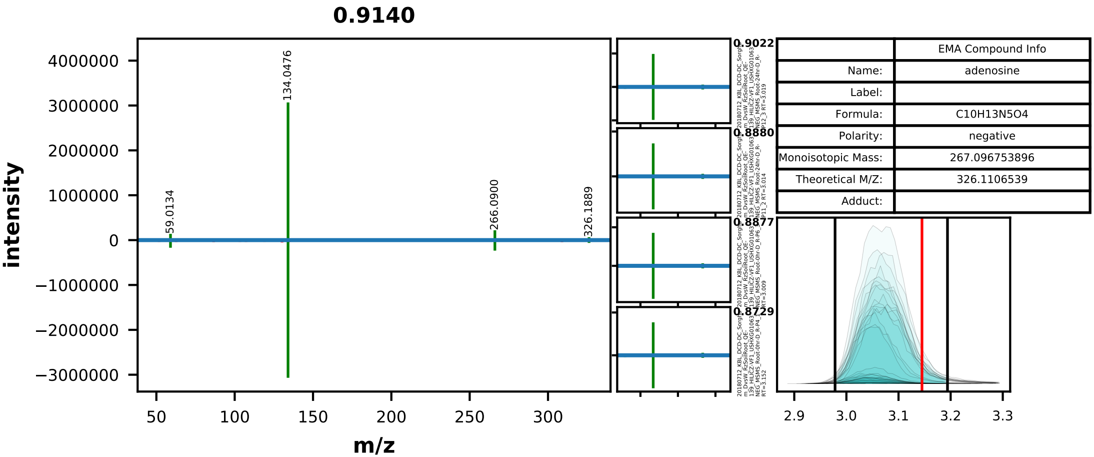


| | EMA Compound Info |
|---|---|
| Name: | adenosine |
| Label: | |
| Formula: | C10H13N5O4 |
| Polarity: | negative |
| Monoisotopic Mass: | 267.096753896 |
| Theoretical M/Z: | 326.1106539 |
| Adduct: | |

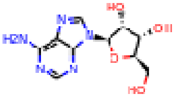


20180712_KBL_DCD-DC_Sorghum_DvsW_RzSoilRoot_QE-139_HILICZ-VF1_USHXG01063_NEG_MSMS_Root-24hr-D_R-P10_1

0032_adenosine_negative_326p1107_3p15
Measured M/Z = 326.1111, 1.4628 ppm difference
Expected Elution of 3.15 minutes, 3.05 min actual

MSMS Scan at 3.006 minutes

Matching M/Zs above 1E-3*max: 59.013, 134.048, 266.090, 326.189

All Matching M/Zs: 59.013, 134.048, 266.090, 326.189

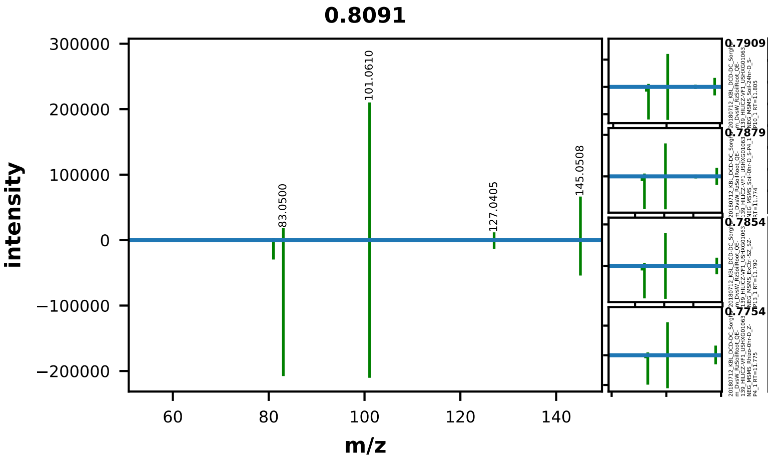


| | EMA Compound Info |
|---|---|
| Name: | adipic acid |
| Label: | |
| Formula: | C6H10O4 |
| Polarity: | negative |
| Monoisotopic Mass: | 146.0579088 |
| Theoretical M/Z: | 145.0506088 |
| Adduct: | |

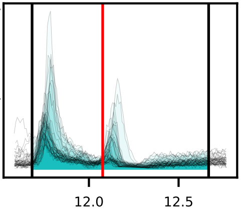


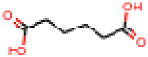


20180712_KBL_DCD-DC_Sorghum_DvsW_RzSoilRoot_QE-139_HILICZ-VF1_USHXG01063_NEG_MSMS_Soil-0hr-W_S-P1_1

0033_adipic_acid_negative_145p0506_12p08
Measured M/Z = 145.0506, 0.1253 ppm difference
Expected Elution of 12.08 minutes, 11.77 min actual

MSMS Scan at 11.778 minutes

Matching M/Zs above 1E-3*max: 80.996, 83.050, 101.061, 127.040, 145.040, 145.051

All Matching M/Zs: 80.996, 83.050, 101.061, 127.040, 145.040, 145.051

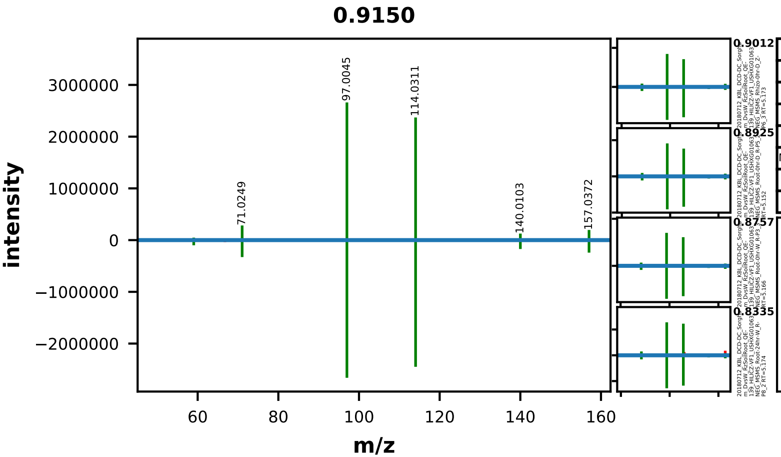


| | EMA Compound Info |
|---|---|
| Name: | allantoin |
| Label: | |
| Formula: | C4H6N4O3 |
| Polarity: | negative |
| Monoisotopic Mass: | 158.043990052 |
| Theoretical M/Z: | 157.0366901 |
| Adduct: | |

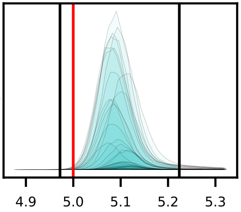


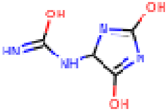


20180712_KBL_DCD-DC_Sorghum_DvsW_RzSoilRoot_QE-139_HILICZ-VF1_USHXG01063_NEG_MSMS_Rhizo-0hr-D_Z-P5_2

0034_allantoin_negative_157p0367_5p00
Measured M/Z = 157.0369, 1.4713 ppm difference
Expected Elution of 5.00 minutes, 5.10 min actual

MSMS Scan at 5.166 minutes

Matching M/Zs above 1E-3*max: 59.025, 71.025, 72.009, 97.004, 114.031, 140.010, 157.037

All Matching M/Zs: 59.025, 71.025, 72.009, 97.004, 114.031, 140.010, 157.037

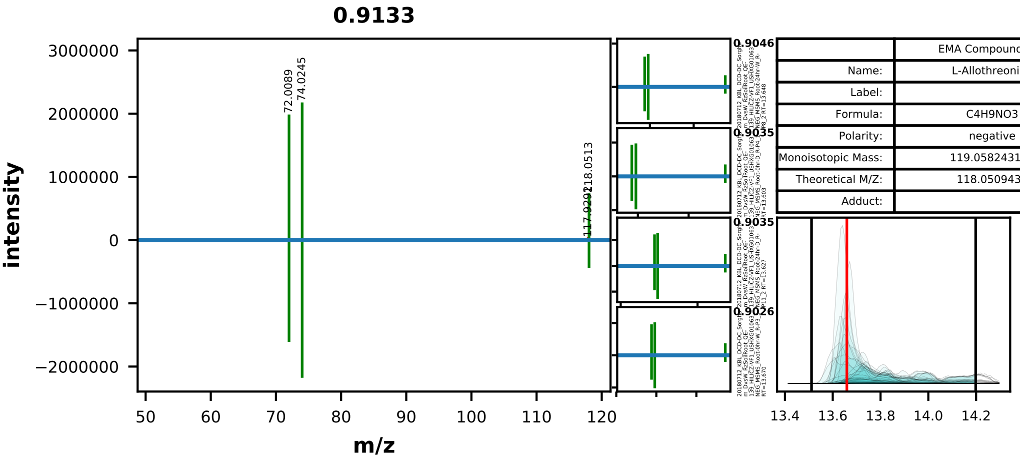


| | EMA Compound Info |
|---|---|
| Name: | L-Allothreonine |
| Label: | |
| Formula: | C4H9NO3 |
| Polarity: | negative |
| Monoisotopic Mass: | 119.058243148 |
| Theoretical M/Z: | 118.0509431 |
| Adduct: | |

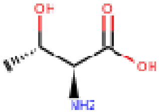


20180712_KBL_DCD-DC_Sorghum_DvsW_RzSoilRoot_QE-139_HILICZ-VF1_USHXG01063_NEG_MSMS_Rhizo-0hr-D_Z-P6_3

0035_allothreonine_negative_118p0509_13p66
Measured M/Z = 118.0512, 2.0683 ppm difference
Expected Elution of 13.66 minutes, 13.61 min actual

MSMS Scan at 13.606 minutes

Matching M/Zs above 1E-3*max: 72.009, 74.025, 118.051

All Matching M/Zs: 72.009, 74.025, 118.051

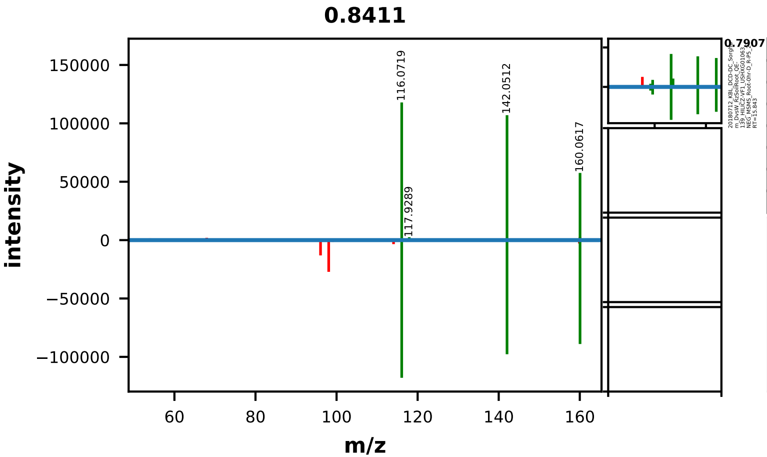


| | EMA Compound Info |
|---|---|
| Name: | L-2-Aminoadipic acid |
| Label: | |
| Formula: | C6H11NO4 |
| Polarity: | negative |
| Monoisotopic Mass: | 161.068807832 |
| Theoretical M/Z: | 160.0615078 |
| Adduct: | |

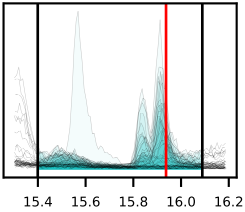


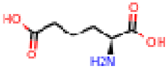


20180712_KBL_DCD-DC_Sorghum_DvsW_RzSoilRoot_QE-139_HILICZ-VF1_USHXG01063_NEG_MSMS_Rhizo-24hr-W_Z-P7_1

0036_alpha-aminoadipic_acid_negative_160p0615_15p94
Measured M/Z = 160.0616, 0.5473 ppm difference
Expected Elution of 15.94 minutes, 15.57 min actual

MSMS Scan at 15.575 minutes

Matching M/Zs above 1E-3*max: 116.072, 117.929, 142.051, 160.062

All Matching M/Zs: 116.072, 117.929, 142.051, 160.062

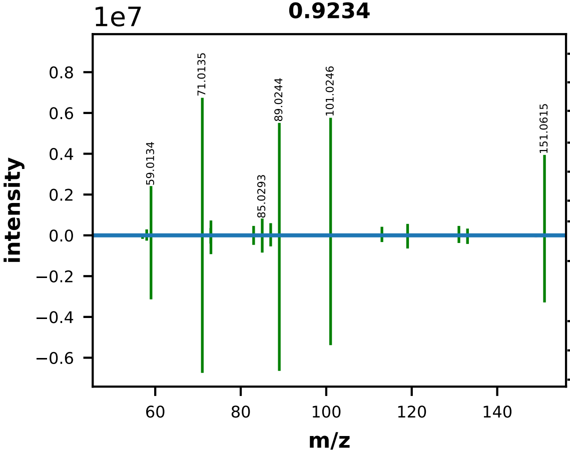


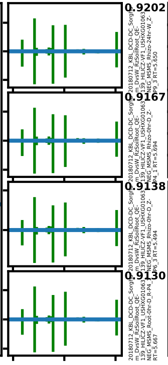


| | EMA Compound Info |
|---|---|
| Name: | L-arabitol |
| Label: | |
| Formula: | C5H12O5 |
| Polarity: | negative |
| Monoisotopic Mass: | 152.068473484 |
| Theoretical M/Z: | 151.0611735 |
| Adduct: | |

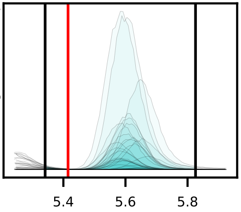


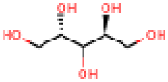


20180712_KBL_DCD-DC_Sorghum_DvsW_RzSoilRoot_QE-139_HILICZ-VF1_USHXG01063_NEG_MSMS_Rhizo-0hr-D_Z-P5_2

0038_arabitol_negative_151p0612_5p42
Measured M/Z = 151.0615, 2.2671 ppm difference
Expected Elution of 5.42 minutes, 5.58 min actual

MSMS Scan at 5.613 minutes

Matching M/Zs above 1E-3*max: 55.018, 57.034, 58.005, 59.013, 71.014, 72.993, 73.029, 75.009, 83.014, 85.029, 87.009, 89.024, 101.025, 103.040, 113.025, 119.035, 131.036, 133.051, 151.061

All Matching M/Zs: 55.018, 57.034, 58.005, 59.013, 71.014, 72.993, 73.029, 75.009, 83.014, 85.029, 87.009, 89.024, 101.025, 103.040, 113.025, 119.035, 131.036, 133.051, 151.061

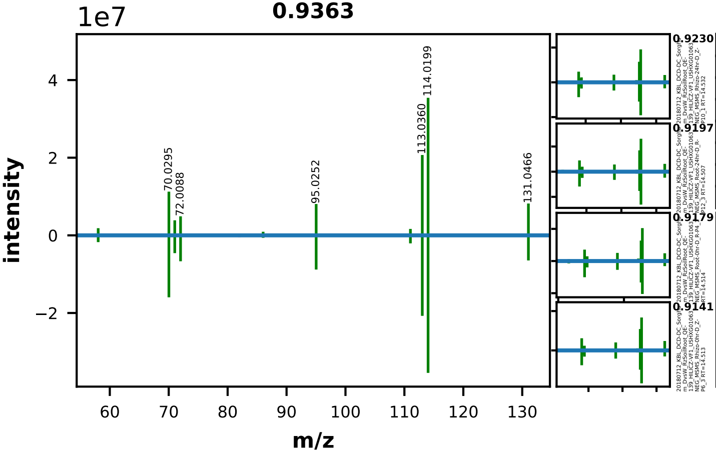


| | EMA Compound Info |
|---|---|
| Name: | L-asparagine |
| Label: | |
| Formula: | C4H8N2O3 |
| Polarity: | negative |
| Monoisotopic Mass: | 132.053492116 |
| Theoretical M/Z: | 131.0461921 |
| Adduct: | |

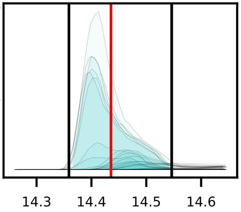

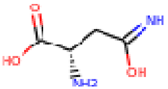


20180712_KBL_DCD-DC_Sorghum_DvsW_RzSoilRoot_QE-139_HILICZ-VF1_USHXG01063_NEG_MSMS_Root-0hr-D_R-P6_3

0040_asparagine_negative_131p0462_14p44
Measured M/Z = 131.0465, 1.9995 ppm difference
Expected Elution of 14.44 minutes, 14.40 min actual

MSMS Scan at 14.510 minutes

Matching M/Zs above 1E-3*max: 58.029, 70.030, 71.014, 71.025, 72.009, 86.025, 95.025, 111.020, 113.036, 114.020, 131.047

All Matching M/Zs: 58.029, 70.030, 71.014, 71.025, 72.009, 86.025, 95.025, 111.020, 113.036, 114.020, 131.047

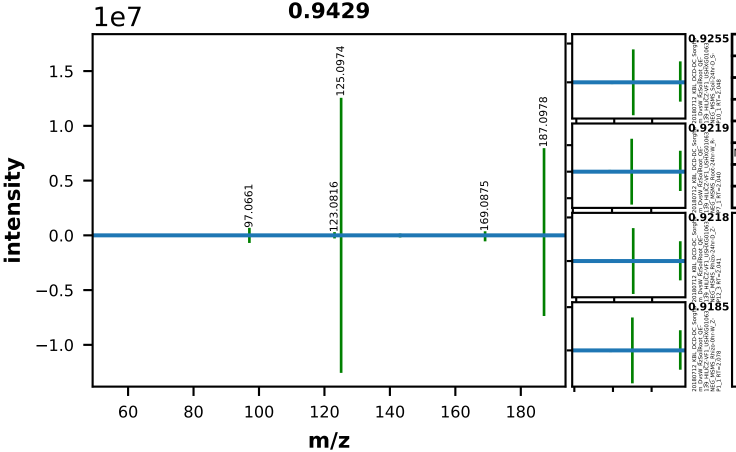


| | EMA Compound Info |
|---|---|
| Name: | azelaic acid |
| Label: | |
| Formula: | C9H16O4 |
| Polarity: | negative |
| Monoisotopic Mass: | 188.104858992 |
| Theoretical M/Z: | 187.097559 |
| Adduct: | |

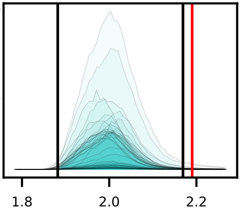

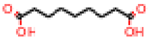

20180712_KBL_DCD-DC_Sorghum_DvsW_RzSoilRoot_QE-139_HILICZ-VF1_USHXG01063_NEG_MSMS_Rhizo-24hr-D_Z-P10_1

0041_azelaic_acid_negative_187p0976_2p19
Measured M/Z = 187.0977, 0.8684 ppm difference
Expected Elution of 2.19 minutes, 2.00 min actual

MSMS Scan at 2.033 minutes

Matching M/Zs above 1E-3*max: 57.034, 97.066, 123.082, 125.097, 143.108, 169.087, 187.098

All Matching M/Zs: 57.034, 97.066, 123.082, 125.097, 143.108, 169.087, 187.098

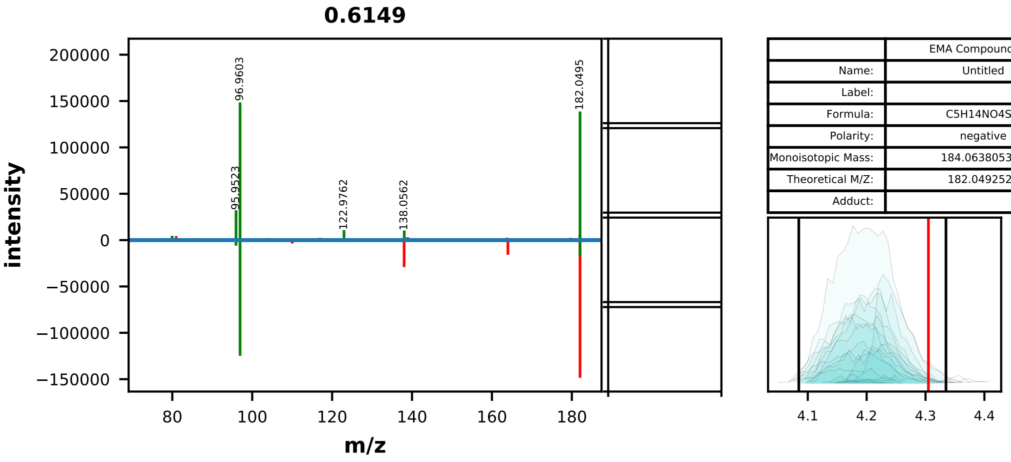


| EMA Compound Info | |
|---|---|
| Name: | Untitled |
| Label: | |
| Formula: | C5H14NO4S+ |
| Polarity: | negative |
| Monoisotopic Mass: | 184.063805348 |
| Theoretical M/Z: | 182.0492524 |
| Adduct: | |

20180712_KBL_DCD-DC_Sorghum_DvsW_RzSoilRoot_QE-139_HILICZ-VF1_USHXG01063_NEG_MSMS_Rhizo-0hr-W_Z-P3_3

0044_choline_o-sulfuric_acid_negative_182p0493_4p30
Measured M/Z = 182.0494, 0.6351 ppm difference
Expected Elution of 4.30 minutes, 4.18 min actual

MSMS Scan at 4.224 minutes

Matching M/Zs above 1E-3*max: 79.957, 95.952, 96.960, 122.976, 138.056, 182.050

All Matching M/Zs: 79.957, 95.952, 96.960, 122.976, 138.056, 182.050

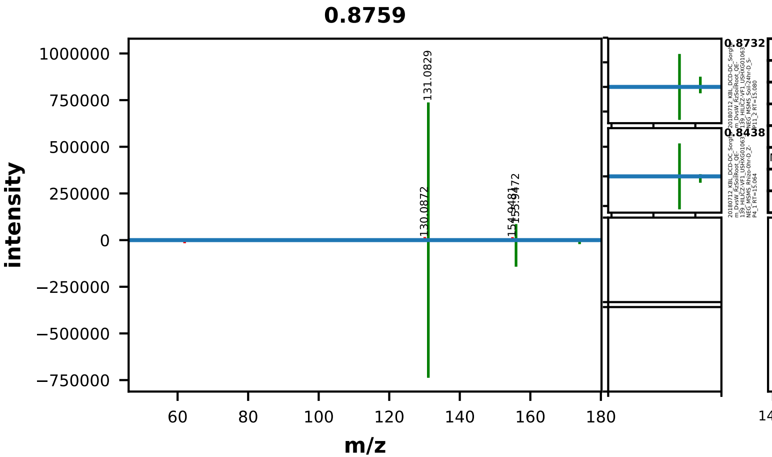


| EMA Compound Info | |
|---|---|
| Name: | L-citrulline |
| Label: | |
| Formula: | C6H13N3O3 |
| Polarity: | negative |
| Monoisotopic Mass: | 175.095691276 |
| Theoretical M/Z: | 174.0883913 |
| Adduct: | |

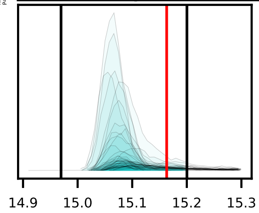

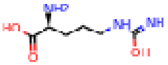

20180712_KBL_DCD-DC_Sorghum_DvsW_RzSoilRoot_QE-139_HILICZ-VF1_USHXG01063_NEG_MSMS_Rhizo-24hr-D_Z-P10_1

0046_citrulline_negative_174p0884_15p16
Measured M/Z = 174.0884, 0.1508 ppm difference
Expected Elution of 15.16 minutes, 15.06 min actual

MSMS Scan at 15.061 minutes

Matching M/Zs above 1E-3*max: 131.083, 155.947, 173.958, 174.089

All Matching M/Zs: 131.083, 155.947, 173.958, 174.089

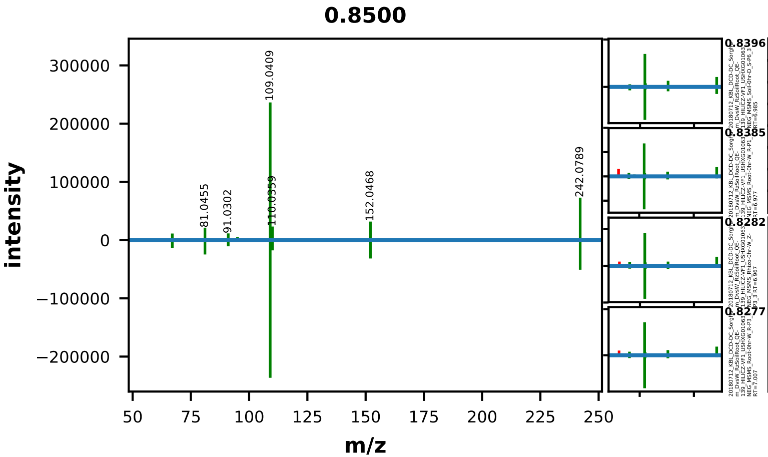


| EMA Compound Info | |
|---|---|
| Name: | cytidine |
| Label: | |
| Formula: | C9H13N3O5 |
| Polarity: | negative |
| Monoisotopic Mass: | 243.085520516 |
| Theoretical M/Z: | 242.0782205 |
| Adduct: | |

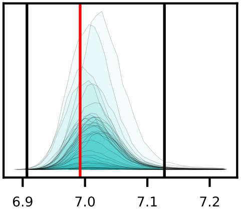

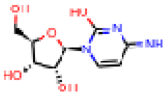

20180712_KBL_DCD-DC_Sorghum_DvsW_RzSoilRoot_QE-139_HILICZ-VF1_USHXG01063_NEG_MSMS_Soil-0hr-D_S-P5_2

0049_cytidine_negative_242p0782_6p99
Measured M/Z = 242.0787, 2.0459 ppm difference
Expected Elution of 6.99 minutes, 6.99 min actual

MSMS Scan at 6.982 minutes

Matching M/Zs above 1E-3*max: 67.030, 81.046, 91.030, 95.025, 109.041, 110.036, 152.047, 242.079

All Matching M/Zs: 67.030, 81.046, 91.030, 95.025, 109.041, 110.036, 152.047, 242.079

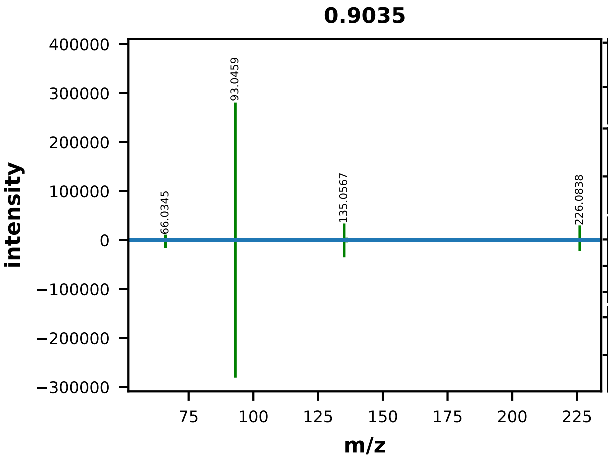


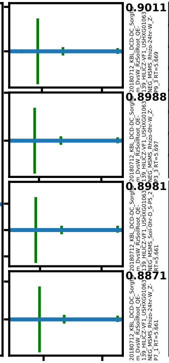


| | EMA Compound Info |
|---|---|
| Name: | 2'-deoxycytidine |
| Label: | |
| Formula: | C9H13N3O4 |
| Polarity: | negative |
| Monoisotopic Mass: | 227.090605896 |
| Theoretical M/Z: | 226.0833059 |
| Adduct: | |

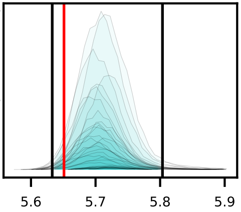


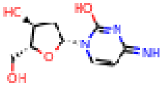


20180712_KBL_DCD-DC_Sorghum_DvsW_RzSoilRoot_QE-139_HILICZ-VF1_USHXG01063_NEG_MSMS_Rhizo-0hr-W_Z-P1_1

0051_deoxycytidine_negative_226p0833_5p65
Measured M/Z = 226.0838, 2.0272 ppm difference
Expected Elution of 5.65 minutes, 5.70 min actual

MSMS Scan at 5.676 minutes

Matching M/Zs above 1E-3*max: 66.035, 93.046, 135.057, 136.052, 183.078, 226.084

All Matching M/Zs: 66.035, 93.046, 135.057, 136.052, 183.078, 226.084

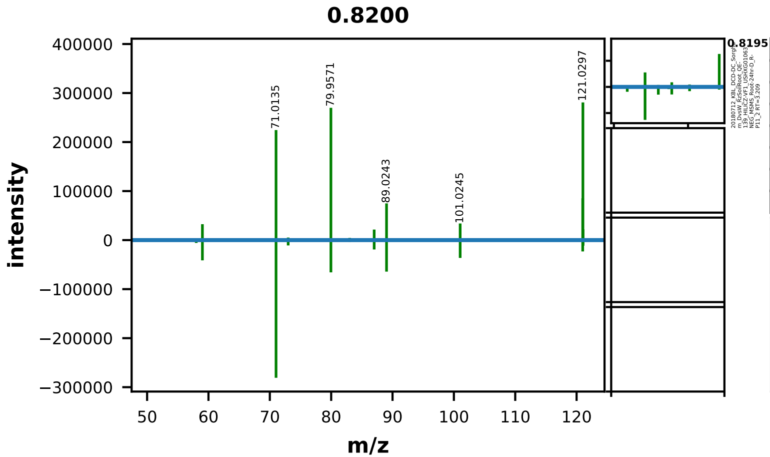


| | EMA Compound Info |
|---|---|
| Name: | meso-Erythritol |
| Label: | |
| Formula: | C4H10O4 |
| Polarity: | negative |
| Monoisotopic Mass: | 122.0579088 |
| Theoretical M/Z: | 121.0506088 |
| Adduct: | |

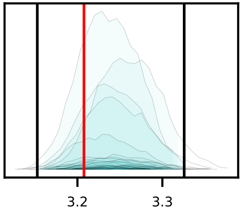


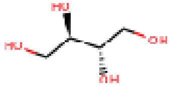


20180712_KBL_DCD-DC_Sorghum_DvsW_RzSoilRoot_QE-139_HILICZ-VF1_USHXG01063_NEG_MSMS_Rhizo-0hr-D_Z-P4_1

0054_erythritol_negative_121p0506_3p21
Measured M/Z = 121.0507, 0.8766 ppm difference
Expected Elution of 3.21 minutes, 3.25 min actual

MSMS Scan at 3.248 minutes

Matching M/Zs above 1E-3*max: 55.018, 58.006, 59.013, 71.014, 72.993, 79.957, 83.014, 87.009, 89.024, 101.025, 120.997, 121.030, 121.050

All Matching M/Zs: 55.018, 58.006, 59.013, 71.014, 72.993, 79.957, 83.014, 87.009, 89.024, 101.025, 120.997, 121.030, 121.050

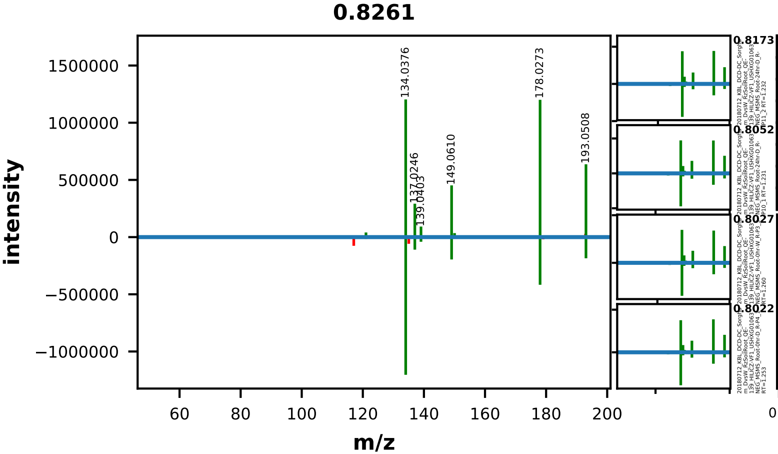


| | EMA Compound Info |
|---|---|
| Name: | ferulic acid |
| Label: | |
| Formula: | C10H10O4 |
| Polarity: | negative |
| Monoisotopic Mass: | 194.0579088 |
| Theoretical M/Z: | 193.0506088 |
| Adduct: | |

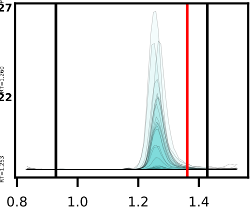


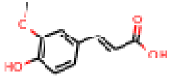


20180712_KBL_DCD-DC_Sorghum_DvsW_RzSoilRoot_QE-139_HILICZ-VF1_USHXG01063_NEG_MSMS_Root-0hr-D_R-P6_3

0055_ferulic_acid_negative_193p0506_1p36
Measured M/Z = 193.0508, 0.8495 ppm difference
Expected Elution of 1.36 minutes, 1.27 min actual

MSMS Scan at 1.263 minutes

Matching M/Zs above 1E-3*max: 121.030, 134.038, 137.025, 139.040, 149.061, 150.032, 178.027, 193.051

All Matching M/Zs: 121.030, 134.038, 137.025, 139.040, 149.061, 150.032, 178.027, 193.051

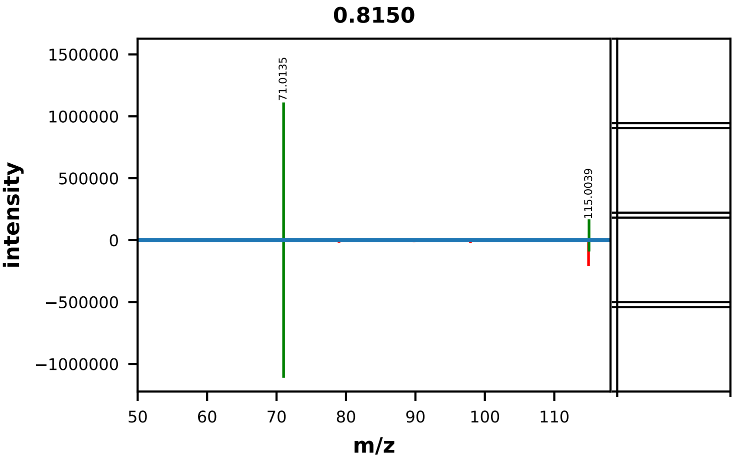


| | EMA Compound Info |
|---|---|
| Name: | fumaric acid |
| Label: | |
| Formula: | C4H4O4 |
| Polarity: | negative |
| Monoisotopic Mass: | 116.010958608 |
| Theoretical M/Z: | 115.0036586 |
| Adduct: | |

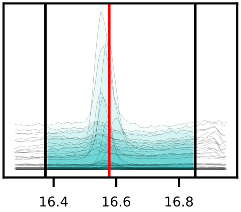


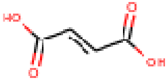


20180712_KBL_DCD-DC_Sorghum_DvsW_RzSoilRoot_QE-139_HILICZ-VF1_USHXG01063_NEG_MSMS_Rhizo-24hr-D_Z-P10_1

0057_fumaric_acid_negative_115p0037_16p58
Measured M/Z = 115.0038, 1.6327 ppm difference
Expected Elution of 16.58 minutes, 16.58 min actual

MSMS Scan at 16.573 minutes

Matching M/Zs above 1E-3*max: 71.014, 115.004

All Matching M/Zs: 71.014, 115.004

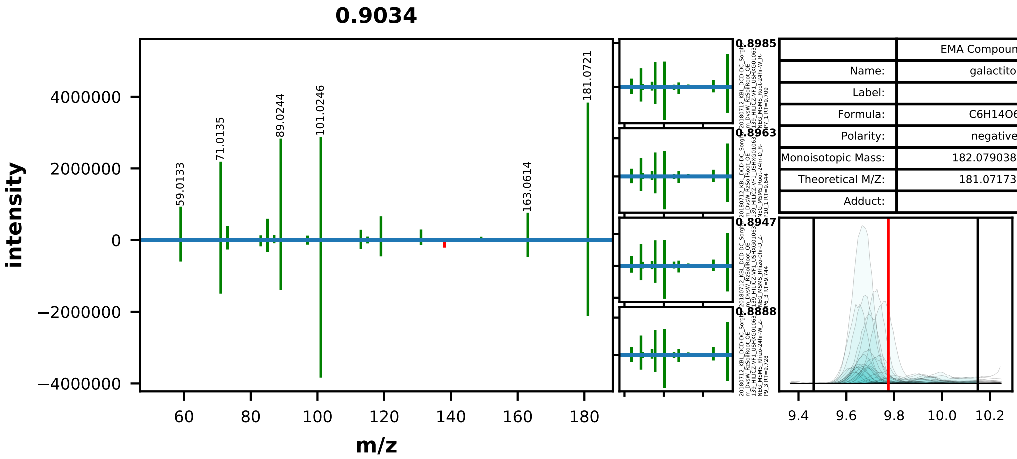


| | EMA Compound Info |
|---|---|
| Name: | galactitol |
| Label: | |
| Formula: | C6H14O6 |
| Polarity: | negative |
| Monoisotopic Mass: | 182.079038168 |
| Theoretical M/Z: | 181.0717382 |
| Adduct: | |

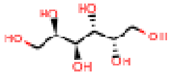

20180712_KBL_DCD-DC_Sorghum_DvsW_RzSoilRoot_QE-139_HILICZ-VF1_USHXG01063_NEG_MSMS_Root-24hr-D_R-P11_2

0058_galactitol_negative_181p0717_9p78
Measured M/Z = 181.0720, 1.5688 ppm difference
Expected Elution of 9.78 minutes, 9.70 min actual

MSMS Scan at 9.653 minutes

Matching M/Zs above 1E-3*max: 55.018, 57.034, 58.005, 59.013, 71.014, 72.993, 73.029, 83.014, 85.029, 87.009, 89.024, 97.030, 99.009, 101.025, 113.025, 115.040, 119.035, 131.035, 149.046, 161.046, 163.061, 181.072

All Matching M/Zs: 55.018, 57.034, 58.005, 59.013, 71.014, 72.993, 73.029, 83.014, 85.029, 87.009, 89.024, 97.030, 99.009, 101.025, 113.025, 115.040, 119.035, 131.035, 149.046, 161.046, 163.061, 181.072

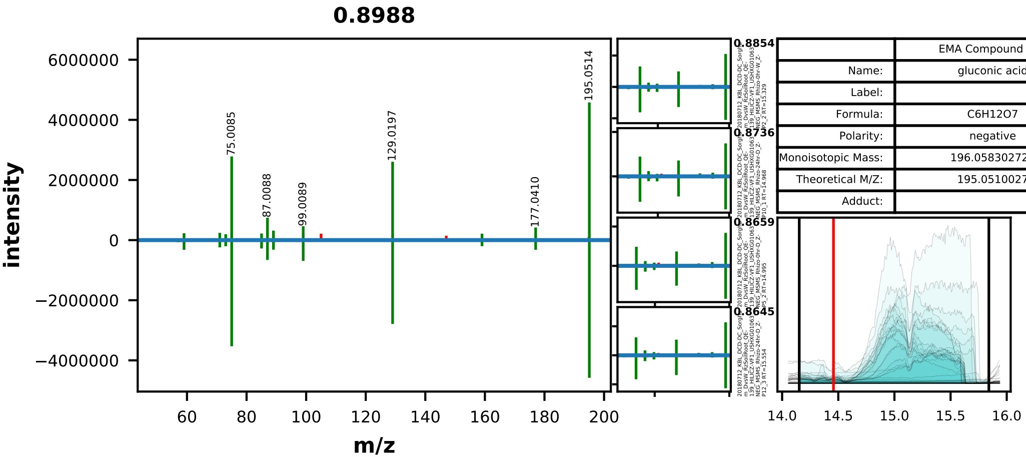


| | EMA Compound Info |
|---|---|
| Name: | gluconic acid |
| Label: | |
| Formula: | C6H12O7 |
| Polarity: | negative |
| Monoisotopic Mass: | 196.058302724 |
| Theoretical M/Z: | 195.0510027 |
| Adduct: | |

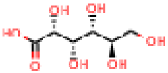

20180712_KBL_DCD-DC_Sorghum_DvsW_RzSoilRoot_QE-139_HILICZ-VF1_USHXG01063_NEG_MSMS_Rhizo-0hr-W_Z-P3_3

0060_gluconic_acid_negative_195p0510_14p46
Measured M/Z = 195.0513, 1.5201 ppm difference
Expected Elution of 14.46 minutes, 15.48 min actual

MSMS Scan at 15.618 minutes

Matching M/Zs above 1E-3*max: 57.034, 59.013, 71.014, 72.993, 75.009, 85.029, 87.009, 89.024, 99.009, 101.025, 129.020, 159.030, 177.041, 195.051



All Matching M/Zs: 57.034, 59.013, 71.014, 72.993, 75.009, 85.029, 87.009, 89.024, 99.009, 101.025, 129.020, 159.030, 177.041, 195.051

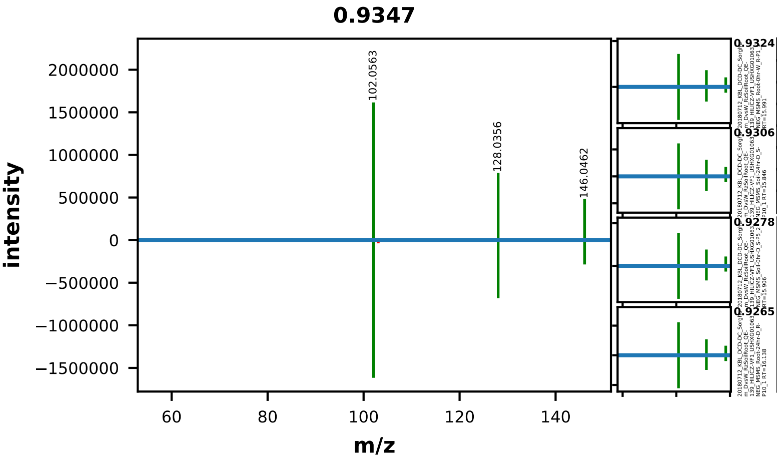


| | EMA Compound Info |
|---|---|
| Name: | L-glutamic acid |
| Label: | |
| Formula: | C5H9NO4 |
| Polarity: | negative |
| Monoisotopic Mass: | 147.053157768 |
| Theoretical M/Z: | 146.0458578 |
| Adduct: | |

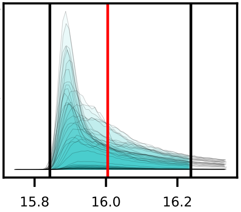


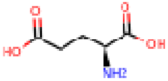


20180712_KBL_DCD-DC_Sorghum_DvsW_RzSoilRoot_QE-139_HILICZ-VF1_USHXG01063_NEG_MSMS_Rhizo-24hr-W_Z-P7_1

0064_glutamic_acid_negative_146p0459_16p00
Measured M/Z = 146.0461, 1.9811 ppm difference
Expected Elution of 16.00 minutes, 15.92 min actual

MSMS Scan at 16.137 minutes

Matching M/Zs above 1E-3*max: 82.030, 85.030, 102.056, 128.036, 146.046

All Matching M/Zs: 82.030, 85.030, 102.056, 128.036, 146.046

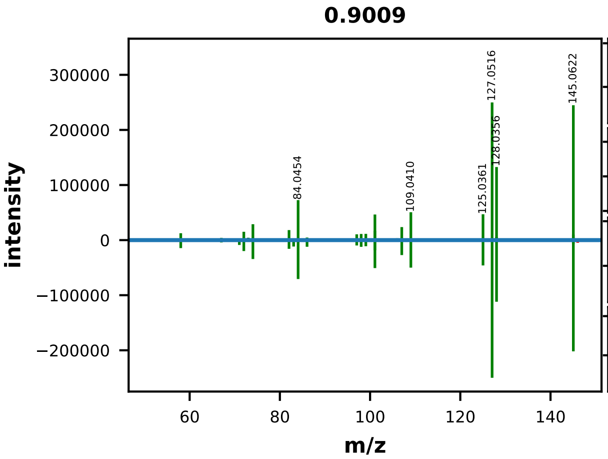


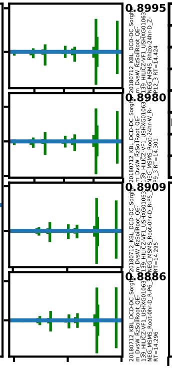


|  | EMA Compound Info |
|---|---|
| Name: | L-glutamine |
| Label: | |
| Formula: | C5H10N2O3 |
| Polarity: | negative |
| Monoisotopic Mass: | 146.06914218 |
| Theoretical M/Z: | 145.0618422 |
| Adduct: | |

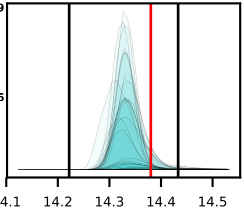


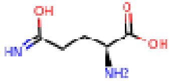


20180712_KBL_DCD-DC_Sorghum_DvsW_RzSoilRoot_QE-139_HILICZ-VF1_USHXG01063_NEG_MSMS_Soil-24hr-D_S-P11_2

0065_glutamine_negative_145p0618_14p38
Measured M/Z = 145.0622, 2.2198 ppm difference
Expected Elution of 14.38 minutes, 14.34 min actual

MSMS Scan at 14.309 minutes

Matching M/Zs above 1E-3*max: 53.921, 58.029, 67.030, 71.014, 72.009, 72.993, 74.024, 82.030, 83.050, 84.045, 86.025, 97.041, 98.025, 99.057, 101.061, 101.072, 106.017, 107.025, 109.041, 125.036, 127.052, 128.026, 128.036, 145.052, 145.062



All Matching M/Zs: 53.921, 58.029, 67.030, 71.014, 72.009, 72.993, 74.024, 82.030, 83.050, 84.045, 86.025, 97.041, 98.025, 99.057, 101.061, 101.072, 106.017, 107.025, 109.041, 125.036, 127.052, 128.026, 128.036, 145.052, 145.062

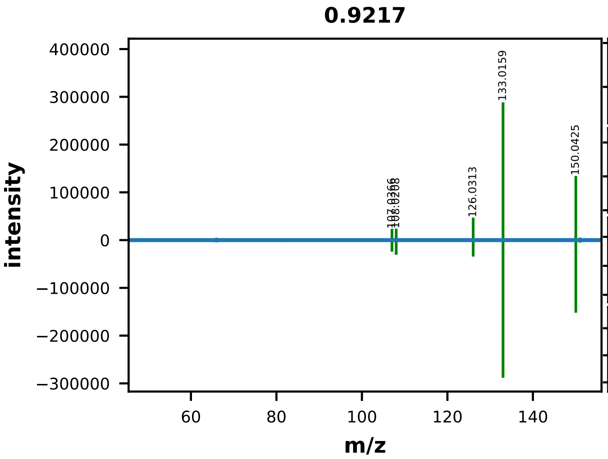


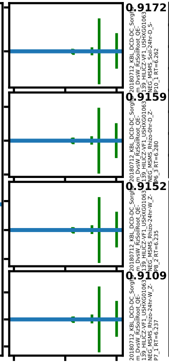


| | EMA Compound Info |
|---|---|
| Name: | guanine |
| Label: | |
| Formula: | C5H5N5O |
| Polarity: | negative |
| Monoisotopic Mass: | 151.04940978 |
| Theoretical M/Z: | 150.0421098 |
| Adduct: | |

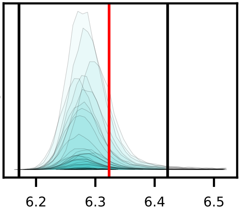


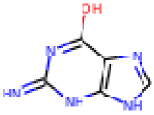


20180712_KBL_DCD-DC_Sorghum_DvsW_RzSoilRoot_QE-139_HILICZ-VF1_USHXG01063_NEG_MSMS_Rhizo-24hr-D_Z-P11_2

0066_guanine_negative_150p0421_6p32
Measured M/Z = 150.0424, 1.7817 ppm difference
Expected Elution of 6.32 minutes, 6.28 min actual

MSMS Scan at 6.266 minutes

Matching M/Zs above 1E-3*max: 66.009, 82.041, 107.037, 108.021, 126.031, 133.016, 150.042, 151.027

All Matching M/Zs: 66.009, 82.041, 107.037, 108.021, 126.031, 133.016, 150.042, 151.027

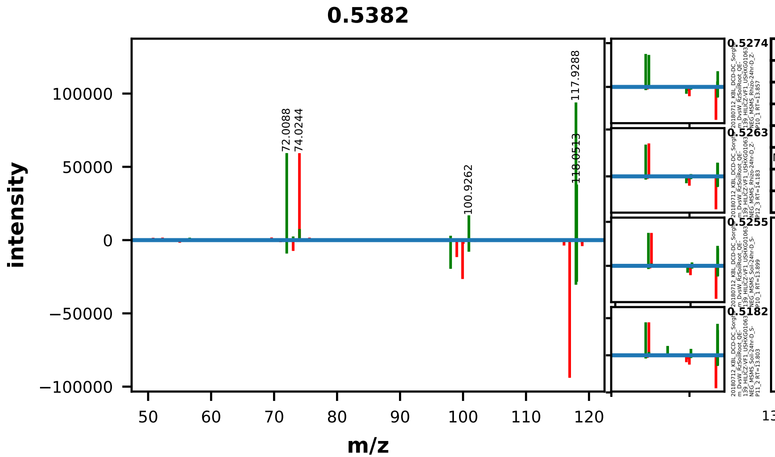


| | EMA Compound Info |
|---|---|
| Name: | L-homoserine |
| Label: | |
| Formula: | C4H9NO3 |
| Polarity: | negative |
| Monoisotopic Mass: | 119.058243148 |
| Theoretical M/Z: | 118.0509431 |
| Adduct: | |

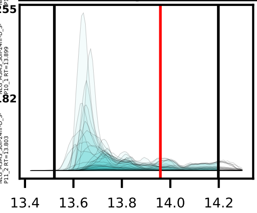


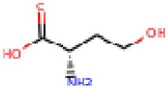


20180712_KBL_DCD-DC_Sorghum_DvsW_RzSoilRoot_QE-139_HILICZ-VF1_USHXG01063_NEG_MSMS_Rhizo-24hr-W_Z-P7_1

0068_homoserine_negative_118p0509_13p96
Measured M/Z = 118.0511, 1.2114 ppm difference
Expected Elution of 13.96 minutes, 13.70 min actual

MSMS Scan at 13.953 minutes

Matching M/Zs above 1E-3*max: 56.595, 72.009, 73.029, 74.032, 98.025, 100.926, 117.929, 118.051

All Matching M/Zs: 56.595, 72.009, 73.029, 74.032, 98.025, 100.926, 117.929, 118.051

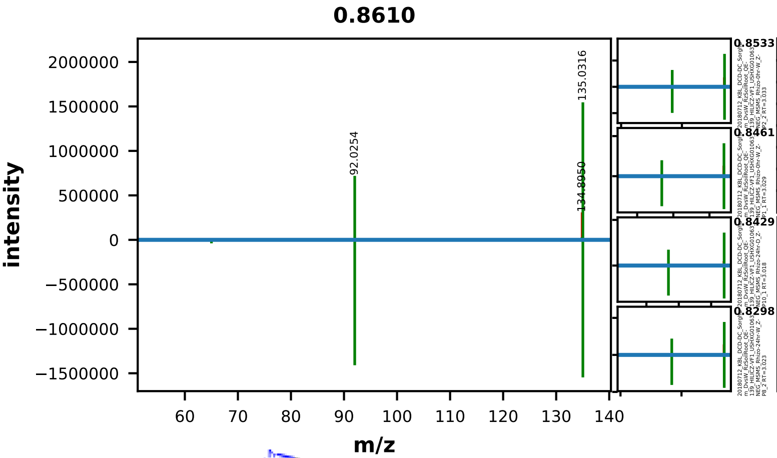


| | EMA Compound Info |
|---|---|
| Name: | hypoxanthine |
| Label: | |
| Formula: | C5H4N4O |
| Polarity: | negative |
| Monoisotopic Mass: | 136.038510748 |
| Theoretical M/Z: | 135.0312107 |
| Adduct: | |

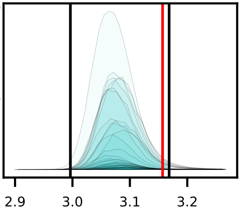


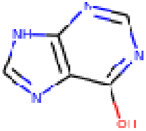


20180712_KBL_DCD-DC_Sorghum_DvsW_RzSoilRoot_QE-139_HILICZ-VF1_USHXG01063_NEG_MSMS_Rhizo-24hr-W_Z-P7_1

0069_hypoxanthine_negative_135p0312_3p16
Measured M/Z = 135.0315, 2.1051 ppm difference
Expected Elution of 3.16 minutes, 3.07 min actual

MSMS Scan at 3.034 minutes

Matching M/Zs above 1E-3*max: 65.014, 66.009, 92.025, 133.016, 134.866, 135.032

All Matching M/Zs: 65.014, 66.009, 92.025, 133.016, 134.866, 135.032

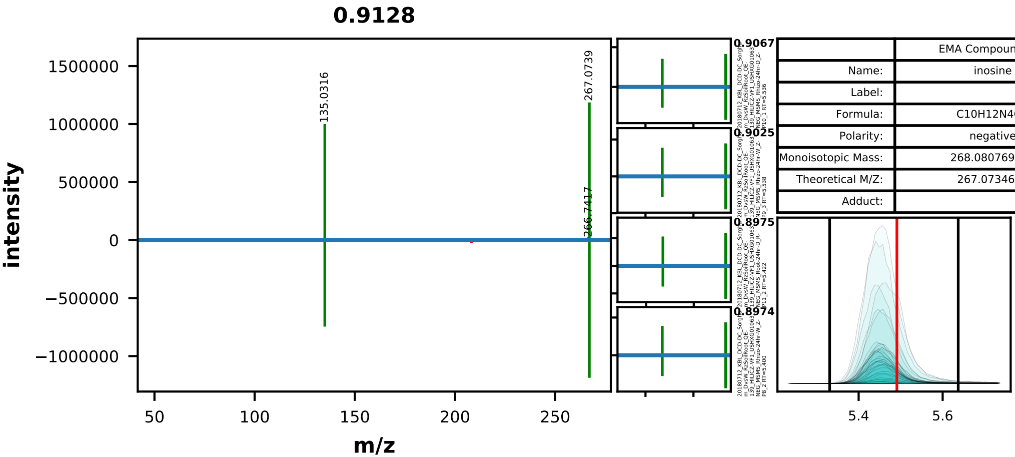


| | EMA Compound Info |
|---|---|
| Name: | inosine |
| Label: | |
| Formula: | C10H12N4O5 |
| Polarity: | negative |
| Monoisotopic Mass: | 268.080769484 |
| Theoretical M/Z: | 267.0734695 |
| Adduct: | |

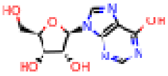


20180712_KBL_DCD-DC_Sorghum_DvsW_RzSoilRoot_QE-139_HILICZ-VF1_USHXG01063_NEG_MSMS_Rhizo-0hr-D_Z-P6_3

0071_inosine_negative_267p0735_5p49
Measured M/Z = 267.0738, 1.3230 ppm difference
Expected Elution of 5.49 minutes, 5.46 min actual

MSMS Scan at 5.552 minutes

Matching M/Zs above 1E-3*max: 135.032, 267.074

All Matching M/Zs: 135.032, 267.074

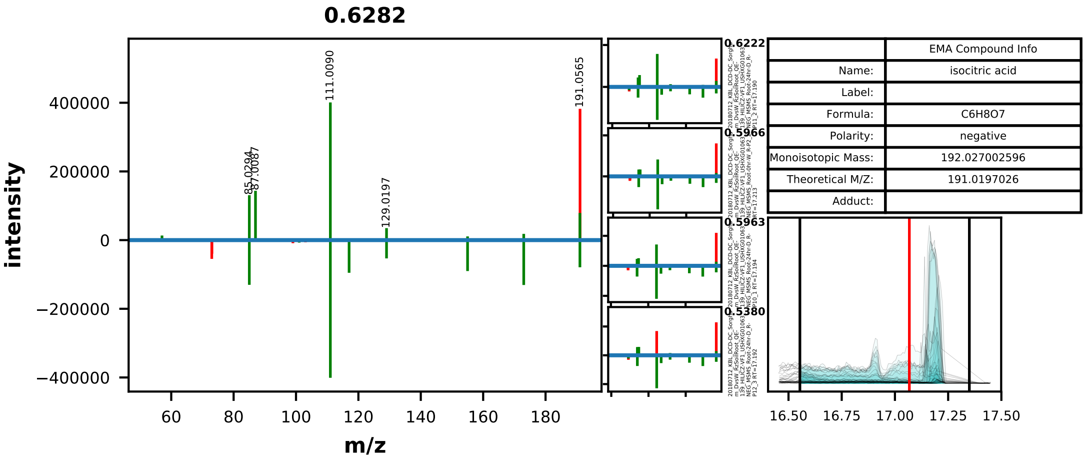


| | EMA Compound Info |
|---|---|
| Name: | isocitric acid |
| Label: | |
| Formula: | C6H8O7 |
| Polarity: | negative |
| Monoisotopic Mass: | 192.027002596 |
| Theoretical M/Z: | 191.0197026 |
| Adduct: | |

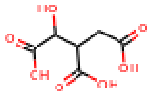

20180712_KBL_DCD-DC_Sorghum_DvsW_RzSoilRoot_QE-139_HILICZ-VF1_USHXG01063_NEG_MSMS_Root-0hr-D_R-P4_1

0072_isocitric_acid_negative_191p0197_17p07
Measured M/Z = 191.0200, 1.6366 ppm difference
Expected Elution of 17.07 minutes, 17.17 min actual

MSMS Scan at 17.188 minutes

Matching M/Zs above 1E-3*max: 57.034, 73.029, 85.029, 87.009, 101.025, 111.009, 117.019, 129.020, 154.999, 173.009, 191.020

All Matching M/Zs: 57.034, 73.029, 85.029, 87.009, 101.025, 111.009, 117.019, 129.020, 154.999, 173.009, 191.020

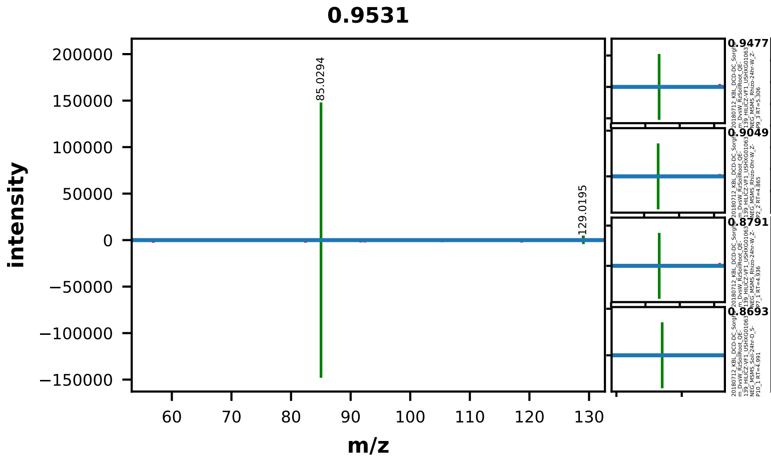


| | EMA Compound Info |
|---|---|
| Name: | Itaconic acid |
| Label: | |
| Formula: | C5H6O4 |
| Polarity: | negative |
| Monoisotopic Mass: | 130.026608672 |
| Theoretical M/Z: | 129.0193087 |
| Adduct: | |

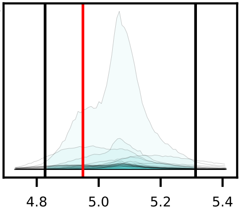

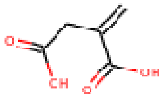

20180712_KBL_DCD-DC_Sorghum_DvsW_RzSoilRoot_QE-139_HILICZ-VF1_USHXG01063_NEG_MSMS_Rhizo-0hr-W_Z-P1_1

0073_itaconic_acid_negative_129p0193_4p95
Measured M/Z = 129.0196, 2.5341 ppm difference
Expected Elution of 4.95 minutes, 5.14 min actual

MSMS Scan at 4.980 minutes

Matching M/Zs above 1E-3*max: 85.029, 129.056

All Matching M/Zs: 85.029, 129.056

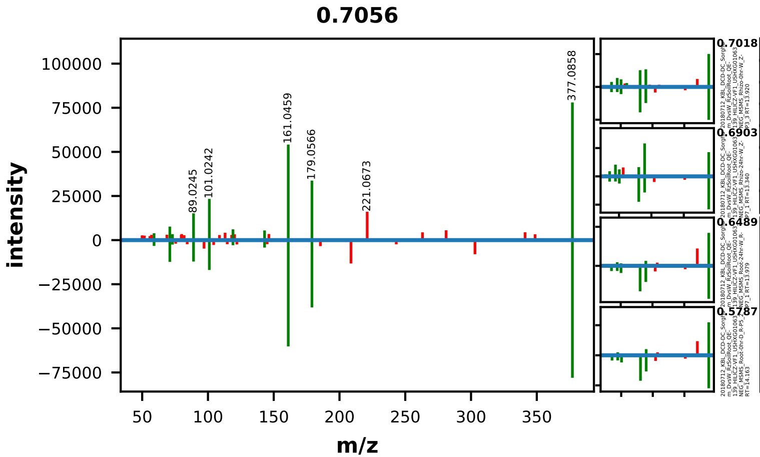


| | EMA Compound Info |
|---|---|
| Name: | lactose |
| Label: | |
| Formula: | C12H22O11 |
| Polarity: | negative |
| Monoisotopic Mass: | 342.116211524 |
| Theoretical M/Z: | 377.0856115 |
| Adduct: | |

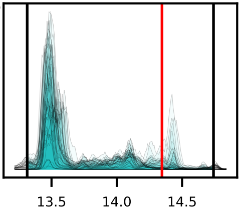

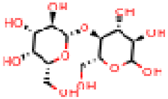

20180712_KBL_DCD-DC_Sorghum_DvsW_RzSoilRoot_QE-139_HILICZ-VF1_USHXG01063_NEG_MSMS_Soil-0hr-W_S-P3_3

0074_lactose_negative_377p0856_14p35
Measured M/Z = 377.0863, 1.8625 ppm difference
Expected Elution of 14.35 minutes, 13.47 min actual

MSMS Scan at 14.079 minutes

Matching M/Zs above 1E-3*max: 59.013, 71.014, 73.029, 89.024, 101.024, 119.035, 143.035, 161.046, 179.057, 377.086



All Matching M/Zs: 59.013, 71.014, 73.029, 89.024, 101.024, 119.035, 143.035, 161.046, 179.057, 377.086

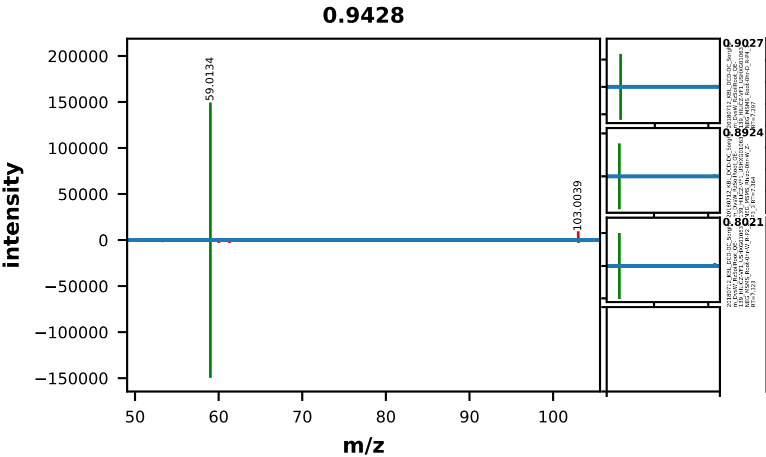


| | EMA Compound Info |
|---|---|
| Name: | malonic acid |
| Label: | |
| Formula: | C3H4O4 |
| Polarity: | negative |
| Monoisotopic Mass: | 104.010958608 |
| Theoretical M/Z: | 103.0036586 |
| Adduct: | |

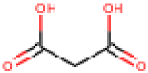


20180712_KBL_DCD-DC_Sorghum_DvsW_RzSoilRoot_QE-139_HILICZ-VF1_USHXG01063_NEG_MSMS_Root-0hr-D_R-P5_2

0076_malonic_acid_negative_103p0037_4p95
Measured M/Z = 103.0038, 1.7503 ppm difference
Expected Elution of 4.95 minutes, 6.66 min actual

MSMS Scan at 8.099 minutes

Matching M/Zs above 1E-3*max: 59.013, 103.040

All Matching M/Zs: 59.013, 103.040

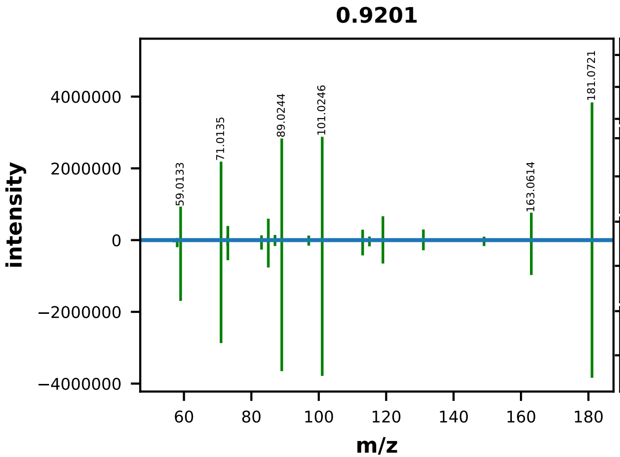


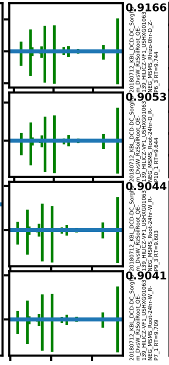


| | EMA Compound Info |
|---|---|
| Name: | D-mannitol |
| Label: | |
| Formula: | C6H14O6 |
| Polarity: | negative |
| Monoisotopic Mass: | 182.079038168 |
| Theoretical M/Z: | 181.0717382 |
| Adduct: | |

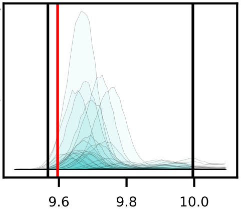


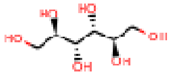


20180712_KBL_DCD-DC_Sorghum_DvsW_RzSoilRoot_QE-139_HILICZ-VF1_USHXG01063_NEG_MSMS_Root-24hr-D_R-P11_2

0077_mannitol_negative_181p0717_9p60
Measured M/Z = 181.0720, 1.5849 ppm difference
Expected Elution of 9.60 minutes, 9.70 min actual

MSMS Scan at 9.653 minutes

Matching M/Zs above 1E-3*max: 57.034, 58.005, 59.013, 71.014, 72.993, 73.029, 83.014, 85.029, 87.009, 89.024, 97.030, 99.009, 101.025, 113.025, 115.040, 119.035, 131.035, 149.046, 161.046, 163.061, 181.072



All Matching M/Zs: 57.034, 58.005, 59.013, 71.014, 72.993, 73.029, 83.014, 85.029, 87.009, 89.024, 97.030, 99.009, 101.025, 113.025, 115.040, 119.035, 131.035, 149.046, 161.046, 163.061, 181.072

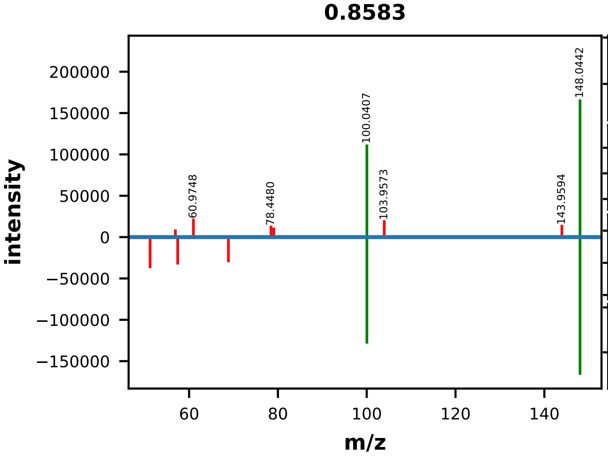


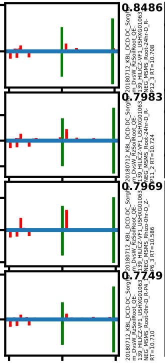


| | EMA Compound Info |
|---|---|
| Name: | L-methionine |
| Label: | |
| Formula: | C5H11NO2S |
| Polarity: | negative |
| Monoisotopic Mass: | 149.051049592 |
| Theoretical M/Z: | 148.0437496 |
| Adduct: | |

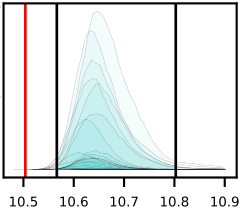


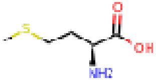


20180712_KBL_DCD-DC_Sorghum_DvsW_RzSoilRoot_QE-139_HILICZ-VF1_USHXG01063_NEG_MSMS_Root-0hr-D_R-P5_2

0079_methionine_negative_148p0437_10p50
Measured M/Z = 148.0441, 2.3198 ppm difference
Expected Elution of 10.50 minutes, 10.65 min actual

MSMS Scan at 10.719 minutes

Matching M/Zs above 1E-3*max: 100.041, 148.044

All Matching M/Zs: 100.041, 148.044

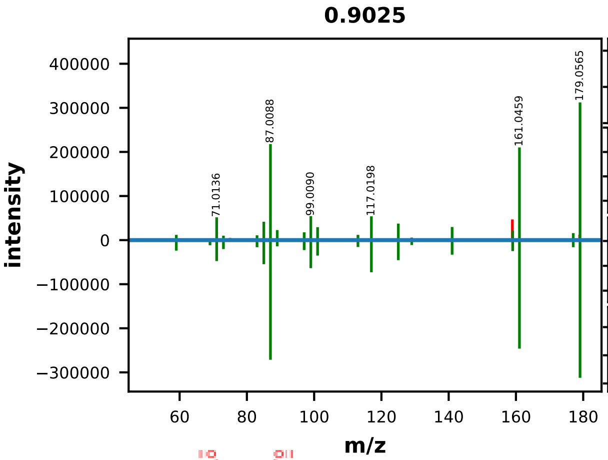


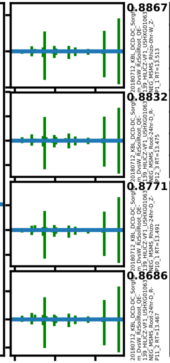


| | EMA Compound Info |
|---|---|
| Name: | myo-inositol |
| Label: | |
| Formula: | C6H12O6 |
| Polarity: | negative |
| Monoisotopic Mass: | 180.063388104 |
| Theoretical M/Z: | 179.0560881 |
| Adduct: | |

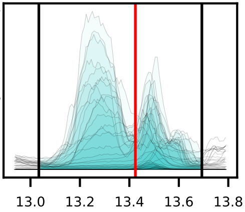


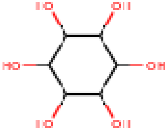


20180712_KBL_DCD-DC_Sorghum_DvsW_RzSoilRoot_QE-139_HILICZ-VF1_USHXG01063_NEG_MSMS_Soil-24hr-D_S-P12_3

0080_myo-inositol_negative_179p0561_13p42
Measured M/Z = 179.0563, 1.2468 ppm difference
Expected Elution of 13.42 minutes, 13.48 min actual

MSMS Scan at 13.466 minutes

Matching M/Zs above 1E-3*max: 57.034, 59.013, 69.034, 71.014, 73.029, 75.009, 81.034, 83.014, 85.029, 87.009, 89.025, 95.014, 97.030, 99.009, 101.025, 113.025, 117.020, 125.025, 129.020, 141.020, 159.031, 161.046, 177.041, 179.057

All Matching M/Zs: 57.034, 59.013, 69.034, 71.014, 73.029, 75.009, 81.034, 83.014, 85.029, 87.009, 89.025, 95.014, 97.030, 99.009, 101.025, 113.025, 117.020, 125.025, 129.020, 141.020, 159.031, 161.046, 177.041, 179.057

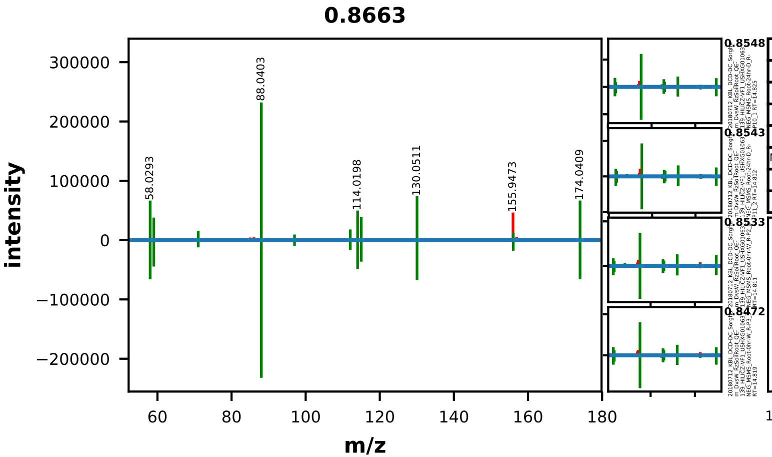


| | EMA Compound Info |
|---|---|
| Name: | 997-55-7 |
| Label: | |
| Formula: | C6H9NO5 |
| Polarity: | negative |
| Monoisotopic Mass: | 175.048072388 |
| Theoretical M/Z: | 174.0407724 |
| Adduct: | |

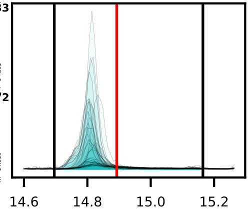

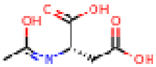

20180712_KBL_DCD-DC_Sorghum_DvsW_RzSoilRoot_QE-139_HILICZ-VF1_USHXG01063_NEG_MSMS_Rhizo-0hr-D_Z-P4_1

0081_N-acetyl-aspartic_acid_negative_174p0408_14p89
Measured M/Z = 174.0407, 0.2800 ppm difference
Expected Elution of 14.89 minutes, 14.83 min actual

MSMS Scan at 14.830 minutes

Matching M/Zs above 1E-3*max: 58.029, 59.013, 71.013, 88.040, 96.993, 112.041, 114.020, 115.004, 130.051, 132.030, 156.031, 174.041



All Matching M/Zs: 58.029, 59.013, 71.013, 88.040, 96.993, 112.041, 114.020, 115.004, 130.051, 132.030, 156.031, 174.041

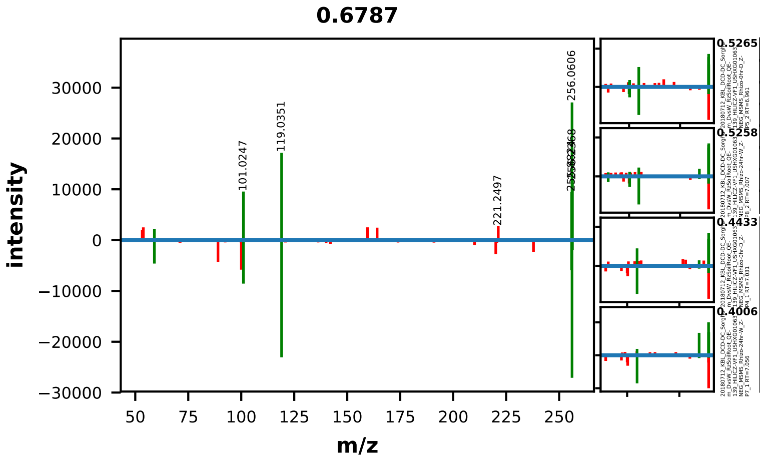


| | EMA Compound Info |
|---|---|
| Name: | N-Acetyl-D-Glucosamine |
| Label: | |
| Formula: | C8H15NO6 |
| Polarity: | negative |
| Monoisotopic Mass: | 221.0899372 |
| Theoretical M/Z: | 256.0593372 |
| Adduct: | |

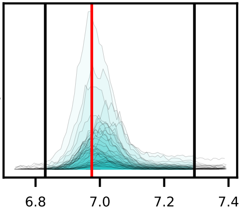

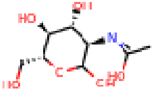

20180712_KBL_DCD-DC_Sorghum_DvsW_RzSoilRoot_QE-139_HILICZ-VF1_USHXG01063_NEG_MSMS_Rhizo-0hr-D_Z-P6_3

0082_N-acetyl-glucosamine_negative_256p0593_6p97
Measured M/Z = 256.0599, 2.1421 ppm difference
Expected Elution of 6.97 minutes, 6.98 min actual

MSMS Scan at 7.008 minutes

Matching M/Zs above 1E-3*max: 59.013, 101.025, 119.035, 255.882, 256.061, 256.237

All Matching M/Zs: 59.013, 101.025, 119.035, 255.882, 256.061, 256.237

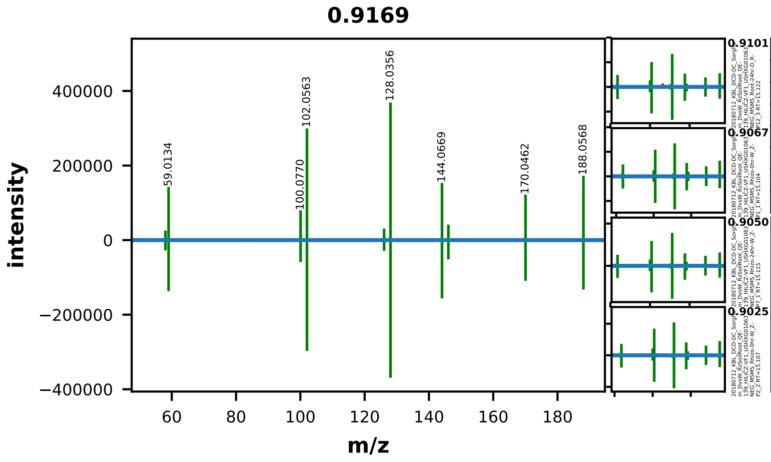


| | EMA Compound Info |
|---|---|
| Name: | N-Acetyl-L-glutamic acid |
| Label: | |
| Formula: | C7H11NO5 |
| Polarity: | negative |
| Monoisotopic Mass: | 189.063722452 |
| Theoretical M/Z: | 188.0564225 |
| Adduct: | |

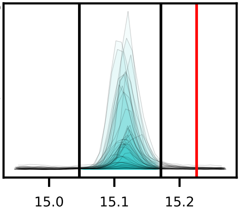


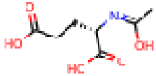

20180712_KBL_DCD-DC_Sorghum_DvsW_RzSoilRoot_QE-139_HILICZ-VF1_USHXG01063_NEG_MSMS_Soil-24hr-D_S-P11_2

0083_N-acetyl-glutamic_acid_negative_188p0564_15p23
Measured M/Z = 188.0567, 1.5158 ppm difference
Expected Elution of 15.23 minutes, 15.12 min actual

MSMS Scan at 15.108 minutes

Matching M/Zs above 1E-3*max: 58.029, 59.013, 100.041, 100.077, 102.056, 126.057, 128.036, 144.067, 146.046, 170.046, 188.057



All Matching M/Zs: 58.029, 59.013, 100.041, 100.077, 102.056, 126.057, 128.036, 144.067, 146.046, 170.046, 188.057

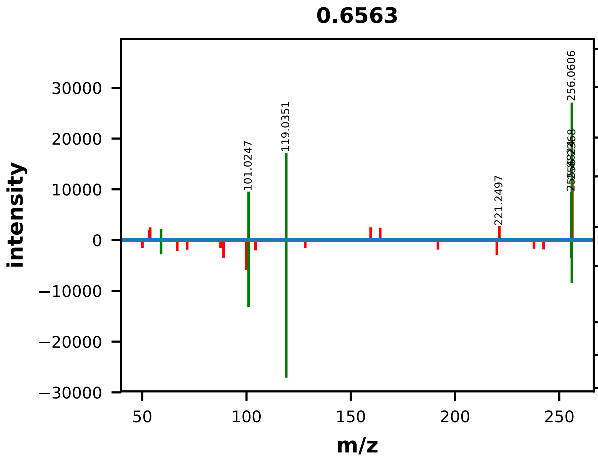


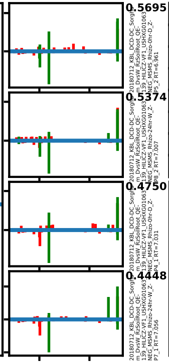


| | EMA Compound Info |
|---|---|
| Name: | N-Acetyl-D-Mannosamine |
| Label: | |
| Formula: | C8H15NO6 |
| Polarity: | negative |
| Monoisotopic Mass: | 221.0899372 |
| Theoretical M/Z: | 256.0593372 |
| Adduct: | |

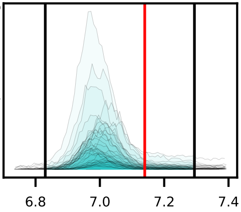


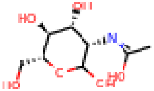

20180712_KBL_DCD-DC_Sorghum_DvsW_RzSoilRoot_QE-139_HILICZ-VF1_USHXG01063_NEG_MSMS_Rhizo-0hr-D_Z-P6_3

0084_N-acetyl-mannosamine_negative_256p0593_7p14
Measured M/Z = 256.0599, 2.1421 ppm difference
Expected Elution of 7.14 minutes, 6.98 min actual

MSMS Scan at 7.008 minutes

Matching M/Zs above 1E-3*max: 59.013, 101.025, 119.035, 255.882, 256.061

All Matching M/Zs: 59.013, 101.025, 119.035, 255.882, 256.061

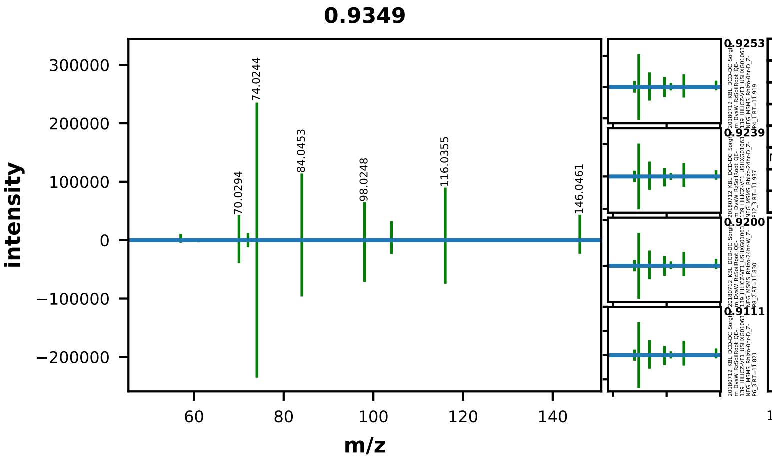


| | EMA Compound Info |
|---|---|
| Name: | 2-acetamido-3-hydroxypropanoic acid |
| Label: | |
| Formula: | C5H9NO4 |
| Polarity: | negative |
| Monoisotopic Mass: | 147.053157768 |
| Theoretical M/Z: | 146.0458578 |
| Adduct: | |

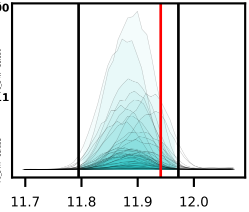


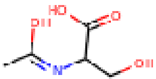


20180712_KBL_DCD-DC_Sorghum_DvsW_RzSoilRoot_QE-139_HILICZ-VF1_USHXG01063_NEG_MSMS_Rhizo-24hr-W_Z-P7_1

0085_N-acetyl-serine_negative_146p0459_11p94
Measured M/Z = 146.0461, 1.3897 ppm difference
Expected Elution of 11.94 minutes, 11.91 min actual

MSMS Scan at 11.843 minutes

Matching M/Zs above 1E-3*max: 57.034, 60.992, 70.029, 72.045, 74.024, 84.045, 98.025, 104.036, 116.036, 146.028, 146.046

All Matching M/Zs: 57.034, 60.992, 70.029, 72.045, 74.024, 84.045, 98.025, 104.036, 116.036, 146.028, 146.046

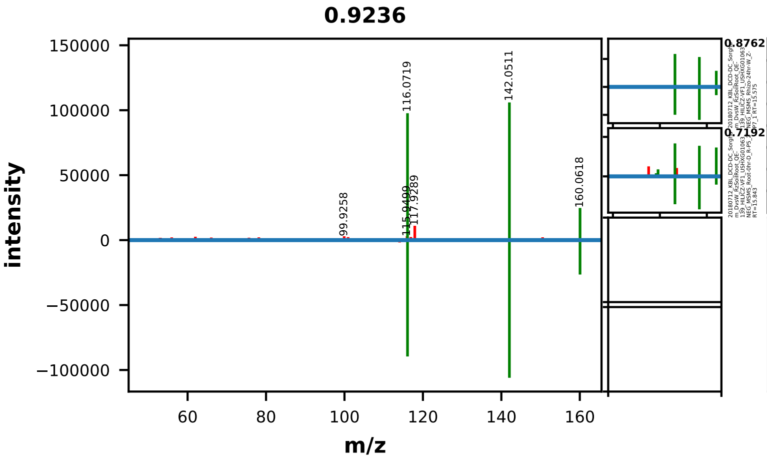


| | EMA Compound Info |
|---|---|
| Name: | N-Methyl-L-glutamic acid |
| Label: | |
| Formula: | C6H11NO4 |
| Polarity: | negative |
| Monoisotopic Mass: | 161.068807832 |
| Theoretical M/Z: | 160.0615078 |
| Adduct: | |

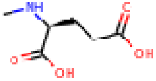

20180712_KBL_DCD-DC_Sorghum_DvsW_RzSoilRoot_QE-139_HILICZ-VF1_USHXG01063_NEG_MSMS_Soil-24hr-D_S-P10_1

0090_N-methyl-glutamic_acid_negative_160p0615_15p21
Measured M/Z = 160.0615, 0.2146 ppm difference
Expected Elution of 15.21 minutes, 15.03 min actual

MSMS Scan at 15.050 minutes

Matching M/Zs above 1E-3*max: 116.072, 142.051, 160.062

All Matching M/Zs: 116.072, 142.051, 160.062

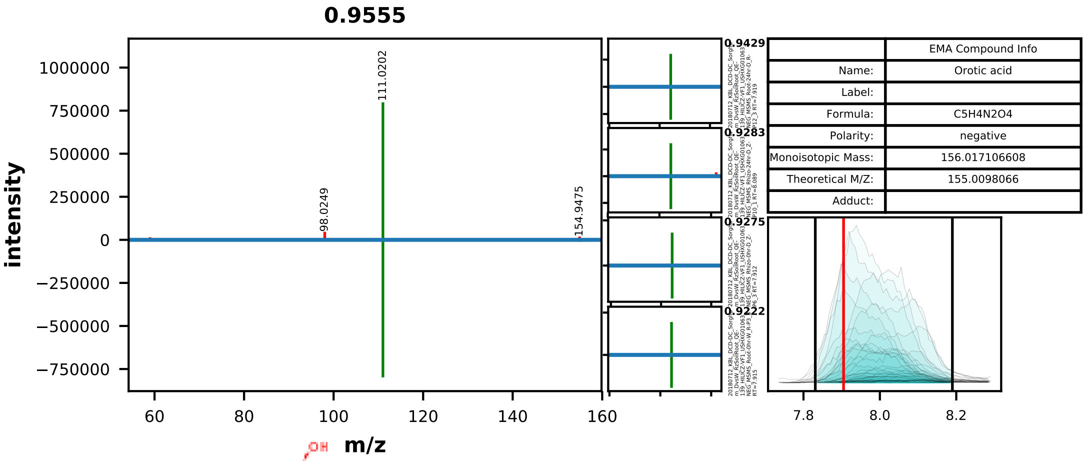


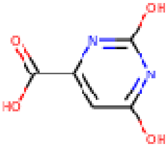


20180712_KBL_DCD-DC_Sorghum_DvsW_RzSoilRoot_QE-139_HILICZ-VF1_USHXG01063_NEG_MSMS_Root-0hr-D_R-P6_3

0093_orotic_acid_negative_155p0098_7p90
Measured M/Z = 155.0100, 1.2545 ppm difference
Expected Elution of 7.90 minutes, 7.90 min actual

MSMS Scan at 8.039 minutes

Matching M/Zs above 1E-3*max: 67.030, 111.020, 155.010

All Matching M/Zs: 67.030, 111.020, 155.010

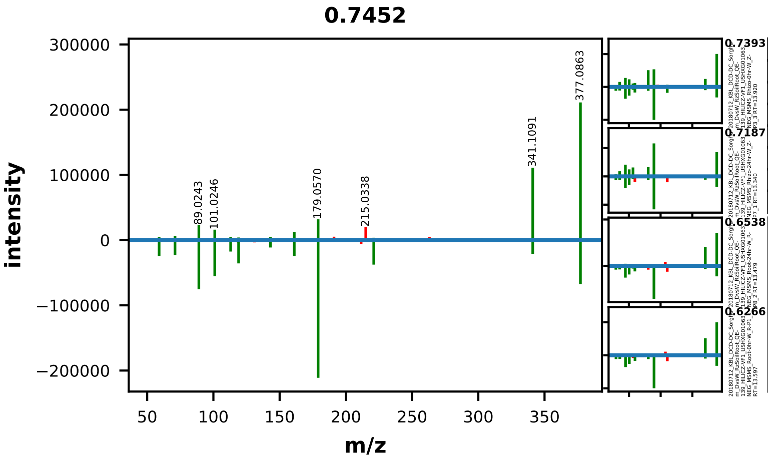


| | EMA Compound Info |
|---|---|
| Name: | Palatinose |
| Label: | |
| Formula: | C12H22O11 |
| Polarity: | negative |
| Monoisotopic Mass: | 342.116211524 |
| Theoretical M/Z: | 377.0856115 |
| Adduct: | |

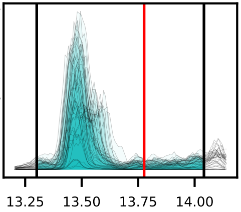


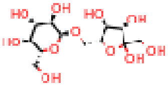


20180712_KBL_DCD-DC_Sorghum_DvsW_RzSoilRoot_QE-139_HILICZ-VF1_USHXG01063_NEG_MSMS_Root-24hr-W_R-P7_1

0094_palatinose_negative_377p0856_13p78
Measured M/Z = 377.0861, 1.3003 ppm difference
Expected Elution of 13.78 minutes, 13.46 min actual

MSMS Scan at 13.979 minutes

Matching M/Zs above 1E-3*max: 59.013, 71.014, 89.024, 101.025, 113.024, 119.035, 143.036, 161.046, 179.057, 221.067, 341.109, 377.086

All Matching M/Zs: 59.013, 71.014, 89.024, 101.025, 113.024, 119.035, 143.036, 161.046, 179.057, 221.067, 341.109, 377.086

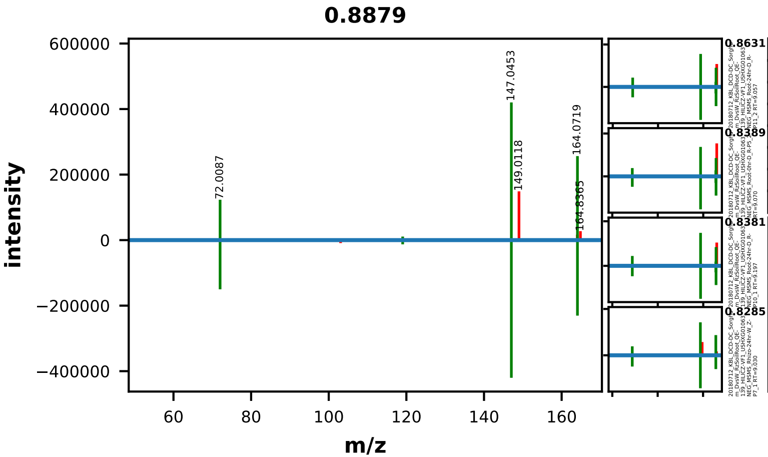


| | EMA Compound Info |
|---|---|
| Name: | L-phenylalanine |
| Label: | |
| Formula: | C9H11NO2 |
| Polarity: | negative |
| Monoisotopic Mass: | 165.078978592 |
| Theoretical M/Z: | 164.0716786 |
| Adduct: | |

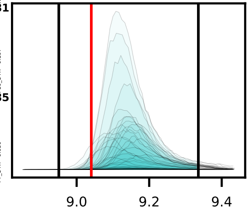


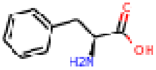


20180712_KBL_DCD-DC_Sorghum_DvsW_RzSoilRoot_QE-139_HILICZ-VF1_USHXG01063_NEG_MSMS_Soil-24hr-D_S-P10_1

0095_phenylalanine_negative_164p0717_9p04
Measured M/Z = 164.0718, 0.8518 ppm difference
Expected Elution of 9.04 minutes, 9.13 min actual

MSMS Scan at 9.039 minutes

Matching M/Zs above 1E-3*max: 72.009, 119.050, 147.045, 164.072

All Matching M/Zs: 72.009, 119.050, 147.045, 164.072

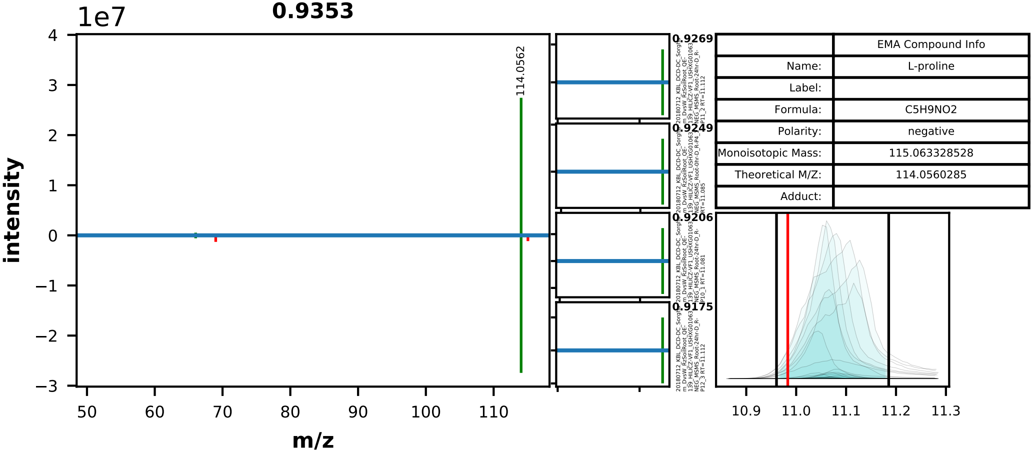


| | EMA Compound Info |
|---|---|
| Name: | L-proline |
| Label: | |
| Formula: | C5H9NO2 |
| Polarity: | negative |
| Monoisotopic Mass: | 115.063328528 |
| Theoretical M/Z: | 114.0560285 |
| Adduct: | |

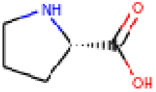


20180712_KBL_DCD-DC_Sorghum_DvsW_RzSoilRoot_QE-139_HILICZ-VF1_USHXG01063_NEG_MSMS_Root-0hr-D_R-P5_2

0096_proline_negative_114p0560_10p98
Measured M/Z = 114.0562, 1.3580 ppm difference
Expected Elution of 10.98 minutes, 11.13 min actual

MSMS Scan at 11.100 minutes

Matching M/Zs above 1E-3*max: 66.034, 86.025, 114.056

All Matching M/Zs: 66.034, 86.025, 114.056

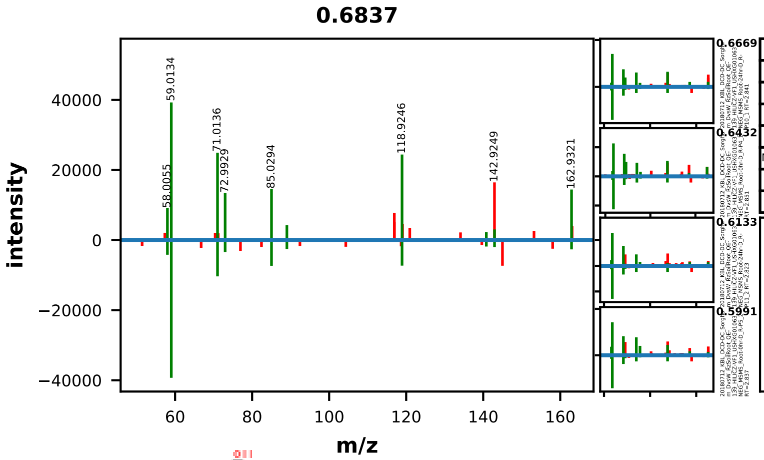


| | EMA Compound Info |
|---|---|
| Name: | L-Rhamnose |
| Label: | |
| Formula: | C6H12O5 |
| Polarity: | negative |
| Monoisotopic Mass: | 164.068473484 |
| Theoretical M/Z: | 163.0611735 |
| Adduct: | |

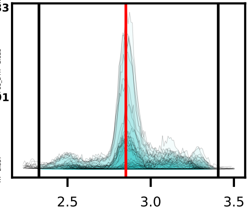


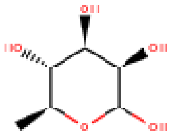

20180712_KBL_DCD-DC_Sorghum_DvsW_RzSoilRoot_QE-139_HILICZ-VF1_USHXG01063_NEG_MSMS_Rhizo-0hr-D_Z-P6_3

0099_rhamnose_negative_163p0612_2p85
Measured M/Z = 163.0614, 1.5456 ppm difference
Expected Elution of 2.85 minutes, 2.86 min actual

MSMS Scan at 2.852 minutes

Matching M/Zs above 1E-3*max: 58.006, 59.013, 71.014, 72.993, 85.029, 89.025, 118.925, 140.811, 142.937, 162.932

All Matching M/Zs: 58.006, 59.013, 71.014, 72.993, 85.029, 89.025, 118.925, 140.811, 142.937, 162.932

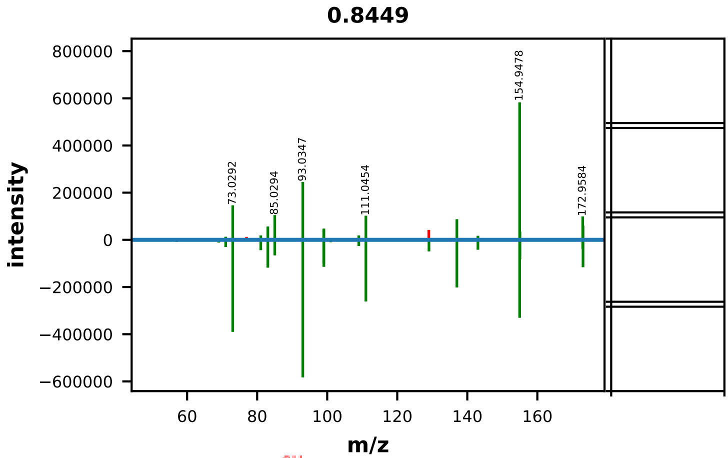


| | EMA Compound Info |
|---|---|
| Name: | shikimic acid |
| Label: | |
| Formula: | C7H10O5 |
| Polarity: | negative |
| Monoisotopic Mass: | 174.05282342 |
| Theoretical M/Z: | 173.0455234 |
| Adduct: | |

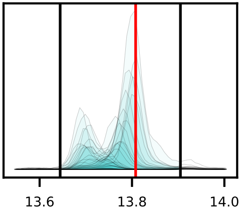


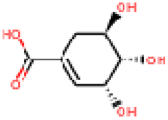


20180712_KBL_DCD-DC_Sorghum_DvsW_RzSoilRoot_QE-139_HILICZ-VF1_USHXG01063_NEG_MSMS_Rhizo-24hr-D_Z-P12_3

0101_shikimic_acid_negative_173p0455_13p81
Measured M/Z = 173.0456, 0.4612 ppm difference
Expected Elution of 13.81 minutes, 13.75 min actual

MSMS Scan at 13.738 minutes

Matching M/Zs above 1E-3*max: 55.018, 57.034, 69.034, 71.014, 73.029, 81.034, 83.050, 85.029, 93.035, 99.009, 99.045, 101.025, 109.030, 111.045, 129.056, 137.025, 143.036, 154.948, 155.035, 172.958, 173.046



All Matching M/Zs: 55.018, 57.034, 69.034, 71.014, 73.029, 81.034, 83.050, 85.029, 93.035, 99.009, 99.045, 101.025, 109.030, 111.045, 129.056, 137.025, 143.036, 154.948, 155.035, 172.958, 173.046

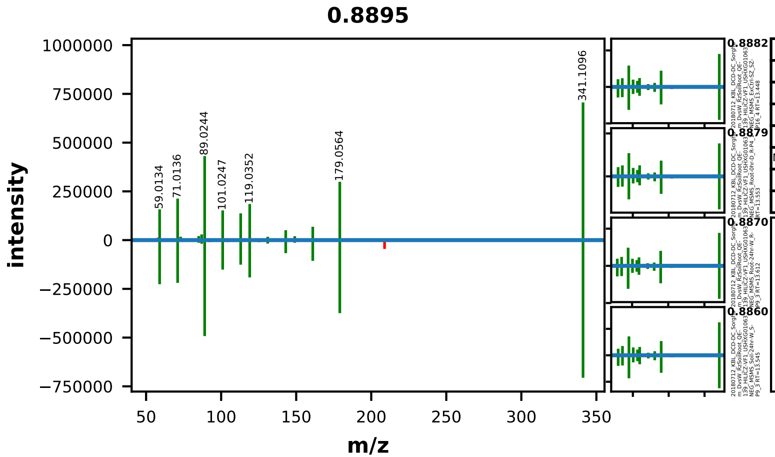


| | EMA Compound Info |
|---|---|
| Name: | sucrose |
| Label: | |
| Formula: | C12H22O11 |
| Polarity: | negative |
| Monoisotopic Mass: | 342.116211524 |
| Theoretical M/Z: | 341.1089115 |
| Adduct: | |

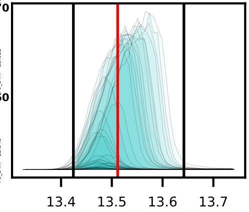


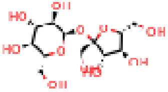

20180712_KBL_DCD-DC_Sorghum_DvsW_RzSoilRoot_QE-139_HILICZ-VF1_USHXG01063_NEG_MSMS_Soil-24hr-D_S-P11_2

0103_sucrose_negative_341p1089_13p51
Measured M/Z = 341.1096, 2.1185 ppm difference
Expected Elution of 13.51 minutes, 13.47 min actual

MSMS Scan at 13.427 minutes

Matching M/Zs above 1E-3*max: 59.013, 71.014, 72.993, 85.029, 87.009, 89.024, 101.025, 113.025, 119.035, 131.035, 143.035, 149.046, 161.046, 179.056, 341.110

All Matching M/Zs: 59.013, 71.014, 72.993, 85.029, 87.009, 89.024, 101.025, 113.025, 119.035, 131.035, 143.035, 149.046, 161.046, 179.056, 341.110

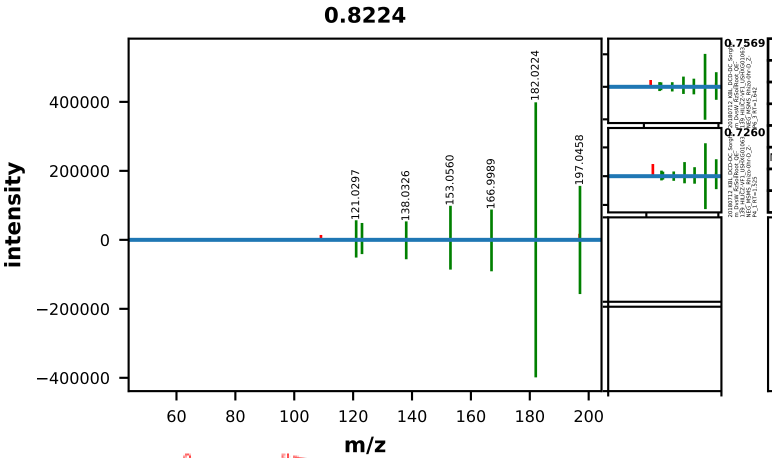


| | EMA Compound Info |
|---|---|
| Name: | SYRINGIC ACID |
| Label: | |
| Formula: | C9H10O5 |
| Polarity: | negative |
| Monoisotopic Mass: | 198.05282342 |
| Theoretical M/Z: | 197.0455234 |
| Adduct: | |

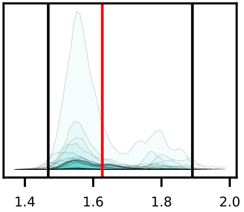


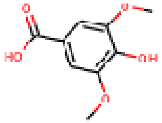

20180712_KBL_DCD-DC_Sorghum_DvsW_RzSoilRoot_QE-139_HILICZ-VF1_USHXG01063_NEG_MSMS_Root-24hr-D_R-P10_1

0104_syringic_acid_negative_197p0455_1p63
Measured M/Z = 197.0456, 0.3110 ppm difference
Expected Elution of 1.63 minutes, 1.55 min actual

MSMS Scan at 1.559 minutes

Matching M/Zs above 1E-3*max: 95.014, 121.030, 123.009, 137.025, 138.033, 153.056, 166.999, 178.891, 182.022, 197.046



All Matching M/Zs: 95.014, 121.030, 123.009, 137.025, 138.033, 153.056, 166.999, 178.891, 182.022, 197.046

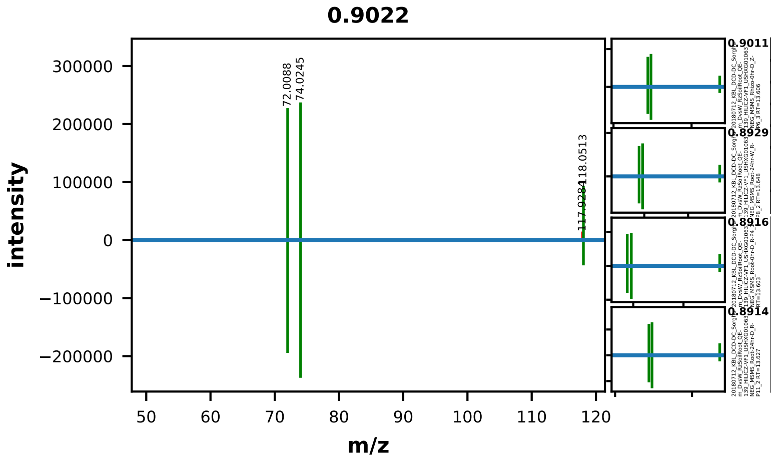


| | EMA Compound Info |
|---|---|
| Name: | L-threonine |
| Label: | |
| Formula: | C4H9NO3 |
| Polarity: | negative |
| Monoisotopic Mass: | 119.058243148 |
| Theoretical M/Z: | 118.0509431 |
| Adduct: | |

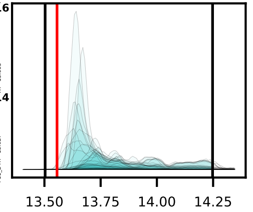


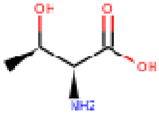


20180712_KBL_DCD-DC_Sorghum_DvsW_RzSoilRoot_QE-139_HILICZ-VF1_USHXG01063_NEG_MSMS_Root-0hr-W_R-P1_1

0107_threonine_negative_118p0509_13p56
Measured M/Z = 118.0511, 1.5081 ppm difference
Expected Elution of 13.56 minutes, 13.74 min actual

MSMS Scan at 13.662 minutes

Matching M/Zs above 1E-3*max: 72.009, 74.024, 92.055, 118.051

All Matching M/Zs: 72.009, 74.024, 92.055, 118.051

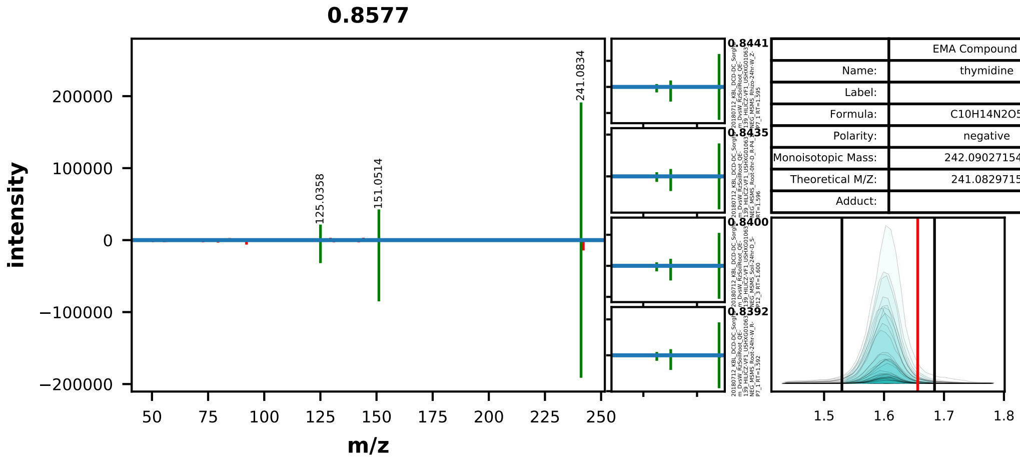


| | EMA Compound Info |
|---|---|
| Name: | thymidine |
| Label: | |
| Formula: | C10H14N2O5 |
| Polarity: | negative |
| Monoisotopic Mass: | 242.090271548 |
| Theoretical M/Z: | 241.0829715 |
| Adduct: | |

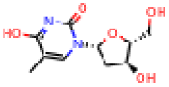


20180712_KBL_DCD-DC_Sorghum_DvsW_RzSoilRoot_QE-139_HILICZ-VF1_USHXG01063_NEG_MSMS_Root-24hr-W_R-P9_3

0108_thymidine_negative_241p0830_1p66
Measured M/Z = 241.0833, 1.5667 ppm difference
Expected Elution of 1.66 minutes, 1.61 min actual

MSMS Scan at 1.593 minutes

Matching M/Zs above 1E-3*max: 125.036, 151.051, 241.083

All Matching M/Zs: 125.036, 151.051, 241.083

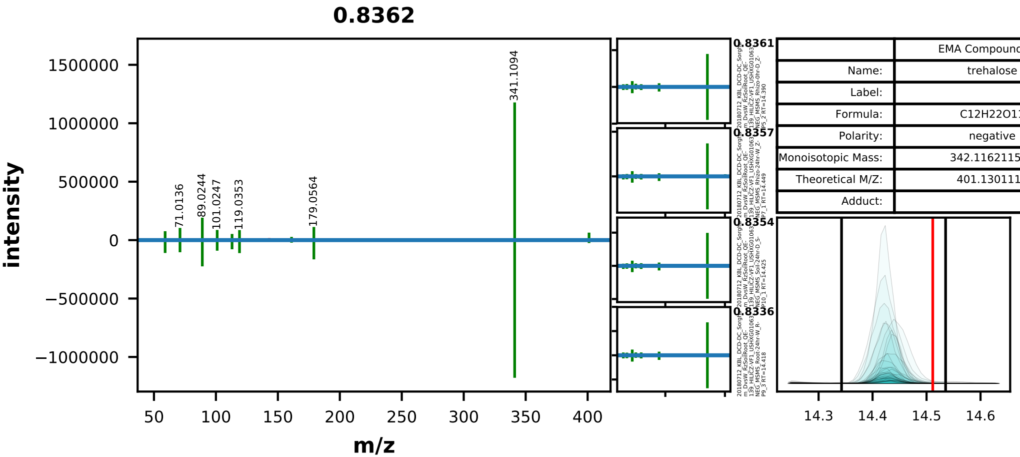


| | EMA Compound Info |
|---|---|
| Name: | trehalose |
| Label: | |
| Formula: | C12H22O11 |
| Polarity: | negative |
| Monoisotopic Mass: | 342.116211524 |
| Theoretical M/Z: | 401.1301115 |
| Adduct: | |

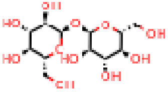

20180712_KBL_DCD-DC_Sorghum_DvsW_RzSoilRoot_QE-139_HILICZ-VF1_USHXG01063_NEG_MSMS_Soil-0hr-W_S-P3_3

0111_trehalose_negative_401p1301_14p51
Measured M/Z = 401.1306, 1.1163 ppm difference
Expected Elution of 14.51 minutes, 14.43 min actual

MSMS Scan at 14.435 minutes

Matching M/Zs above 1E-3*max: 59.013, 71.014, 89.024, 101.025, 113.025, 119.035, 161.046, 179.056, 341.109, 401.130



All Matching M/Zs: 59.013, 71.014, 89.024, 101.025, 113.025, 119.035, 161.046, 179.056, 341.109, 401.130

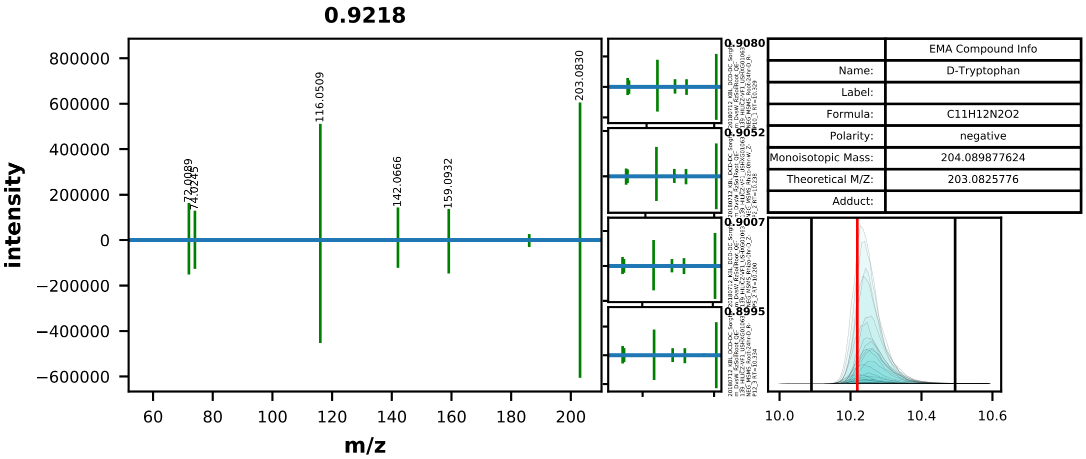


| | EMA Compound Info |
|---|---|
| Name: | D-Tryptophan |
| Label: | |
| Formula: | C11H12N2O2 |
| Polarity: | negative |
| Monoisotopic Mass: | 204.089877624 |
| Theoretical M/Z: | 203.0825776 |
| Adduct: | |

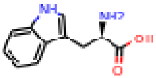


20180712_KBL_DCD-DC_Sorghum_DvsW_RzSoilRoot_QE-139_HILICZ-VF1_USHXG01063_NEG_MSMS_Root-24hr-D_R-P11_2

0112_tryptophan_negative_203p0826_10p22
Measured M/Z = 203.0829, 1.6922 ppm difference
Expected Elution of 10.22 minutes, 10.23 min actual

MSMS Scan at 10.334 minutes

Matching M/Zs above 1E-3*max: 72.009, 74.024, 116.051, 130.066, 142.067, 159.093, 161.048, 186.056, 203.083

All Matching M/Zs: 72.009, 74.024, 116.051, 130.066, 142.067, 159.093, 161.048, 186.056, 203.083

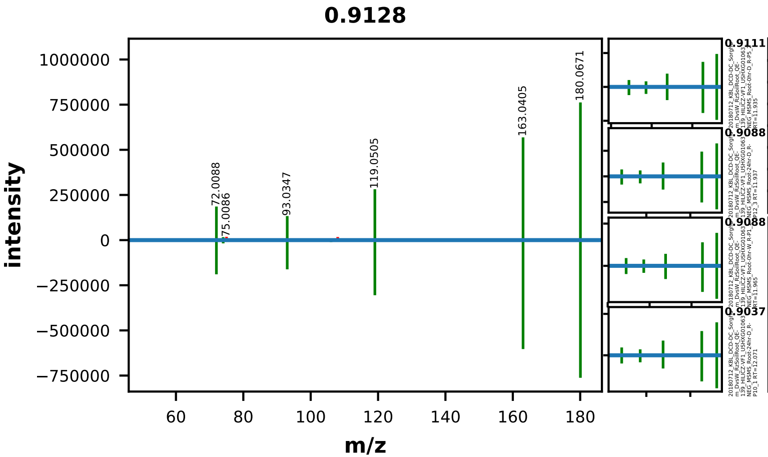


| | EMA Compound Info |
|---|---|
| Name: | L-tyrosine |
| Label: | |
| Formula: | C9H11NO3 |
| Polarity: | negative |
| Monoisotopic Mass: | 181.073893212 |
| Theoretical M/Z: | 180.0665932 |
| Adduct: | |

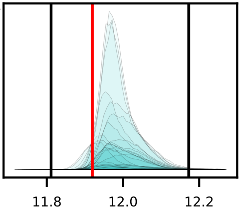


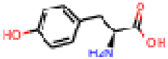


20180712_KBL_DCD-DC_Sorghum_DvsW_RzSoilRoot_QE-139_HILICZ-VF1_USHXG01063_NEG_MSMS_Rhizo-24hr-D_Z-P10_1

0113_tyrosine_negative_180p0666_11p92
Measured M/Z = 180.0669, 1.4927 ppm difference
Expected Elution of 11.92 minutes, 11.94 min actual

MSMS Scan at 11.905 minutes

Matching M/Zs above 1E-3*max: 72.009, 74.025, 93.035, 106.043, 107.051, 119.051, 134.062, 163.040, 180.067



All Matching M/Zs: 72.009, 74.025, 93.035, 106.043, 107.051, 119.051, 134.062, 163.040, 180.067

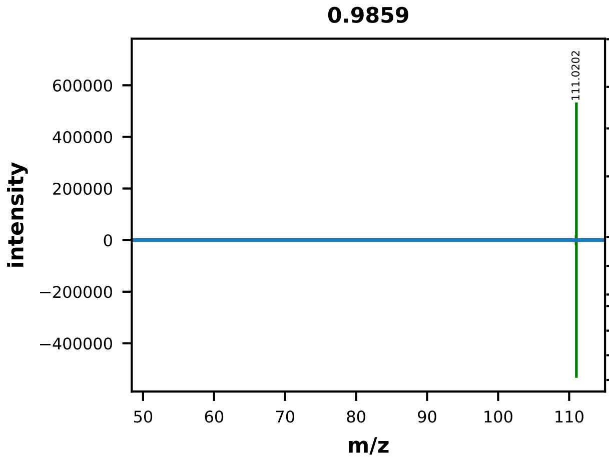


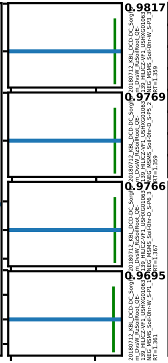


| | EMA Compound Info |
|---|---|
| Name: | uracil |
| Label: | |
| Formula: | C4H4N2O2 |
| Polarity: | negative |
| Monoisotopic Mass: | 112.027277368 |
| Theoretical M/Z: | 111.0199774 |
| Adduct: | |

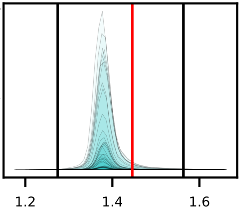


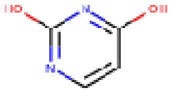


20180712_KBL_DCD-DC_Sorghum_DvsW_RzSoilRoot_QE-139_HILICZ-VF1_USHXG01063_NEG_MSMS_Rhizo-24hr-D_Z-P10_1

0114_uracil_negative_111p0200_1p45
Measured M/Z = 111.0202, 1.8295 ppm difference
Expected Elution of 1.45 minutes, 1.38 min actual

MSMS Scan at 1.457 minutes

Matching M/Zs above 1E-3*max: 110.976, 111.020

All Matching M/Zs: 110.976, 111.020

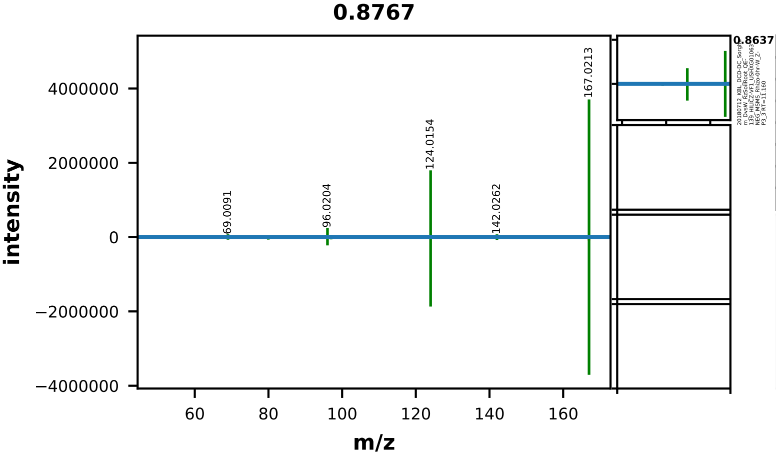


| | EMA Compound Info |
|---|---|
| Name: | uric acid |
| Label: | |
| Formula: | C5H4N4O3 |
| Polarity: | negative |
| Monoisotopic Mass: | 168.028339988 |
| Theoretical M/Z: | 167.02104 |
| Adduct: | |

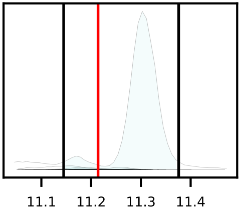


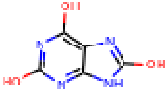


20180712_KBL_DCD-DC_Sorghum_DvsW_RzSoilRoot_QE-139_HILICZ-VF1_USHXG01063_NEG_MSMS_Rhizo-0hr-D_Z-P4_1

0115_uric_acid_negative_167p0210_11p21
Measured M/Z = 167.0212, 0.6817 ppm difference
Expected Elution of 11.21 minutes, 11.30 min actual

MSMS Scan at 11.268 minutes

Matching M/Zs above 1E-3*max: 69.009, 79.957, 96.020, 97.004, 122.023, 123.032, 124.015, 142.026, 167.021

All Matching M/Zs: 69.009, 79.957, 96.020, 97.004, 122.023, 123.032, 124.015, 142.026, 167.021

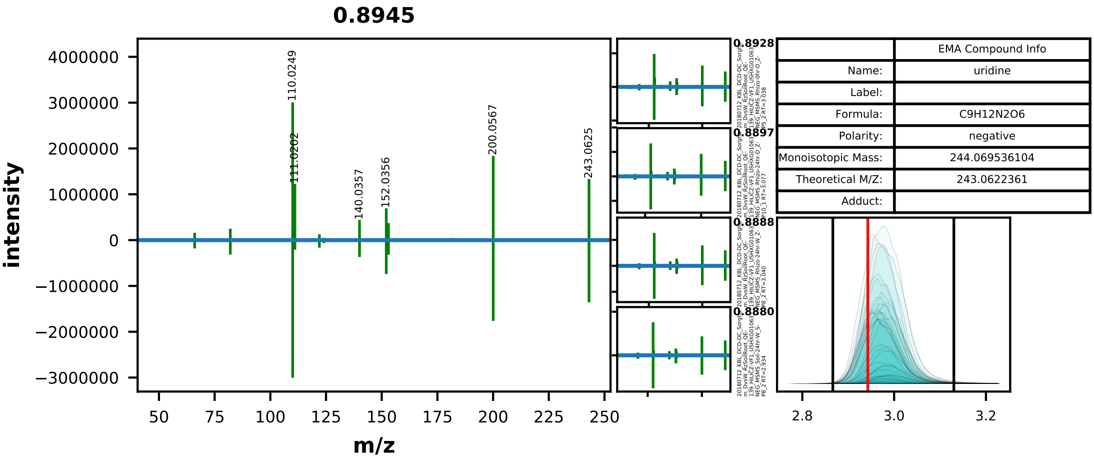


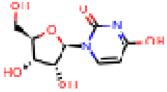

20180712_KBL_DCD-DC_Sorghum_DvsW_RzSoilRoot_QE-139_HILICZ-VF1_USHXG01063_NEG_MSMS_Root-24hr-W_R-P8_2

0116_uridine_negative_243p0622_2p94
Measured M/Z = 243.0624, 0.5744 ppm difference
Expected Elution of 2.94 minutes, 2.96 min actual

MSMS Scan at 3.025 minutes

Matching M/Zs above 1E-3*max: 66.034, 82.030, 96.046, 110.025, 111.020, 122.025, 124.040, 140.036, 152.036, 153.031, 200.057, 243.063



All Matching M/Zs: 66.034, 82.030, 96.046, 110.025, 111.020, 122.025, 124.040, 140.036, 152.036, 153.031, 200.057, 243.063

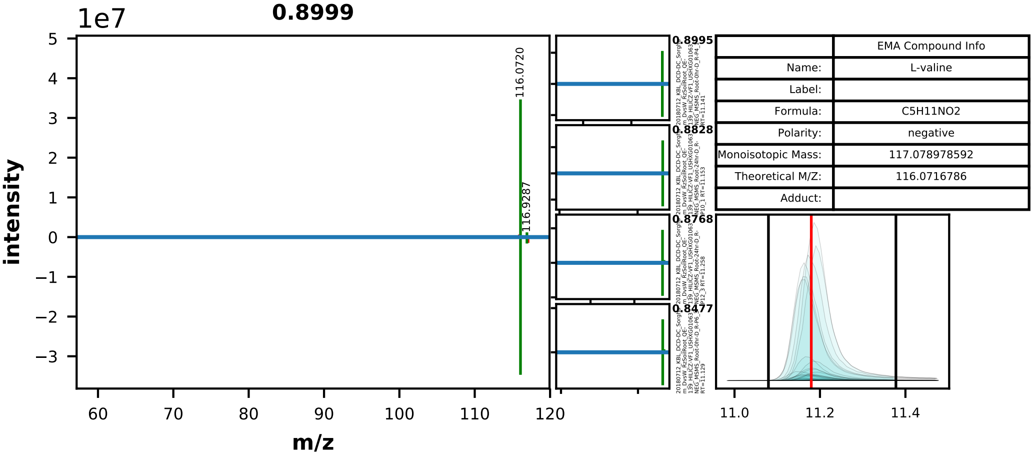


| | EMA Compound Info |
|---|---|
| Name: | L-valine |
| Label: | |
| Formula: | C5H11NO2 |
| Polarity: | negative |
| Monoisotopic Mass: | 117.078978592 |
| Theoretical M/Z: | 116.0716786 |
| Adduct: | |

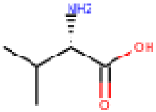


20180712_KBL_DCD-DC_Sorghum_DvsW_RzSoilRoot_QE-139_HILICZ-VF1_USHXG01063_NEG_MSMS_Root-24hr-D_R-P11_2

0118_valine_negative_116p0717_11p18
Measured M/Z = 116.0719, 2.1534 ppm difference
Expected Elution of 11.18 minutes, 11.19 min actual

MSMS Scan at 11.161 minutes

Matching M/Zs above 1E-3*max: 99.926, 116.072, 116.929

All Matching M/Zs: 99.926, 116.072, 116.929

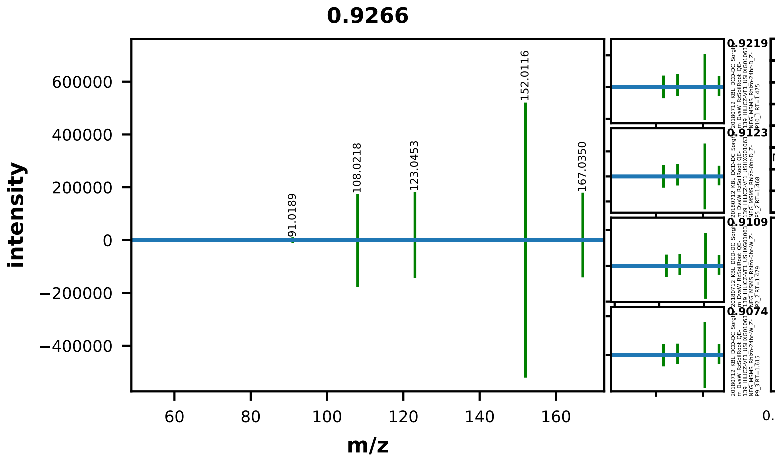


| | EMA Compound Info |
|---|---|
| Name: | Vanillic acid |
| Label: | |
| Formula: | C8H8O4 |
| Polarity: | negative |
| Monoisotopic Mass: | 168.042258736 |
| Theoretical M/Z: | 167.0349587 |
| Adduct: | |

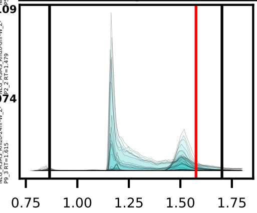


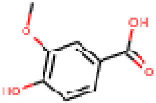


20180712_KBL_DCD-DC_Sorghum_DvsW_RzSoilRoot_QE-139_HILICZ-VF1_USHXG01063_NEG_MSMS_Rhizo-0hr-D_Z-P4_1

0119_vanillic_acid_negative_167p0350_1p58
Measured M/Z = 167.0349, 0.1197 ppm difference
Expected Elution of 1.58 minutes, 1.51 min actual

MSMS Scan at 1.474 minutes

Matching M/Zs above 1E-3*max: 91.019, 108.022, 111.010, 123.045, 152.012, 167.035

All Matching M/Zs: 91.019, 108.022, 111.010, 123.045, 152.012, 167.035

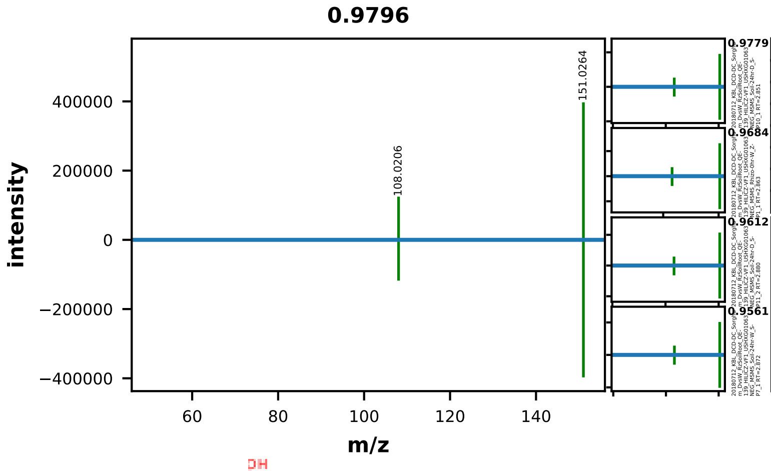


| | EMA Compound Info |
|---|---|
| Name: | xanthine |
| Label: | |
| Formula: | C5H4N4O2 |
| Polarity: | negative |
| Monoisotopic Mass: | 152.033425368 |
| Theoretical M/Z: | 151.0261254 |
| Adduct: | |

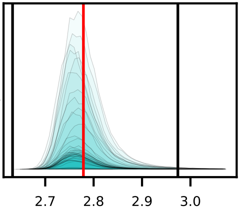


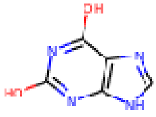


20180712_KBL_DCD-DC_Sorghum_DvsW_RzSoilRoot_QE-139_HILICZ-VF1_USHXG01063_NEG_MSMS_Soil-24hr-D_S-P12_3

0120_xanthine_negative_151p0261_2p78
Measured M/Z = 151.0265, 2.3188 ppm difference
Expected Elution of  2.78 minutes,  2.75 min actual

MSMS Scan at 2.722 minutes

Matching M/Zs above 1E-3*max: 108.021, 126.031, 151.026

All Matching M/Zs: 108.021, 126.031, 151.026

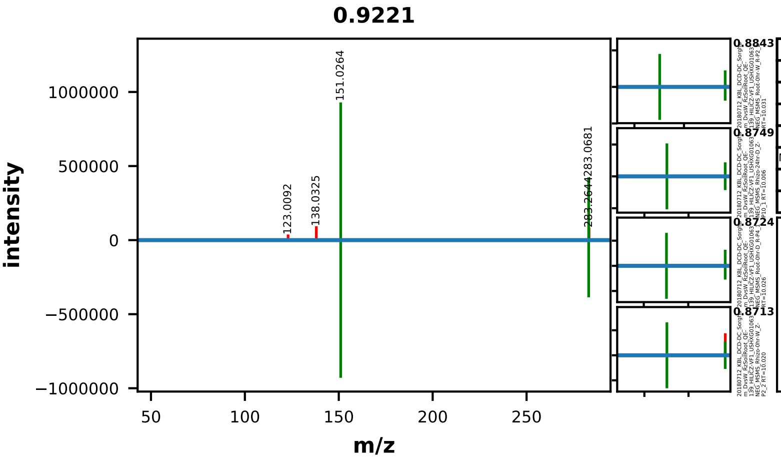


| | EMA Compound Info |
|---|---|
| Name: | xanthosine |
| Label: | |
| Formula: | C10H12N4O6 |
| Polarity: | negative |
| Monoisotopic Mass: | 284.075684104 |
| Theoretical M/Z: | 283.0683841 |
| Adduct: | |

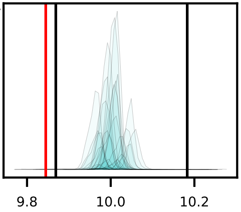


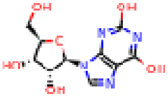


20180712_KBL_DCD-DC_Sorghum_DvsW_RzSoilRoot_QE-139_HILICZ-VF1_USHXG01063_NEG_MSMS_Root-0hr-D_R-P6_3

0121_xanthosine_negative_283p0684_9p84
Measured M/Z = 283.0687, 1.2829 ppm difference
Expected Elution of 9.84 minutes, 9.99 min actual

MSMS Scan at 10.002 minutes

Matching M/Zs above 1E-3*max: 108.021, 151.026, 283.068

All Matching M/Zs: 108.021, 151.026, 283.068
