## Supplementary figures and images for "Drought shifts sorghum root metabolite and microbiome profiles and enriches the stress response factor pipecolic acid"

### Supplemental Figure 2

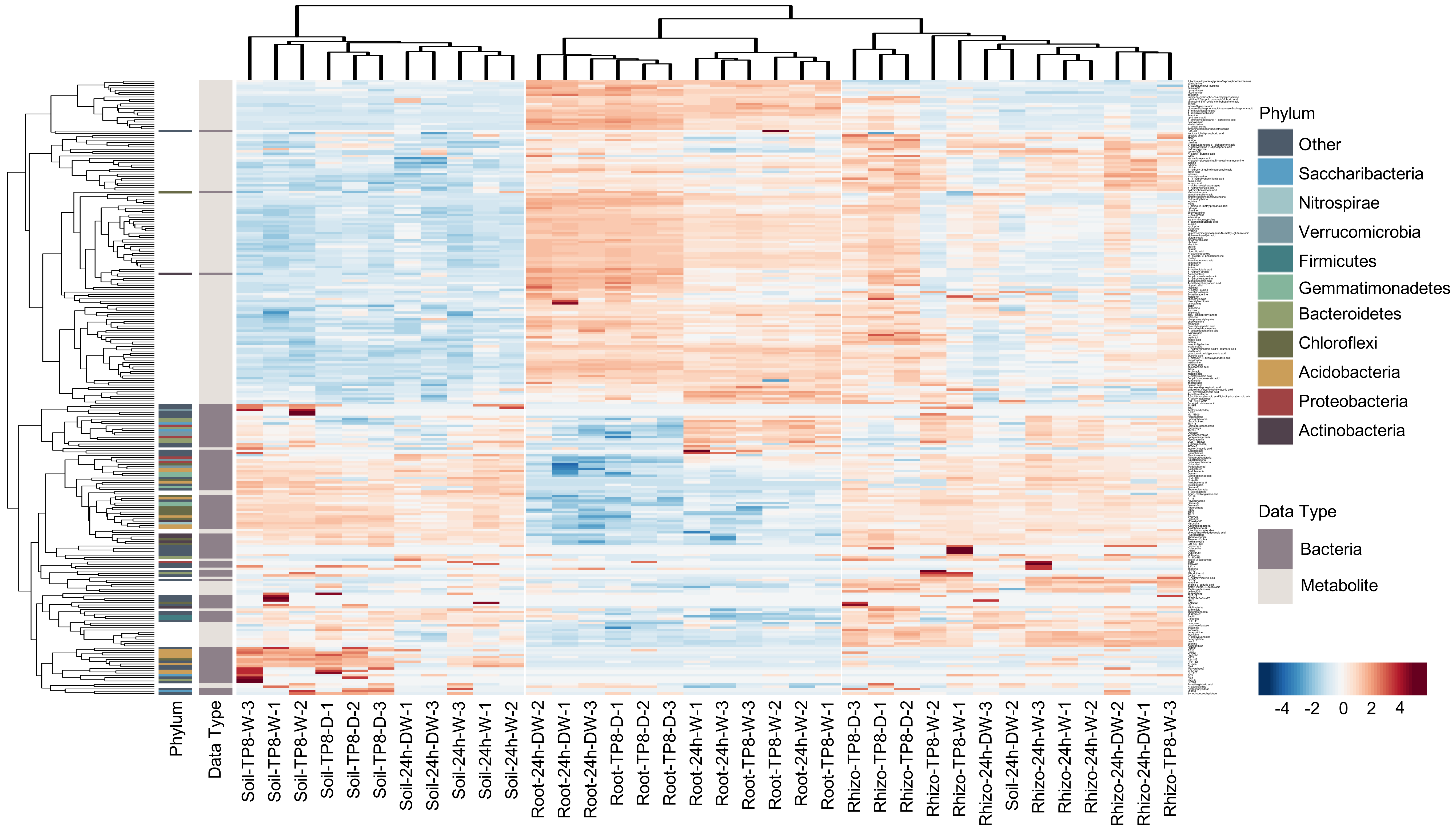

### Supplemental Figure 3

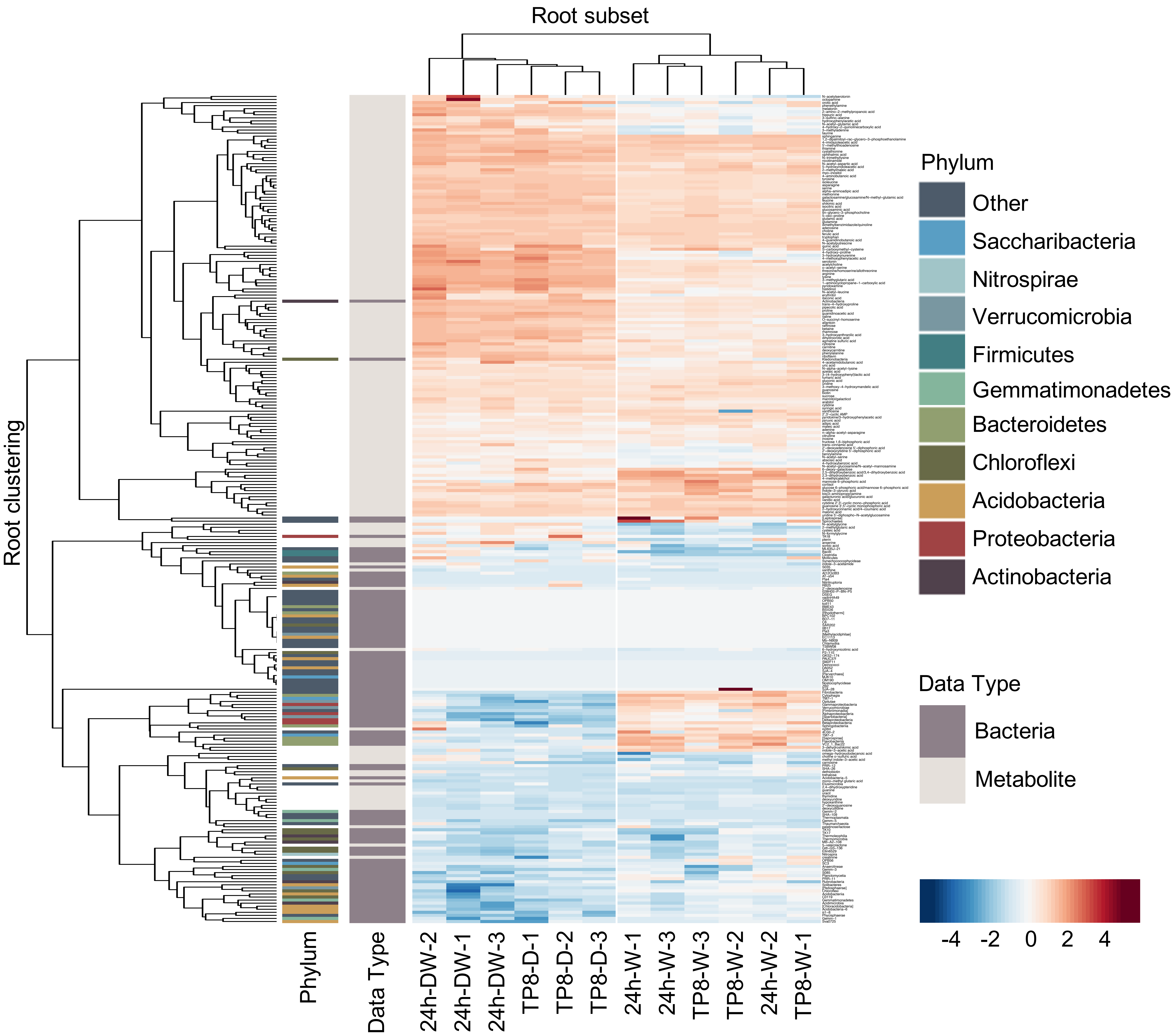
